# Two Distinct Binding Modes Govern High-Affinity Ligand Interactions with Amyloid Fibrils

**DOI:** 10.64898/2026.02.15.705980

**Authors:** Timothy S. Chisholm

## Abstract

Fibrillar protein aggregates are a defining feature of neurodegenerative diseases and are attractive biomarkers and therapeutic targets. However, rational ligand design is limited by a poor mechanistic understanding of fibril binding. This work demonstrates that high-affinity binding to amyloid fibrils occurs via two topologically distinct binding modes, informing design changes that enhance ligand binding. Mathematical models were outlined that demonstrate these binding modes can be distinguished using diagnostic features from standard binding assays. Reanalysis of published binding data indicates that these binding modes are likely widespread amongst common ligand scaffolds. Guided by these binding modes, new ligands were designed with improved binding affinities and distinct fluorescence responses. Together, these findings support the presence of two prevalent binding modes and establish new design principles for enhancing interactions between ligands and amyloid fibrils.

## Introduction

Many neurodegenerative diseases, such as Alzheimer’s disease (AD) and Parkinson’s disease (PD), are characterised by fibrillar protein aggregates that form in the brain.^1–3^ These aggregates, commonly referred to as amyloid fibrils, are composed primarily of amyloid-β and tau in AD, and α-synuclein in PD. Extensive research efforts have gone toward developing high-affinity small molecule ligands targeting amyloid fibrils for diagnostic and therapeutic applications.^4–7^

However, the precise binding mode of ligands to amyloid fibrils remains obscure. Ligands bind multiple types of sites on amyloid fibril,^4,8–10^ or bind via multiple binding modes.^11,12^ Ligands appear to bind to β-sheet-rich grooves along the fibril surface with regular orientation and spacing, aligned either parallel or perpendicular to the length of the fibril.^13–17^

Recent cryo-EM structures of fibrils with bound ligands suggest two primary binding modes may occur (Figure 1).^18–22^ A *stacked* binding mode, also referred to as horizontal or diagonal binding, is defined by bound ligands that are in close contact and with one ligand bound per protein monomer (Figure 1a). A *linear* binding mode, also referred to as vertical binding, is defined by ligands aligned along the fibril axis spanning multiple protein monomers (Figure 1b). A *linear-new site* binding mode, where binding of one ligand creates a new site underpinned by ligand-ligand interactions, has also been suggested (Figure 1c).^22^ Notably, these structures were obtained using concentrations of ligand (50-500 µM) at least an order of magnitude greater than their dissociation constants.

**Figure 1.**
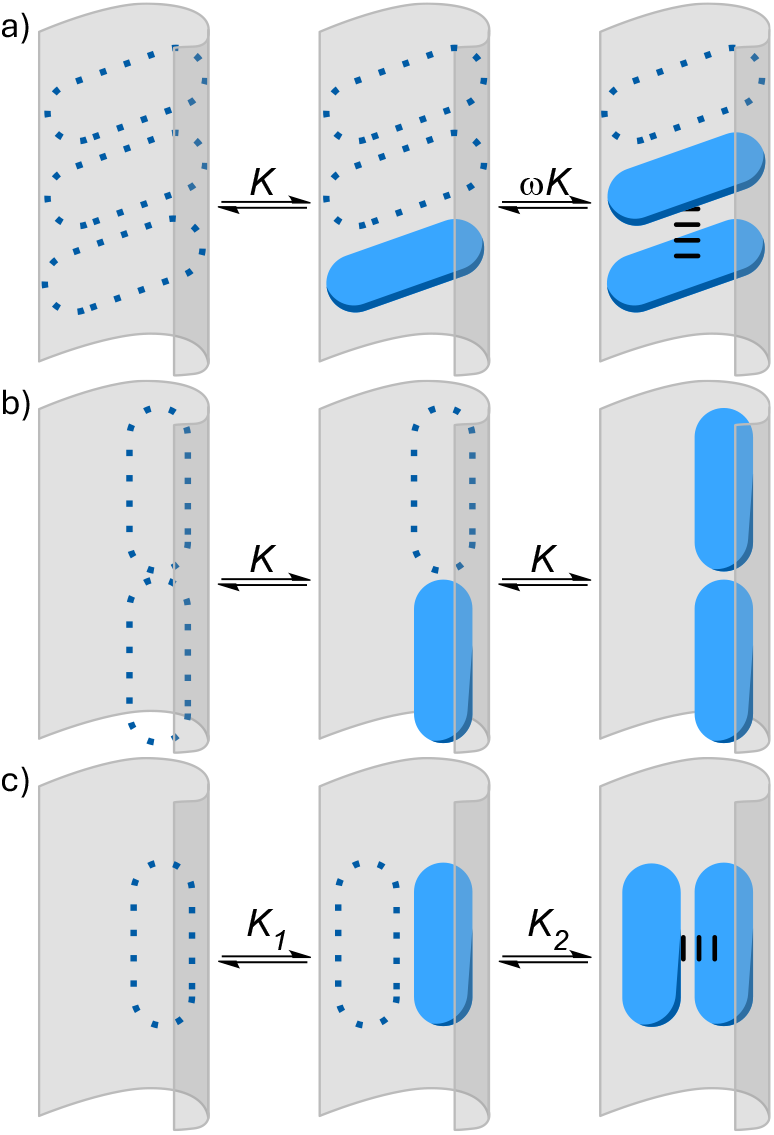
(a) Stacked binding mode. (b) Linear binding mode. (c) Linear-new site binding mode, where a bound ligand generates a new binding site. *K* is the binding constant, and ω is the cooperativity parameter.

These reported bound poses do not necessarily represent the thermodynamic minima present for ligands binding with nanomolar or low-micromolar dissociation constants. Further evidence is therefore needed to establish which binding modes dominate at meaningful ligand concentrations. Standard binding assays may provide indirect evidence, as different binding modes could produce binding isotherms with potentially diagnostic features. Such data are typically analysed using models based on the Langmuir isotherm, or variations thereof. However, models used for canonical protein-ligand interactions are not directly appropriate for fibril-ligand interactions where ligands bind to a continuum of possible binding sites along a hydrophobic groove rather than at discrete sites.

A binding model is therefore required that can account for this continuum of adjacent sites, and that can model stacked and linear binding. A stacked binding model must account for cooperative binding, and the different probability for ligands to bind adjacently and non-adjacently. A linear binding model must account for a single ligand binding to multiple protein monomers, and that the number of available binding sites depends on the size and spatial arrangement of bound ligands. Previous work has provided a model that can address cooperative binding in a fibril system, but that could not account for linear binding, nor multiple types of ligands and binding sites.^23^ Here, such a model is implemented using the McGhee and von Hippel procedure,^24^ providing analytical evidence supporting stacked and linear binding modes at nanomolar binding regimes. Knowledge of these binding modes is then used to prepare ligands with improved binding affinities.

### Derivation of a Binding Model

First, a set of equations are derived to model the equilibrium binding of multiple types of ligands to multiple types of sites on a 1D polymer, representing an amyloid fibril, while accounting for both ligand size and cooperativity. In a typical binding equilibrium, the concentration of binding sites is conserved according to the equation

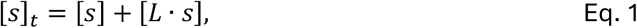

where [*s*]_*t*_ is the total concentration of binding sites, [*s*]is the concentration of free binding sites, and [*L* ∙ *s*]is the concentration of bound ligand. This equation does not hold on a 1D polymer, where the concentration of available binding sites also depends on the size and spatial distribution of bound ligands (Figure 2a). This challenge can be addressed following a procedure described by McGhee and von Hippel,^24^ and later expanded by Villaluenga *et al*,^25^ to count the number of possible binding sites on a 1D polymer by describing ligand binding using conditional probabilities.

**Figure 2.**
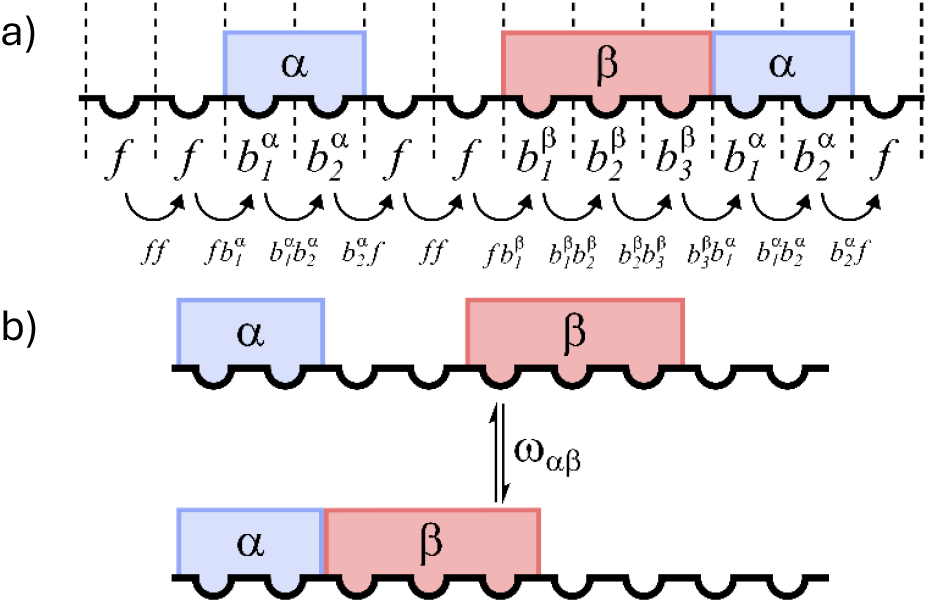
(a) Schematic illustrating the binding of two ligands (α, β) to lattice residues, the type of lattice residue (*f* or 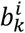), and the conditional probabilities. (b) Definition of the cooperativity parameter ω used in this model.

Consider a linear polymer composed of *N* individual lattice residues, analogous to an amyloid fibril composed of *N* protein monomers. A ligand of type *i* covers *m*_*i*_ lattice residues when bound and is described as consisting of *m*_*i*_ segments labelled 1, 2, …, *m*_*i*_ from left to right. Across *T* total types of ligands there are therefore 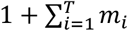possible types of residues in the linear polymer. Free residues are denoted *f*, and residues bound to a ligand are denoted by 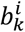for segment *k* of a ligand of type *i*.

Conditional probabilities are then defined as a sequence of two types of lattice residues. There are four relevant conditional probabilities: (*ff*) is the probability that, given a free residue, there is an adjacent free residue to the right; 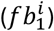 is the probability that, given a free residue, there is an adjacent residue bound to the first section of a ligand of type *i* to the right; 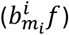 is the probability that, given a residue bound to the final section of a ligand of type *i*, that there is an adjacent free residue to the right; 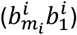 is the probability that, given a residue bound to the final section of a ligand of type *i*, that there is an adjacent residue bound to the first section of a ligand of type *i* to the right.

Using these conditional probabilities, equations can be constructed to model the binding of ligands to a 1D polymer while considering cooperativity and ligand size. Key assumptions are made to enable this approach; ligands are assumed to be either symmetric or bind in a single orientation, and fibril length is assumed to be infinite.^24^

First, only two types of residues can lie to the immediate right of a free residue, yielding

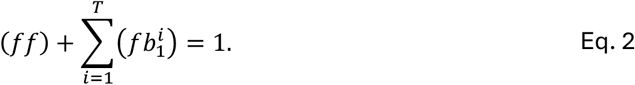

If the final segment of a bound ligand is selected, only a free site or a second bound ligand, denoted as type *j*, can lie to the immediate right, giving

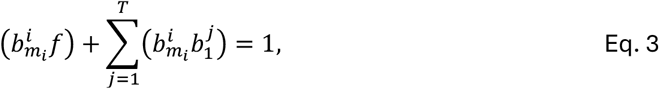

The coverage of a ligand is then defined as the fraction of lattice residues covered by a ligand of type *i*,

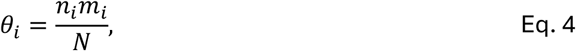

where *n*_*i*_ ligands of type *i* are bound. If a residue is selected at random, the next residue will be free with a probability (1 − *θ*) where 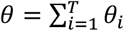, giving

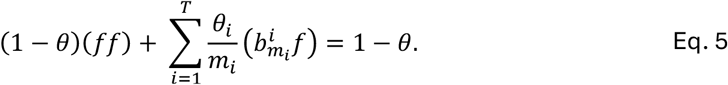

Cooperative binding affects these conditional probabilities and can be accounted for by introducing a cooperativity parameter, *ω*_*ij*_, describing the equilibrium between binding configurations where two ligands are adjacent and non-adjacent (Figure 2b).

While these conditional probabilities describe local adjacency, the number of available binding sites depends on stretches of free residues. Let *α* denote a specific ligand type to be considered. A gap is defined as a sequence of *g* free lattice residues, where *g* = 0 corresponds to adjacent ligands, and a ligand *α* can only bind if *g* ≥ *m*_*α*_. Note that while conditional probabilities are defined per lattice residue, binding site availability is most naturally expressed per gap, following the statistical treatment introduced by McGhee and von Hippel.^24^ This approach allows for the average number of free ligand binding sites per gap, 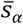, to be computed (see SI),

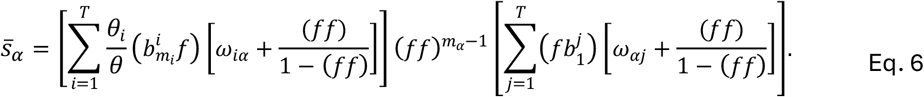

The total number of bound ligands is equal to the number of gaps plus one. The concentration of free binding sites [*s*]can then be written as 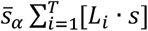, leading to the mass action equation,

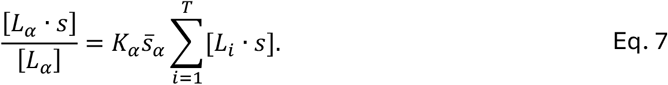

These equations form the basis of a binding model that can consider multiple ligands binding to a linear polymer with multiple sites, allowing for cooperativity and the effect of ligand size to be accounted for. A linear-new site binding mode can be modelled by introducing a second type of binding site for every ligand bound directly to the polymer (Figure 1c). Note that a non-cooperative stacked binding mode where ligand size is 1 (*ω* = 1, *m* = 1) is identical to a standard 1:1 Langmuir isotherm.

### Modelling Saturation Binding

This mathematical model was then applied to compute isotherms and identify unique features for each binding mode. Binding isotherms were first calculated for a single type of ligand binding to a single type of site under four different binding modes: standard 1:1 binding (*m* = 1, ω = 1), linear binding (*m* = 4, ω = 1), cooperative stacked binding (*m* = 1, ω = 4), and linear-new site binding (*m* = 4, ω = 1). The binding isotherms for all binding modes had a similar shape (Figure 3a), although the Scatchard (Figure 3b) and Hill plots (Figure 3c) showed key differences. Plots of the local Hill coefficient, *n*_*H*_, against coverage, more clearly show changes in the Hill plot (Figure 3d). While Scatchard, Hill, and local Hill coefficient plots are sensitive to noise and experimental errors, they more readily display binding mode-specific features than binding isotherms

**Figure 3.**
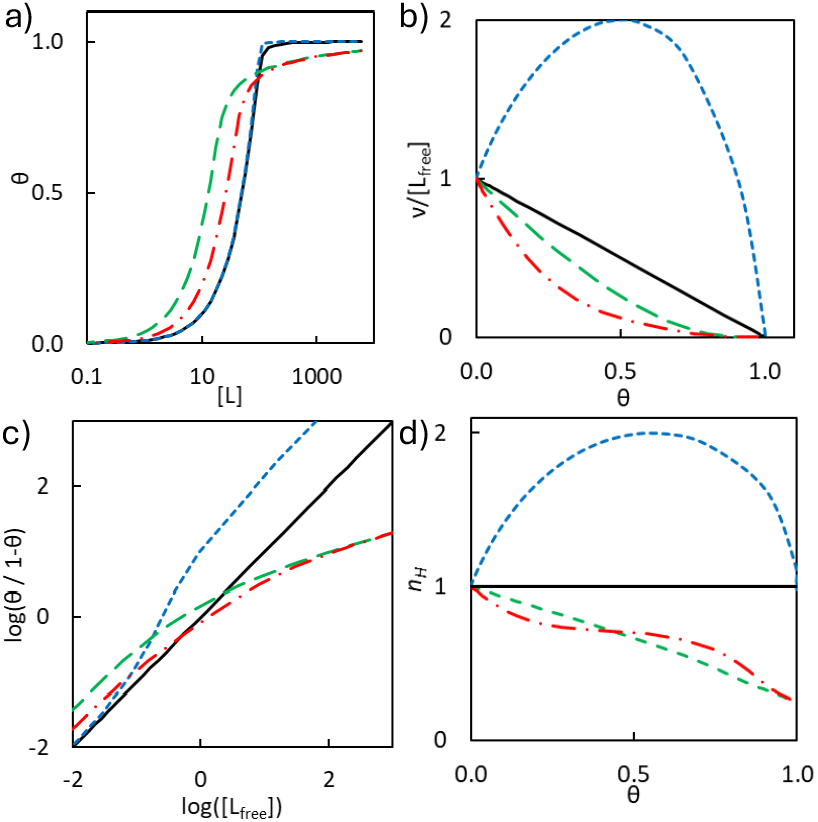
Representative (a) binding isotherms, (b) Scatchard plots, (c) Hill plots, and (d) local Hill coefficient plot for standard 1:1 binding (black line), cooperative stacked binding (*m* = 1, ω = 4, blue short-dash line), linear binding (*m* = 4, ω = 1, green long-dash line), and linear-new site (*m* = 4, ω = 1, red dot-dash line), calculated with *K* = 1 and a lattice site concentration of 100. *v* is the average number of ligands bound per unit lattice.

Cooperative binding produced a concave-down Scatchard plot as expected, and a slope greater than one in the Hill plot. Linear binding led to a shallow binding isotherm at higher concentrations, a concave-up Scatchard plot, and a slope below one in the Hill plot. These features are also characteristic for negative cooperativity and reflect that when a ligand binds, greater than one available binding site is lost due to non-optimal configurations of bound ligands. A linear-new site model demonstrated similar plotted features to a linear model except the Hill coefficient had a non-linear relationship with lattice site occupancy, reflecting the creation of additional binding sites upon ligand binding. Linear and linear-new site models are therefore unlikely to be distinguishable from saturation data alone.

As multiple binding grooves appear to exist on amyloid fibrils, isotherms were computed for a single type of ligand binding to two different types of binding sites. In the absence of any cooperativity and *m* = 1, a single ligand binding to two sites with different affinities yielded a concave-up Scatchard plot resembling linear binding (Figure 4a). The slope of the Hill plot had a local minima below one and returned to unity at low and high occupancy, whereas the Hill coefficient for single-site linear binding was monotonically decreasing with increasing lattice site occupancy (Figure 4b).

**Figure 4.**
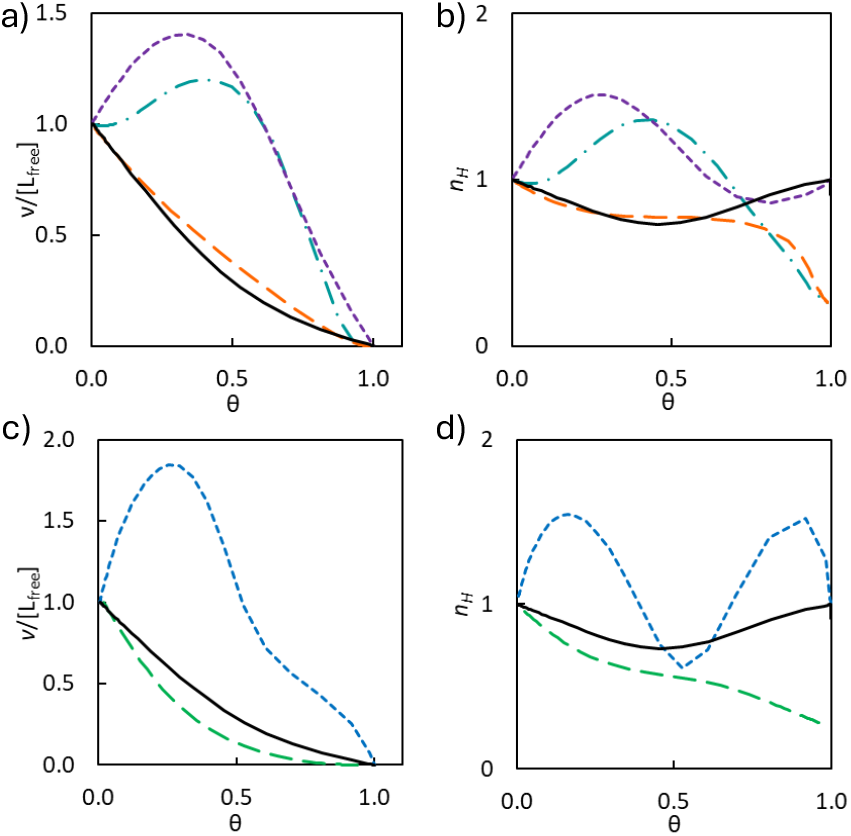
Representative plots for one-ligand, two-site binding models. (a) Scatchard plot and (b) Hill plot for the binding of a ligand to one cooperative stacked and one linear site (teal dash-dot line), one standard and one cooperative stacked site (purple short-dash line), and one standard and one linear site (orange long-dash line) with the same affinity (*K*_*1*_ = *K*_*2*_ = 1). (c) Scatchard plot and (d) Hill plot for the binding of a ligand to two standard sites (black line), two cooperative stacked sites (blue short-dash line), and two linear sites (green long-dash line) with different binding affinities (*K*_*1*_ = 1, *K*_*2*_ = 10). Data for two standard sites (black line, *K*_*1*_ = 1, *K*_*2*_ = 10) are plotted as a reference. Linear binding was modelled with *m* = 4, and stacked binding was modelled with ω = 4.

In other two-site models, a single cooperative site introduced concave-down character in Scatchard plots and a region in the Hill coefficient plot where *n*_*H*_ exceeds unity (Figure 4a-b). A single linear site produced concave-up character in Scatchard plots and a Hill coefficient less than unity at high lattice site occupancies (Figure 4a-b). Two cooperative sites produced a concave-down Scatchard plot with a shoulder and a Hill coefficient plot with two local maxima (Figure 4c-d). Two linear sites produced similar plots to the linear-new site model (Figure 4c-d). If one linear site and one cooperative site was present, the resultant Scatchard and Hill plots combined these features (Figure 4a-b). Characteristic features of Scatchard and Hill coefficient plots are therefore useful in identifying the underlying binding mode present (Table 1). More complex multi-site models will introduce additional features that would likely be challenging to distinguish.

**Table 1.** Diagnostic features of different binding modes.

| Scatchard Features | Local Hill Coefficient Features | Possible Binding Sites |
| --- | --- | --- |
| Concave down | One maxima | 1 cooperative |
|  | Two maxima | 2 cooperative |
|  | One maxima, one minima | 1 standard + 1 cooperative |
| Concave up | Linearly decreasing | 1 linear |
|  | Inflection point | 1 linear new-site,<br>or 2 linear,<br>or 1 standard + 1 linear |
|  | One minima | 2 standard |
| Concave up and concave down sections | Concave up and concave down sections | 1 cooperative + 1 linear |

### Reanalysis of Fluorescence Binding Assays

These models were used to reanalyse previously reported data for the binding of the fluorescent ligand **ThT** to αSyn fibrils derived from the brains of individuals with PD (PD fibrils, Figure 5a,b), and αSyn fibrils aggregated at pH 6.5 (f65 fibrils, Figure 5c,d).^26^ The concentration of lattice sites is required to generate Scatchard and Hill plots, but is generally not known in binding assays using amyloid fibrils. However, diagnostic features were apparent even if imprecise lattice concentrations were used. Additionally, the derived binding model equations were incorporated into the fitting software *Musketeer* to allow accurate fitting and comparison of different binding models.^27^

**Figure 5.**
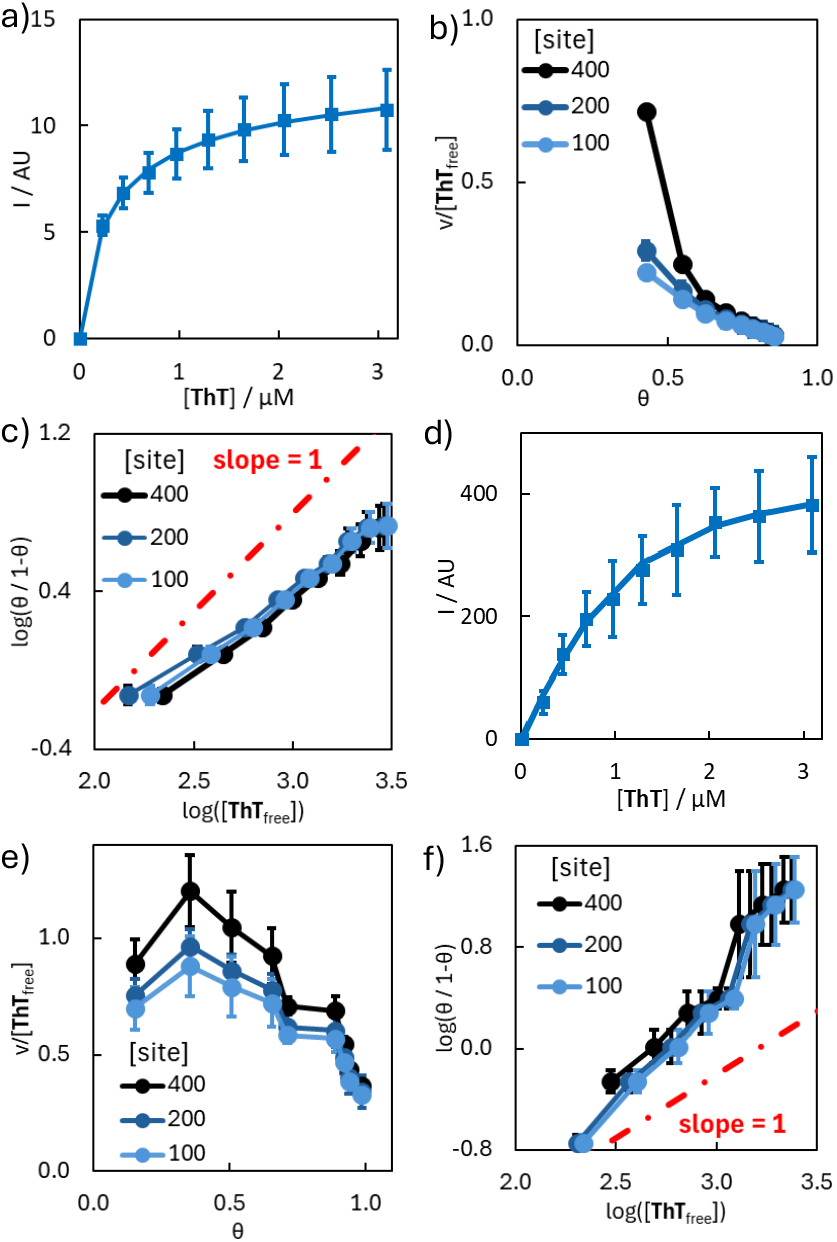
Fluorescence binding assays of **ThT** (λ_ex_ = 440 nm, λ_em_ = 483 nm) to (a) PD fibrils (500 nM), fit to a linear binding model (*m* = 4), with corresponding (b) Scatchard plot and (c) Hill plot; and to (d) f65 fibrils, fit to a non-cooperative stacked binding model, with corresponding (e) Scatchard plot and (f) Hill plot. Data points are the average of at least three experimental measurements with 95% confidence intervals shown.

For **ThT** binding to PD fibrils, both a concave up Scatchard plot and a Hill plot with a slope less than one was generated at multiple plausible lattice site concentrations (Figure 5a-c). These features are indicative of a linear binding mode. Fitting the raw data with a linear binding mode and ligand length of four gave a superior fit (RMSE = 0.09) compared to a non-cooperative stacked model (RMSE = 0.17). A ligand size of four agrees with cryo-EM structures of linearly bound **ThT**, which spans four protein monomers (PDB ID: 8×7B).^19^ For **ThT** binding to f65 fibrils, both a concave-down Scatchard plot and a Hill plot with a slope greater than one was generated at multiple plausible lattice site concentrations (Figure 5d-f). These features are indicative of a cooperative stacked binding mode. Fitting the raw data with a cooperative binding mode yielded a cooperativity parameter ω of 1.73 and a lower RMSE than a non-cooperative stacked model. However, as including ω introduces an additional fitted parameter and increases model complexity, models were compared using the Akaika Information Criterion with small-sample correction (AICc). The cooperative and non-cooperative models were statistically indistinguishable (ΔAICc = 0.9) and the non-cooperative model was therefore preferred for simplicity.^28,29^ These examples demonstrate that both linear and stacked binding modes are plausible as the primary binding mode of ligands, and that the same ligand is able to engage in different binding modes to different fibril structures.

### Radioligand Self-Displacement Assays

Data indicative of different binding modes can also be obtained by comparing radioligand saturation and self-displacement assays. In self-displacement assays, involving the displacement of a radiolabelled ligand with a structurally identical unlabelled ligand, effects from cooperative and linear binding modes are equivalent between the labelled and unlabelled ligands. However, effects from cooperative and linear binding modes will be present in saturation binding assays, meaning a difference in the dissociation constant of a ligand measured using a radioligand saturation assay (*K*_*d*_) and a radioligand self-displacement assay (*K*_*i*_) may indicate the presence of a linear or cooperative binding mode. Specifically, a *K*_*i*_ > *K*_*d*_ indicates cooperative binding, whereas a *K*_*i*_ < *K*_*d*_ indicates linear binding.

There are 23 examples from the literature where the dissociation constant of a radioligand has been measured using both a saturation and self-displacement assay, and screened against the same fibril target, in the same publication (Figure 6a).^30–52^ Previous work has found that the median variation in error of reported binding constants in single studies was 0.35 log units.^4^ A threshold of 1.0 log units was chosen to constitute a significant difference, and four examples were found (Figure 6b).^30–32^ **TZDM** and **THK523** had a higher dissociation constant measured from self-displacement assays, indicating cooperative binding may be occurring, while **PiB** and **MODAG-001** had a higher dissociation constant measured from saturation assays, indicating linear binding may be occurring. While **TZDM** and **PiB** are structurally similar, they appear to bind via different modes to different targets. Given the ubiquity of the scaffolds of these four ligands, and that differences in dissociation constant may be produced that are less than an order of magnitude, it is likely many reported ligands undergo cooperative or linear binding.

**Figure 6.**
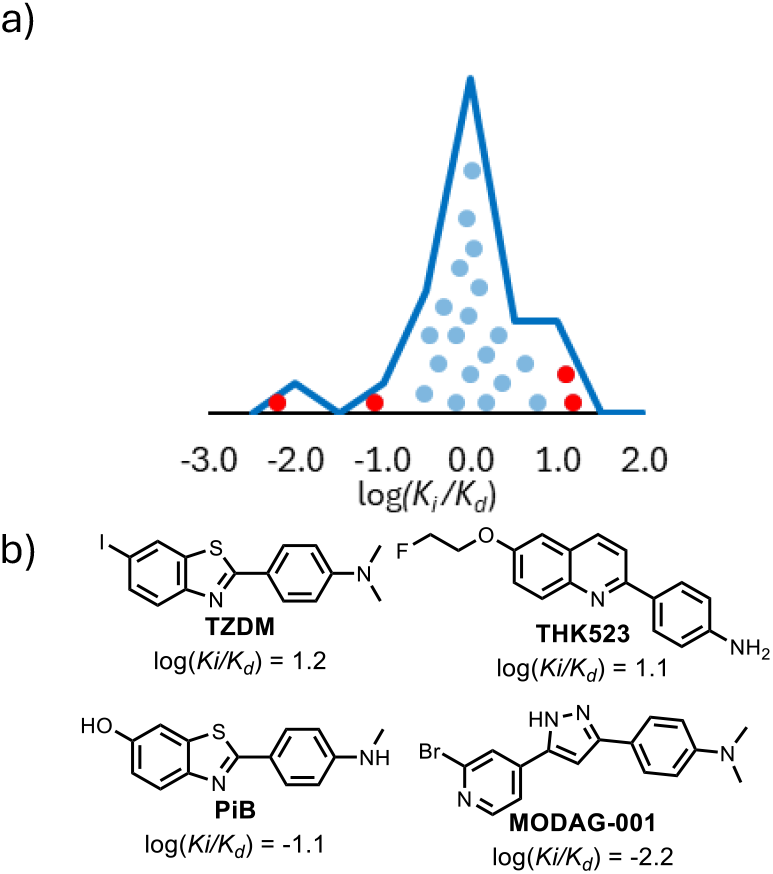
(a) Distribution of log(*K*_*i*_/*K*_*d*_) ratios for radioligands with dissociation constants measured from saturation and self-displacement assays in the same publication. (b) Ligands with a difference in dissociation constants of at least an order of magnitude, shown in red in panel (a).

### Designing Ligands for Complex Binding Modes

The presence of linear and stacked binding mode suggests new design principles can be applied for developing high-affinity ligands. For linear binding, long and narrow ligands may yield stronger binding. For stacked binding, ligands should be flat to enable stacking, and cooperative interactions may be possible. The binding mode present also determines the maximum number of ligands that can bind to a target, which may be an important consideration in applications such as imaging. To test these principles, ligands were prepared based on **ThT** as a known binding motif. Ligand **1** incorporates two **ThT** headgroups joined to favour linear binding by increasing the length of the ligand. A short linker was chosen to prevent binding in different grooves, and to minimise any added entropic penalty to binding.^53^ Ligand **2** incorporates a naphthalene diamine (NDI) moiety that self-associates in aqueous media to form supramolecular polymers driven by π-π interactions.^54–56^

Binding assays were performed using **ThT, 1**, and **2** against αSyn fibrils (Figure 7). Two fibril morphologies were used; αSyn-PBS fibrils were prepared in 1xPBS buffer, and αSyn-Tris fibrils were prepared in Tris buffer. Different fibril morphologies, while composed of the same protein, have different chemical and biological properties including distinct ligand-binding sites.^8^ The fluorescence emission of the ThT headgroups (λ_ex_ = 440 nm) were monitored, alongside the fluorescence of the NDI moiety of **2** (λ_ex_ = 332 nm). Models with the same number of fitted parameters were evaluated by comparing the RMSE of the fit, and models with different numbers of fitted parameters were evaluated and compared using AICc.

**Figure 7.**
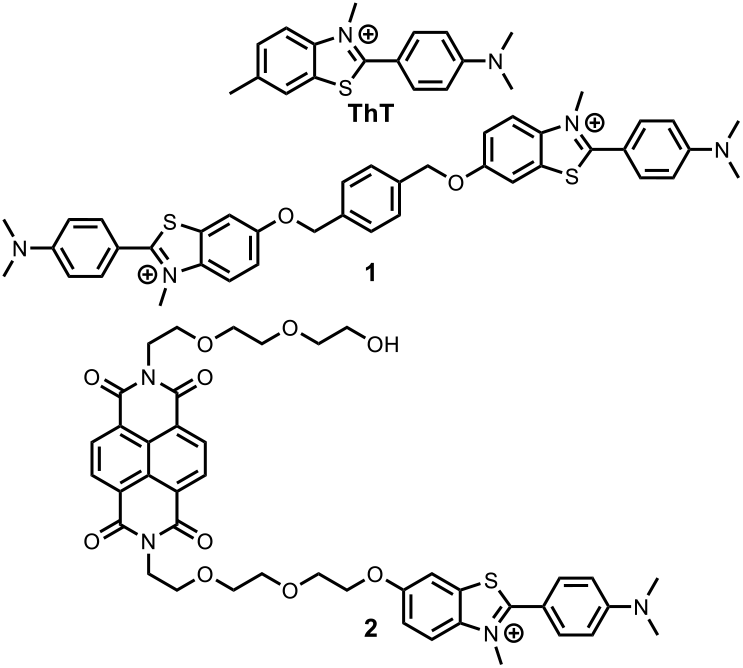
Structures of the **ThT** derivatives used.

Binding of **ThT** to αSyn fibrils produced an enhanced emission at 483 nm upon excitation at 440 nm (Figure 8a-b). Scatchard and Hill analyses suggest that **ThT** bound to αSyn-PBS fibrils in a non-cooperative stacked binding mode (Figure 8b,d,f, see SI), but potentially bound to αSyn-Tris fibrils in a linear binding mode (Figure 8b,c,e, see SI). Data fitting supported these observations, where a non-cooperative stacked binding mode produced the best fit for binding to αSyn-PBS (*K*_*d*_ = 1,600 ± 500 nM), and a linear binding mode (*m* = 4) gave the best fit for binding to αSyn-Tris (*K*_*d*_ = 8,000 ± 1,000 nM).

**Figure 8.**
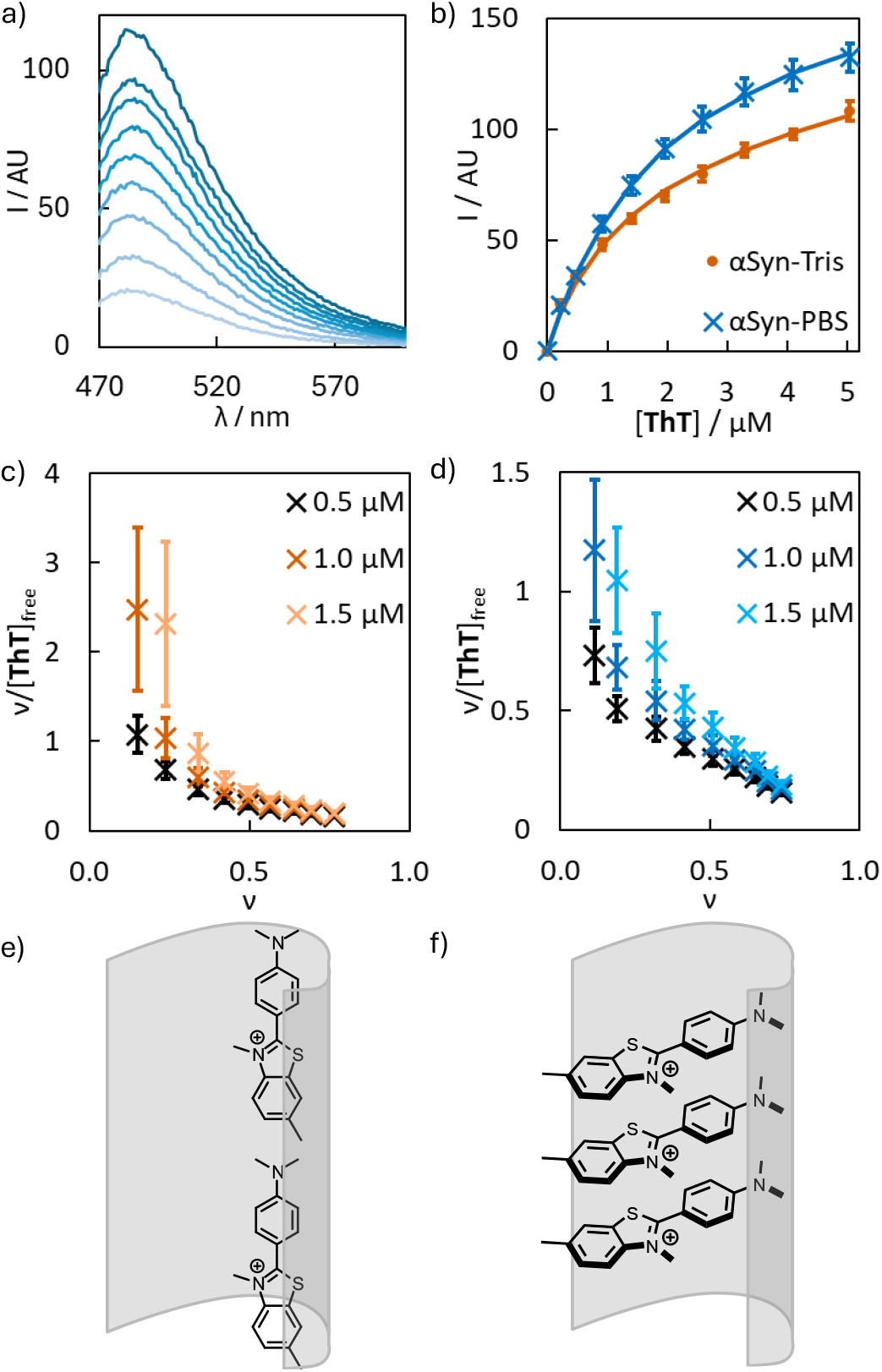
Data from fluorescence binding assays of **ThT** (λ_ex_ = 440 nm) with αSyn-Tris and αSyn-PBS fibrils (500 nM) in 1xPBS (pH 7.4). (a) Representative emission spectra showing increasing **ThT** concentrations added to αSyn-Tris fibrils. (b) Fluorescence titration data (λ_em_ = 483 nm). Lines are the best fit to a linear binding isotherm (*m* = 4) for αSyn-Tris fibrils and a non-cooperative stacked binding isotherm for αSyn-PBS fibrils. Scatchard plots for **ThT** binding to (c) αSyn-Tris fibrils and (d) αSyn-PBS fibrils. Schematics illustrating the proposed binding modes of **ThT** to (e) αSyn-Tris and (f) αSyn-PBS fibrils. Data points are the average of at least three experimental measurements with 95% confidence intervals shown.

Distinct binding isotherms were observed for **1**. The emission of the **ThT** headgroup at 483 nm decreased at higher ligand concentrations, and a new emission peak appeared at 582 nm (Figure 9a-c). This red-shifted emission is similar to experimental measurements of **ThT** excimers in aqueous solution.^57^ Excimer formation can be approximated using the binding model outlined by assuming that adjacently bound ligands are able to form excimers and have a different emission spectra. Using this model, accurate fits were obtained for ligand **1** binding to αSyn-PBS (*K*_*d*_ = 300 ± 80 nM) and αSyn-Tris (*K*_*d*_ = 3,000 ± 600 nM) in a linear binding mode (*m* = 8) showing improved binding affinities compared to **ThT** (Figure 9b-d). A ligand size of eight aligns with expectations based on cryo-EM structures.

**Figure 9.**
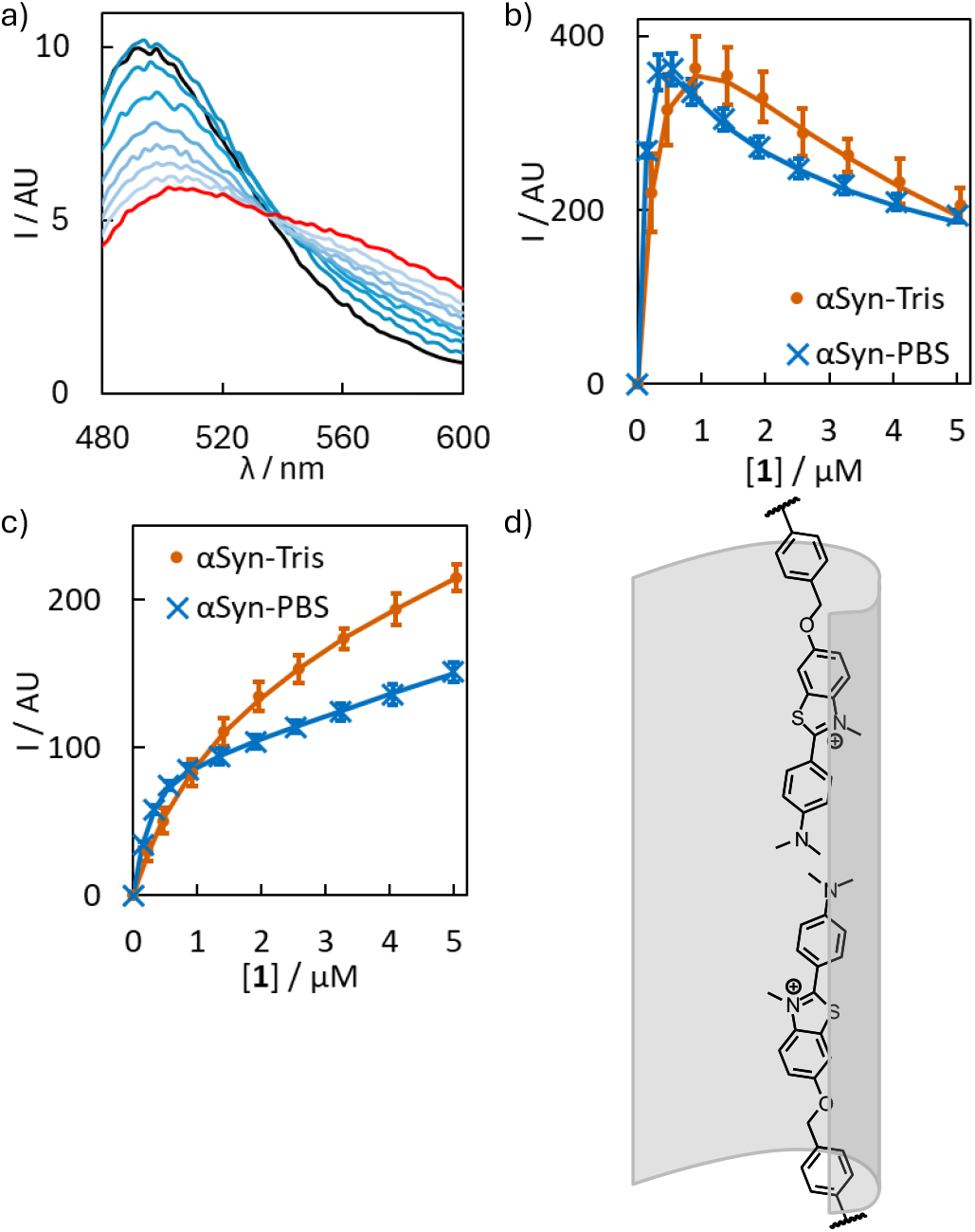
Data from fluorescence binding assays of **1** (λ_ex_ = 440 nm) with αSyn-Tris and αSyn-PBS fibrils (500 nM) in 1xPBS (pH 7.4). (a) Representative emission spectra for increasing concentrations of **1** added to αSyn-PBS fibrils (black line: 326 nM **1**; black line: 5.0 µM **1**). Fluorescence titration data monitored at (b) λ_em_ = 483 nm and (c) λ_em_ = 582 nm for the addition of **1** to αSyn fibrils. Lines are the best fit to a linear binding isotherm (*m* = 8) accounting for adjacency. (d) Schematic illustrating the proposed binding mode of **1.** Data points are the average of at least three experimental measurements with 95% confidence intervals shown.

For the binding of **2**, NDI excitation at 332 nm produced a new emission band at 488 nm upon binding both fibril morphologies (Figure 10a). This emission is red-shifted and stronger than the emission observed in the absence of fibril, in agreement with literature observations for the formation of π-stacked NDI supramolecular polymers in aqueous media.^58^ A non-cooperative stacked binding model considering adjacency gave the best fit for both fibril targets (Figure 10b-d). Binding of **2** to αSyn-PBS (*K*_*d*_ = 1,800 ± 100 nM) was not enhanced compared to **ThT**, but binding to αSyn-Tris (*K*_*d*_ = 600 ± 60 nM) was significantly stronger.

**Figure 10.**
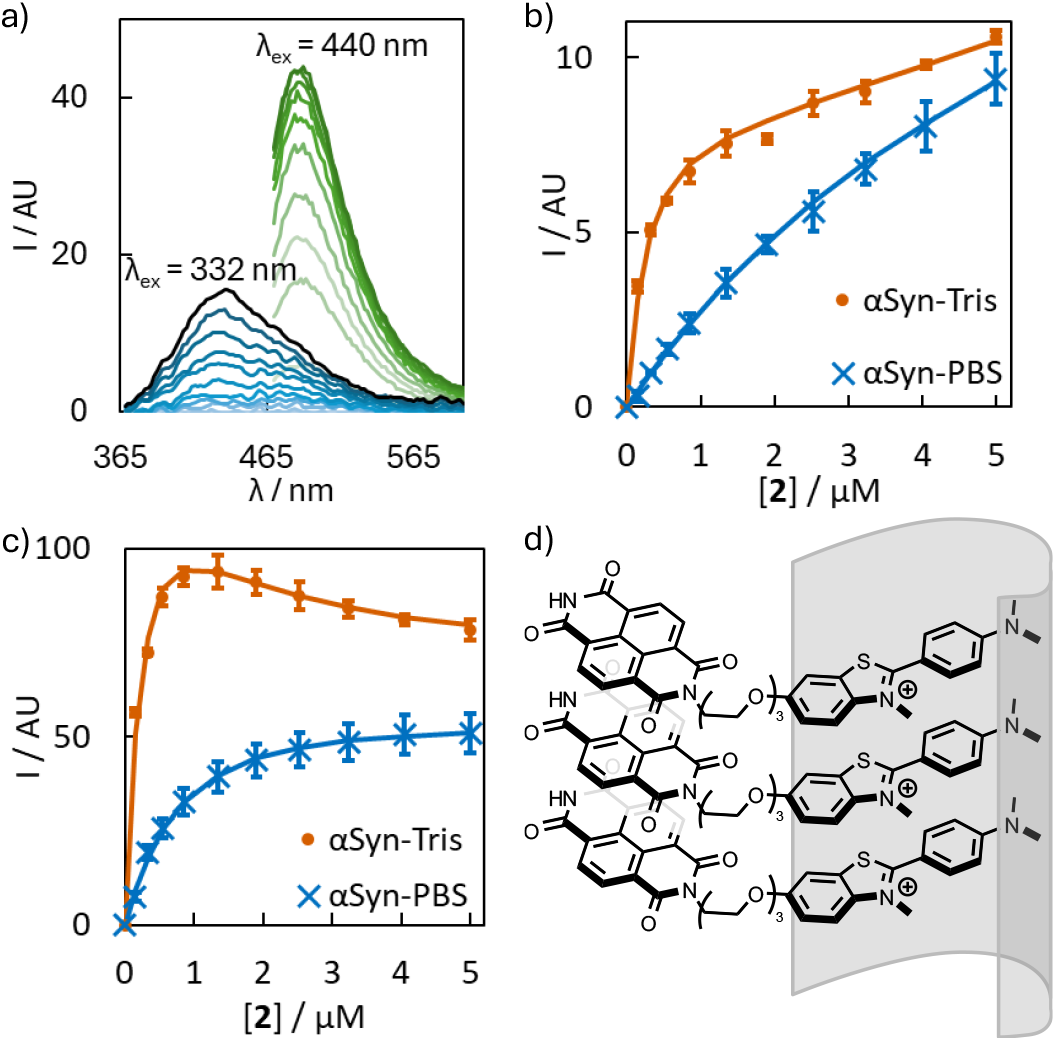
Data from fluorescence binding assays of **2** with αSyn-Tris and αSyn-PBS fibrils (500 nM) in 1xPBS (pH 7.4). (a) Representative emission spectra (λ_ex_ = 332 nm, λ_ex_ = 440 nm) for increasing concentrations of **2** added to αSyn-PBS fibrils. Fluorescence titration data monitored at (b) λ_ex_ = 332 nm; λ_em_ = 440 nm and (c) λ_ex_ = 440; λ_em_ = 483 nm for the addition of **2** to αSyn fibrils with fits to a non-cooperative stacked binding isotherm accounting for adjacency. (d) Schematic illustrating the proposed binding mode of **2.** Data points are the average of at least three experimental measurements with 95% confidence intervals shown.

These results support the existence of distinct stacked and linear binding modes, and provide a proof-of-principle that binding mode-informed ligand design can both influence binding mechanism and enhance affinity. Ligand **1** exhibited enhanced affinity and enforced a linear binding mode. In contrast, NDI functionalisation in **2** did not induce cooperativity but did promote a stacked binding mode, enhanced binding to αSyn-Tris, and provided an additional fluorescent readout.

## Conclusion

This work provides analytical evidence that ligands bind to amyloid fibrils in linear and stacked binding modes, alongside a mathematical framework for distinguishing these mods using experimental binding data. These binding modes produce diagnostic features in binding assays that enable their discrimination, and predictions made using this framework are consistent with previously reported saturation and competition binding assays. Knowledge of these binding modes can inform the design of ligands engineered to preferentially bind in either a linear or stacked mode. Fluorescence measurements of ligands **1** and **2** are consistent with excimer formation and NDI self-association, supporting the proposed binding modes and enabling stronger binding to occur. Together these results support the existence of linear and stacked binding modes at concentrations relevant to binding assays and biological applications, and provide a quantitative framework for more accurately interpreting binding assays and guiding the rational design of ligands for amyloid fibrils.

## Conflicts of interest

There are no conflicts of interest to declare.

## Data availability

All experimental procedures, analytical data, and spectroscopic data are provided in the supplementary information. The dataset of radioligands used is provided as a .csv file, and python code used is provided as .ipynb and .py files.

## Acknowledgements

T.S.C. thanks the Cambridge Philosophical Society Henslow Fellowship for funding, Professor Christopher Hunter for useful discussions and providing feedback on manuscript drafts, Ben Iddon for providing feedback on manuscript drafts, and Daniil Soloviev for providing useful guidance on the modification of *Musketeer* code. T.S.C thanks the EPSRC Underpinning MultiUser Equipment Call (EP/P030467/1).

## Notes

### Competing Interest Statement

The authors have declared no competing interest.

## References

(1) Chiti, F.; Dobson, C. M. Protein Misfolding, Amyloid Formation, and Human Disease: A Summary of Progress Over the Last Decade. Annu. Rev. Biochem. 2017, 86 (1), 27–68. 10.1146/annurev-biochem-061516-045115.

(2) Parkinson, J. An Essay on the Shaking Palsy.; Sherwood, Neely and Jones: London, 1817.

(3) Alzheimer, A. Über Einen Eigenartigen Schweren ErkrankungsprozeB Der Hirnrinde. Neurol. Cent. 1906, 23, 1129–1136.

(4) Chisholm, T. S.; Hunter, C. A. A Closer Look at Amyloid Ligands, and What They Tell Us about Protein Aggregates. Chem. Soc. Rev. 2024, 53, 1354–1374. 10.1039/D3CS00518F.

(5) Chisholm, T. S.; Hunter, C. A. Ligands for Protein Fibrils of Amyloid-β, α-Synuclein, and Tau. Chem. Rev. 2025, 125 (11), 5282–5348. 10.1021/acs.chemrev.4c00838.

(6) Mathis, C. A.; Lopresti, B. J.; Ikonomovic, M. D.; Klunk, W. E. Small-Molecule PET Tracers for Imaging Proteinopathies. Semin. Nucl. Med. 2017, 47 (5), 553–575. 10.1053/j.semnuclmed.2017.06.003.

(7) Aliyan, A.; Cook, N. P.; Martí, A. A. Interrogating Amyloid Aggregates Using Fluorescent Probes. Chem. Rev. 2019, 119 (23), 11819–11856. 10.1021/acs.chemrev.9b00404.

(8) Chisholm, T. S.; Hunter, C. A. Ligand Profiling to Characterize Different Polymorphic Forms of α-Synuclein Aggregates. J. Am. Chem. Soc. 2023, 145 (49), 27030–27037. 10.1021/jacs.3c10521.

(9) Zhuang, Z. P.; Kung, M. P.; Hou, C.; Skovronsky, D. M.; Gur, T. L.; Plössl, K.; Trojanowski, J. Q.; Lee, V. M. Y.; Kung, H. F. Radioiodinated Styrylbenzenes and Thioflavins as Probes for Amyloid Aggregates. J. Med. Chem. 2001, 44 (12), 1905– 1914. 10.1021/jm010045q.

(10) Lockhart, A.; Ye, L.; Judd, D. B.; Merrittu, A. T.; Lowe, P. N.; Morgenstern, J. L.; Hong, G.; Gee, A. D.; Brown, J. Evidence for the Presence of Three Distinct Binding Sites for the Thioflavin T Class of Alzheimer’s Disease PET Imaging Agents on β-Amyloid Peptide Fibrils. J. Biol. Chem. 2005, 280 (9), 7677–7684. 10.1074/jbc.M412056200.

(11) Sulatskaya, A. I.; Kuznetsova, I. M.; Turoverov, K. K. Interaction of Thioflavin T with Amyloid Fibrils: Fluorescence Quantum Yield of Bound Dye. J. Phys. Chem. B 2012, 116, 2358–2544. 10.1021/jp2083055.

(12) Sulatskaya, A.; Rodina, N.; Polyakov, D.; Sulatsky, M.; Artamonova, T.; Khodorkovskii, M.; Shavlovsky, M.; Kuznetsova, I.; Turoverov, K. Structural Features of Amyloid Fibrils Formed from the Full-Length and Truncated Forms of Beta-2-Microglobulin Probed by Fluorescent Dye Thioflavin T. Int. J. Mol. Sci. 2018, 19 (9), 2762. 10.3390/ijms19092762.

(13) Groenning, M. Binding Mode of Thioflavin T and Other Molecular Probes in the Context of Amyloid Fibrils-Current Status. J. Chem. Biol. 2010, 3 (1), 1–18. 10.1007/s12154-009-0027-5.

(14) Krebs, M. R. H.; Bromley, E. H. C.; Donald, A. M. The Binding of Thioflavin-T to Amyloid Fibrils: Localisation and Implications. J. Struct. Biol. 2005, 149 (1), 30–37. 10.1016/j.jsb.2004.08.002.

(15) Groenning, M.; Norrman, M.; Flink, J. M.; van de Weert, M.; Bukrinsky, J. T.; Schluckebier, G.; Frokjaer, S. Binding Mode of Thioflavin T in Insulin Amyloid Fibrils. J. Struct. Biol. 2007, 159 (3), 483–497. 10.1016/j.jsb.2007.06.004.

(16) Biancalana, M.; Koide, S. Molecular Mechanism of Thioflavin-T Binding to Amyloid Fibrils. Biochim. Biophys. Acta, Proteins Proteoics 2010, 1804, 1405–1412. 10.1016/j.bbapap.2010.04.001.

(17) Carter, D. B.; Chou, K.-C. A Model for Structure-Dependent Binding of Congo Red to Alzheimer β-Amyloid Fibrils. Neurobiol. Aging 1998, 19 (1), 37–40. 10.1016/S0197-4580(97)00164-4.

(18) Tao, Y.; Xia, W.; Zhao, Q.; Xiang, H.; Han, C.; Zhang, S.; Gu, W.; Tang, W.; Li, Y.; Tan, L.; Li, D.; Liu, C. Structural Mechanism for Specific Binding of Chemical Compounds to Amyloid Fibrils. Nat. Chem. Biol. 2023, 1–11. 10.1038/s41589-023-01370-x.

(19) Liu, K.; Tao, Y.; Zhao, Q.; Xia, W.; Li, X.; Zhang, S.; Yao, Y.; Xiang, H.; Han, C.; Tan, L.; Sun, B.; Li, D.; Li, A.; Liu, C. Binding Adaptability of Chemical Ligands to Polymorphic α-Synuclein Amyloid Fibrils. Proc. Natl. Acad. Sci. 2024, 121 (35), e2321633121. 10.1073/pnas.2321633121.

(20) Shi, Y.; Ghetti, B.; Goedert, M.; Scheres, S. H. W. Cryo-EM Structures of Chronic Traumatic Encephalopathy Tau Filaments with PET Ligand Flortaucipir. J. Mol. Biol. 2023, 435 (11), 168025. 10.1016/j.jmb.2023.168025.

(21) Seidler, P. M.; Boyer, D. R.; Sawaya, M. R.; Ge, P.; Shin, W. S.; DeTure, M. A.; Dickson, D. W.; Jiang, L.; Eisenberg, D. S. CryoEM Reveals How the Small Molecule EGCG Binds to Alzheimer’s Brain-Derived Tau Fibrils and Initiates Fibril Disaggregation. Nat. Commun. 2022, 13, 5451. 10.1101/2020.05.29.124537.

(22) Shi, Y.; Murzin, A.G.; Falcon, B.; Epstein, A.; Machin, J.; Tempest, P.; Newell, K. L.; Vidal, R.; Garringer, H.J.; Sahara, N.; Higuchi, M.; Ghetti, B.; Jang, M.K.; Scheres, S.H.W.; Goedert, M. Cryo-EM Structures of Tau Filaments from Alzheimer’s Disease with PET Ligand APN-1607. Acta Neuropathol. (Berl.) 2021, 1, 3. 10.1007/s00401-021-02294-3.

(23) Smith, M. S.; DeGrado, W. F.; Grabe, M.; Shoichet, B. K. A Cooperative Model for Symmetric Ligand Binding to Protein Fibrils. Biochem. 2025, 64 (15), 3382–3392. 10.1021/acs.biochem.5c00068.

(24) McGhee, J. D.; von Hippel, P. H. Theoretical Aspects of DNA-Protein Interactions: Co-Operative and Non-Co-Operative Binding of Large Ligands to a One-Dimensional Homogeneous Lattice. J. Mol. Biol. 1974, 86 (2), 469–489. 10.1016/0022-2836(74)90031-X.

(25) Villaluenga, J. P. G.; Brunete, D.; Cao-García, F. J. Competitive Ligand Binding Kinetics to Linear Polymers. Phys. Rev. E 2023, 107 (2), 024401. 10.1103/PhysRevE.107.024401.

(26) Chisholm, T. S.; Melki, R.; Hunter, C. Ligand Profiling as a Diagnostic Tool to Differentiate Patient-Derived α-Synuclein Polymorphs. ACS Chem. Neurosci. 2024, 15 (10), 2080–2088. 10.1021/acschemneuro.4c00178.

(27) O. Soloviev, D.; A. Hunter, C. Musketeer: A Software Tool for the Analysis of Titration Data. Chem. Sci. 2024, 15 (37), 15299–15310. 10.1039/D4SC03354J.

(28) Hurvich, C.; Tsai, C.-L. Regression and Time Series Model Selection in Small Samples. Biometrika 1989, 76 (2), 297–307. 10.1093/biomet/76.2.297.

(29) Sugiura, N. Further Analysis of the Data by Akaike’s Information Criterion and the Finite Corrections: Further Analysis of the Data by Akaike’ s. Commun. Stat. -Theory Methods 1978, 7 (1), 13–26. 10.1080/03610927808827599.

(30) Cai, L.; Qu, B.; Hurtle, B. T.; Dadiboyena, S.; Diaz-Arrastia, R.; Pike, V. W. Candidate PET Radioligand Development for Neurofibrillary Tangles: Two Distinct Radioligand Binding Sites Identified in Postmortem Alzheimer’s Disease Brain Graphical Abstract HHS Public Access. ACS Chem. Neurosci. 2016, 7 (7), 897–911. 10.1021/acschemneuro.6b00051.

(31) Di Nanni, A.; Saw, R. S.; Battisti, U. M.; Bowden, G. D.; Boeckermann, A.; Bjerregaard-Andersen, K.; Pichler, B. J.; Herfert, K.; Herth, M. M.; Maurer, A. A Fluorescent Probe as a Lead Compound for a Selective α-Synuclein PET Tracer: Development of a Library of 2-Styrylbenzothiazoles and Biological Evaluation of [18F]PFSB and [18F]MFSB. ACS Omega 2023, 8 (34), 31450–31467. 10.1021/acsomega.3c04292.

(32) Kung, H. F.; Lee, C. W.; Zhuang, Z. P.; Kung, M. P.; Hou, C.; Plössl, K. Novel Stilbenes as Probes for Amyloid Plaques. J. Am. Chem. Soc. 2001, 123 (50), 12740–12741. 10.1021/ja0167147.

(33) Choi, S. R.; Golding, G.; Zhuang, Z.; Zhang, W.; Lim, N.; Hefti, F.; Benedum, T. E.; Kilbourn, M. R.; Skovronskyhank, D.; Kung, H. F. Preclinical Properties of 18F-AV-45: A PET Agent for Aβ Plaques in the Brain. J. Nucl. Med. 2009, 50 (11), 1887–1894. 10.2967/jnumed.109.065284.

(34) Declercq, L.; Rombouts, F.; Koole, M.; Fierens, K.; Mariën, J.; Langlois, X.; Andrés, J. I.; Schmidt, M.; MacDonald, G.; Moechars, D.; Vanduffel, W.; Tousseyn, T.; Vandenberghe, R.; Van Laere, K.; Verbruggen, A.; Bormans, G. Preclinical Evaluation of 18F-JNJ64349311, a Novel PET Tracer for Tau Imaging. J. Nucl. Med. 2017, 58 (6), 975–981. 10.2967/jnumed.116.185199.

(35) Duan, X. H.; Qiao, J. P.; Yang, Y.; Cui, M. C.; Zhou, J. N.; Liu, B. L. Novel Anilinophthalimide Derivatives as Potential Probes for β-Amyloid Plaque in the Brain. Bioorg. Med. Chem. 2010, 18 (3), 1337–1343. 10.1016/j.bmc.2009.12.023.

(36) Hsieh, C. J.; Ferrie, J. J.; Xu, K.; Lee, I.; Graham, T. J. A.; Tu, Z.; Yu, J.; Dhavale, D.; Kotzbauer, P.; Petersson, E. J.; Mach, R. H. Alpha Synuclein Fibrils Contain Multiple Binding Sites for Small Molecules. ACS Chem. Neurosci. 2018, 9 (11), 2521–2527. 10.1021/acschemneuro.8b00177.

(37) Johnson, A. E.; Jeppsson, F.; Sandell, J.; Wensbo, D.; Neelissen, J. A. M.; Juréus, A.; Ström, P.; Norman, H.; Farde, L.; Svensson, S. P. S. AZD2184: A Radioligand for Sensitive Detection of β-Amyloid Deposits. J. Neurochem. 2009, 108 (5), 1177– 1186. 10.1111/j.1471-4159.2008.05861.x.

(38) Kung, M. P.; Hou, C.; Zhuang, Z. P.; Skovronsky, D.; Kung, H. F. Binding of Two Potential Imaging Agents Targeting Amyloid Plaques in Postmortem Brain Tissues of Patients with Alzheimer’s Disease. Brain Res. 2004, 1025 (1–2), 98–105. 10.1016/j.brainres.2004.08.004.

(39) Lemoine, L.; Saint-Aubert, L.; Marutle, A.; Antoni, G.; Eriksson, J. P.; Ghetti, B.; Okamura, N.; Nennesmo, I.; Gillberg, P. G.; Nordberg, A.; Gillberg, G.; Nordberg, A. Visualization of Regional Tau Deposits Using 3H-THK5117 in Alzheimer Brain Tissue. Acta Neuropathol. Commun. 2015, 3, 40. 10.1186/s40478-015-0220-4.

(40) Lemoine, L.; Gillberg, P.-G.; Svedberg, M.; Stepanov, V.; Jia, Z.; Huang, J.; Nag, S.; Tian, H.; Ghetti, B.; Okamura, N.; Higuchi, M.; Halldin, C.; Nordberg, A. Comparative Binding Properties of the Tau PET Tracers THK5117, THK5351, PBB3, and T807 in Postmortem Alzheimer Brains. Alzheimer’s Res. Ther. 2017, 9, 96. 10.1186/s13195-017-0325-z.

(41) Miranda-Azpiazu, P.; Svedberg, M.; Higuchi, M.; Ono, M.; Jia, Z.; Sunnemark, D.; Elmore, C. S.; Schou, M.; Varrone, A. Identification and in Vitro Characterization of C05-01, a PPB3 Derivative with Improved Affinity for Alpha-Synuclein. Brain Res. 2020, 1749, 147131. 10.1016/j.brainres.2020.147131.

(42) Okamura, N.; Furumoto, S.; Harada, R.; Tago, T.; Yoshikawa, T.; Fodero-Tavoletti, M.; Mulligan, R. S.; Villemagne, V. L.; Akatsu, H.; Yamamoto, T.; Arai, H.; Iwata, R.; Yanai, K.; Kudo, Y. Novel 18F-Labeled Arylquinoline Derivatives for Noninvasive Imaging of Tau Pathology in Alzheimer Disease. J. Nucl. Med. 2013, 54 (8), 1420– 1427. 10.2967/jnumed.112.117341.

(43) Ono, M.; Yoshida, N.; Ishibashi, K.; Haratake, M.; Arano, Y.; Mori, H.; Nakayama, M. Radioiodinated Flavones for in Vivo Imaging of β-Amyloid Plaques in the Brain. J. Med. Chem. 2005, 48 (23), 7253–7260. 10.1021/jm050635e.

(44) Ono, M.; Maya, Y.; Haratake, M.; Ito, K.; Mori, H.; Nakayama, M. Aurones Serve as Probes of β-Amyloid Plaques in Alzheimer’s Disease. Biochem. Biophys. Res. Commun. 2007, 361 (1), 116–121. 10.1016/j.bbrc.2007.06.162.

(45) Ono, M.; Haratake, M.; Mori, H.; Nakayama, M. Novel Chalcones as Probes for in Vivo Imaging of β-Amyloid Plaques in Alzheimer’s Brains. Bioorg. Med. Chem. 2007, 15 (21), 6802–6809. 10.1016/j.bmc.2007.07.052.

(46) Ono, M.; Sahara, N.; Kumata, K.; Ji, B.; Ni, R.; Koga, S.; Dickson, D. W.; Trojanowski, J. Q.; Lee, V. M.-Y.; Yoshida, M.; Hozumi, I.; Yoshiyama, Y.; Van Swieten, J. C.; Nordberg, A.; Suhara, T.; Zhang, M.-R.; Higuchi, M. Distinct Binding of PET Ligands PBB3 and AV-1451 to Tau Fibril Strains in Neurodegenerative Tauopathies Maiko. Brain J. Neurol. 2017, 140 (3), 764–780. 10.1093/brain/aww339.

(47) Tago, T.; Furumoto, S.; Okamura, N.; Harada, R.; Adachi, H.; Ishikawa, Y.; Yanai, K.; Iwata, R.; Kudo, Y. Structure-Activity Relationship of 2-Arylquinolines as PET Imaging Tracers for Tau Pathology in Alzheimer Disease. J. Nucl. Med. 2016, 57 (4), 608–614. 10.2967/jnumed.115.166652.

(48) Walji, A. M.; Hostetler, E.; Grrshock, T. J.; Li, J.; Moore, K. P.; Bennacef, I.; Mulhearn, J.; Selnick, H.; Wang, Y.; Yang, K.; Fu, J. Pyrrolo[2,3-C]Pyridines as Imaging Agents for Neurofibrilary Tangles. US10022461B2, 2018.

(49) Wang, Y.; Klunk, W. E.; Huang, G. F.; Debnath, M. L.; Holt, D. P.; Mathis, C. A. Synthesis and Evaluation of 2-(3′-Iodo-4′-Aminophenyl)-6-Hydroxybenzothiazole for in Vivo Quantitation of Amyloid Deposits in Alzheimer’s Disease. J. Mol. Neurosci. 2002, 19 (1–2), 11–16. 10.1007/s12031-002-0004-8.

(50) Wang, Y.; Klunk, W. E.; Debnath, M. L.; Huang, G. F.; Holt, D. P.; Shao, L.; Mathis, C. Development of a PET/SPECT Agent for Amyloid Imaging in Alzheimer’s Disease. J. Mol. Neurosci. 2004, 24 (1), 55–62. 10.1385/jmn:24:1:055.

(51) Hellström-Lindahl, E.; Westermark, P.; Antoni, G.; Estrada, S. In Vitro Binding of [3H]PIB to Human Amyloid Deposits of Different Types. Amyloid 2014, 21 (1), 21– 27. 10.3109/13506129.2013.860895.

(52) Xiang, J.; Tao, Y.; Xia, Y.; Luo, S.; Zhao, Q.; Li, B.; Zhang, X.; Sun, Y.; Xia, W.; Zhang, M.; Kang, S. S.; Ahn, E.-H.; Liu, X.; Xie, F.; Guan, Y.; Yang, J. J.; Bu, L.; Wu, S.; Wang, X.; Cao, X.; Liu, C.; Zhang, Z.; Li, D.; Ye, K. Development of an α-Synuclein Positron Emission Tomography Tracer for Imaging Synucleinopathies. Cell 2023, 186 (16), 3350-3367.e19. 10.1016/j.cell.2023.06.004.

(53) Sanna, E.; Rodrigues, M.; Fagan, S. G.; Chisholm, T. S.; Kulenkampff, K.; Klenerman, D.; Spillantini, M. G.; Aigbirhio, F. I.; Hunter, C. A. Mapping the Binding Site Topology of Amyloid Protein Aggregates Using Multivalent Ligands. Chem. Sci. 2021, 12 (25), 8892–8899. 10.1039/d1sc01263k.

(54) De Greef, T. F. A.; Smulders, M. M. J.; Wolffs, M.; Schenning, A. P. H. J.; Sijbesma, R. P.; Meijer, E. W. Supramolecular Polymerization. Chem. Rev. 2009, 109 (11), 5687– 5754. 10.1021/cr900181u.

(55) Chen, Z.; Lohr, A.; Saha-Möller, C. R.; Würthner, F. Self-Assembled π-Stacks of Functional Dyes in Solution: Structural and Thermodynamic Features. Chem. Soc. Rev. 2009, 38 (2), 564–584. 10.1039/B809359H.

(56) Scott Lokey, R.; Iverson, B. L. Synthetic Molecules That Fold into a Pleated Secondary Structure in Solution. Nature 1995, 375 (6529), 303–305. 10.1038/375303a0.

(57) Sulatskaya, A. I.; Lavysh, A. V.; Maskevich, A. A.; Kuznetsova, I. M.; Turoverov, K. K. Thioflavin T Fluoresces as Excimer in Highly Concentrated Aqueous Solutions and as Monomer Being Incorporated in Amyloid Fibrils. Sci. Rep. 2017, 7 (1), 2146. 10.1038/s41598-017-02237-7.

(58) Choudhury, P.; Sarkar, S.; Das, P. K. Tunable Aggregation-Induced Multicolor Emission of Organic Nanoparticles by Varying the Substituent in Naphthalene Diimide. Langmuir 2018, 34 (47), 14328–14341. 10.1021/acs.langmuir.8b02996.

