## Supplementary material for "Two Distinct Binding Modes Govern High-Affinity Ligand Interactions with Amyloid Fibrils": Model - Main Fitter.ipynb

```
# %% [code]
import pandas as pd
import numpy as np
import math
import matplotlib.pyplot as plt
import matplotlib.cm as cm
import matplotlib.colors as mcolors
from scipy.optimize import least_squares
from scipy.optimize import fsolve
from scipy.optimize import minimize
from scipy.optimize import root_scalar
from operator import itemgetter
from typing import Callable, Dict, Optional, List, Sequence, Tuple, Iterable

# %% [markdown]
### Ligand Binding to a 1D Lattice

This code models the binding of ligands to a 1D lattice, accounting for ligand cooperativity, ligand size, and the possibility of a bound ligand creating a new "secondary" binding site.

This code is based on the McGhee-von Hippel model, and an extended version by Villaluenga et al. (DOI: 10.1103/PhysRevE.107.024401).

There are three main sections of code, corresponding to different models.

(1) equations_2lig_multisite() models binding for 2 ligands to a variable number of sites
(2) equations_1lig_multisite() models binding for 1 ligand to a variable number of sites
(3) equations_1lig_multisite_with_secondary() models binding for 1 ligand to a variable number of sites, accounting for the creation of a secondary site when a ligand binds to a "primary" site

Note - there are several common reasons why fits fail, or the generation of synthetic data produces incorrect or volatile results. Areas in the code prone to cause these errors are annotated with "Note". In brief, poor modelling is generally obtained when (a) input parameters differ by several orders of magnitude, (b) ligand total concentration begins very small, or ends very high (c) non-sensible initial guesses are made.

General variables are:
- n_i: length of ligand i in lattice units
- omega_ij: cooperativity parameter for ligand i binding directly adjacent to ligand l
- v_i: average number of bound ligand i per lattice unit
- theta_i: fractional lattice coverage by ligand i
- s_i: the average number of binding sites for ligand i per gap
- p_i: conditional probability of ligand i following a randomly selected free site
- q_i: conditional probability of a free lattice site following a randomly selected ligand i
- ff: conditional probability of a free lattice site following a randomly selected free lattice site
- L_i_free: concentration of unbound ligand i
- L_i_tot: total concentration of ligand i
- res_tot: total concentration of lattice residues
- res_free: concentration of unbound lattice residues
- bound_i: concentration of bound ligand


# %% [code]
# --- Setting up general required functions --- #

# Settings up a global epsilon to prevent division by very small numbers/et. Note - the size of GLOBAL_EPS can have a significant effect on
#     fitting/model success if operating near edge cases. This can be useful to modify if sensible results aren't obtained
GLOBAL_EPS = 1e-14

def _as_list(x, T):
    """Broadcast scalar or sequence to length T (site classes)."""
    if isinstance(x, (list, tuple, np.ndarray)):
        assert len(x) == T
        return list(x)
    return [x] * T

# --- Calculates the average number of binding sites per lattice unit for a single-ligand model --- #
def calculate_1_s_x(n_A, v_A, p_A, q_A, omega_AA, ff):
    """
    Average number of binding sites per lattice unit for a single-ligand model.   

    Returns
    -------
    s_A : float
        Av. binding site density  used in the mass-action equations for a 1-ligand model.
    """
    eps = GLOBAL_EPS
    denom_cov = max(n_A * v_A, eps)
    ff_ratio = ff / max(1 - ff, eps)

    sum_left_A  = (n_A * v_A / denom_cov) * q_A * (omega_AA + ff_ratio)
    sum_right_A = p_A * (omega_AA + ff_ratio)

    s_A = sum_left_A * ff ** (n_A - 1) * sum_right_A
    return s_A

# --- Calculates the average number of binding sites per lattice unit for a two-ligand model --- #
def calculate_2_s_x(n_A, v_A, n_B, v_B, p_A, p_B, q_A, q_B,
                  omega_AA, omega_AB, omega_BA, omega_BB, ff):
    """
    Average number of binding sites per lattice unit for a two-ligand model.

    Returns
    -------
    s_A, s_B : float
         Av. binding site density  used in the mass-action equations for a 2-ligand model.
    """
    eps = GLOBAL_EPS
    denom_cov = max(n_A * v_A + n_B * v_B, eps)
    ff_ratio = ff / max(1 - ff, eps)

    sum_left_A  = (n_A * v_A / denom_cov) * q_A * (omega_AA + ff_ratio) + (n_B * v_B / denom_cov) * q_B * (omega_BA + ff_ratio)
    sum_right_A = p_A * (omega_AA + ff_ratio) + p_B * (omega_AB + ff_ratio)

    sum_left_B  = (n_A * v_A / denom_cov) * q_A * (omega_AB + ff_ratio) + (n_B * v_B / denom_cov) * q_B * (omega_BB + ff_ratio)
    sum_right_B = p_A * (omega_BA + ff_ratio) + p_B * (omega_BB + ff_ratio)

    s_A = sum_left_A * ff ** (n_A - 1) * sum_right_A
    s_B = sum_left_B * ff ** (n_B - 1) * sum_right_B
    return s_A, s_B

# --- Calculates the weighted totals for each ligand across multiple binding site classes - useful for plotting --- #
def totals_from_sites(df: pd.DataFrame) -> pd.DataFrame:
    """
    Size-weighted totals across site classes (weights = res_tot).

    Returns totals vs L_A_tot:
      theta_A_tot, theta_B_tot, v_A_tot, v_B_tot and free ligand concs (first per L_A_tot).
    """
    total_res = df.drop_duplicates("site")["res_tot"].sum()
    wcols = ["theta_A", "theta_B", "v_A", "v_B"]
    dfw = df.assign(**{f"{c}_w": df[c] * df["res_tot"] for c in wcols})
    tot = (dfw.groupby("L_A_tot", as_index=False)
              .agg(**{f"{c}_w": (f"{c}_w", "sum") for c in wcols},
                   L_A_free=("L_A_free", "first"), L_B_free=("L_B_free", "first")))
    for c in wcols:
        tot[f"{c}_tot"] = tot[f"{c}_w"] / total_res
    return tot


# %% [code]
### ------- Two ligand, multi-site code -------- ###
# This code is for modelling the binding of two ligands to multiple different types of binding sites

# ---- Equations for 2 ligands across multiple site classes ----
def equations_2lig_multisite(vars_flat, params):
    """
    Residuals for the 2-ligand, multi-site model.

    Unknowns per site class i (block size 7):
      [v_A_i, v_B_i, p_A_i, p_B_i, q_A_i, q_B_i, ff_i]
    plus global free ligand pools:
      [L_A_free, L_B_free]

    params keys:
      - site_params (list of dicts): n_A_i, res_tot, K_A, K_B, omega_**
      - n_A, n_B
      - L_A_tot, L_B_tot

    Returns:
        List of residuals: [eq1_i, eq2_i, eq3_i, eq5_i for each site] + [eq4_total]
    """
    site_params = params["site_params"]
    T = len(site_params)
    L_A_tot, L_B_tot = params["L_A_tot"], params["L_B_tot"]

    v_A = vars_flat[0*T:1*T]; v_B = vars_flat[1*T:2*T]
    p_A = vars_flat[2*T:3*T]; p_B = vars_flat[3*T:4*T]
    q_A = vars_flat[4*T:5*T]; q_B = vars_flat[5*T:6*T]
    ff  = vars_flat[6*T:7*T]
    L_A_free, L_B_free = vars_flat[7*T:7*T+2]

    res, total_bound_A, total_bound_B = [], 0.0, 0.0
    eps = GLOBAL_EPS

    for i, spi in enumerate(site_params):
        # extracting parameters
        res_tot_i = spi["res_tot"]
        n_A, n_B = spi["n_A"], spi["n_B"]
        K_A_i, K_B_i = spi["K_A"], spi["K_B"]
        omega_AA = spi.get("omega_AA", 1.0); omega_AB = spi.get("omega_AB", 1.0)
        omega_BA = spi.get("omega_BA", 1.0); omega_BB = spi.get("omega_BB", 1.0)
        vAi, vBi, pAi, pBi, qAi, qBi, ffi = v_A[i], v_B[i], p_A[i], p_B[i], q_A[i], q_B[i], ff[i]

        # calculating parameters
        theta_i   = n_A * vAi + n_B * vBi
        bound_A_i = vAi * res_tot_i
        bound_B_i = vBi * res_tot_i
        res_free_i = res_tot_i * (1 - theta_i)

        # calculating binding site concentration
        s_A_i, s_B_i = calculate_2_s_x(n_A, vAi, n_B, vBi, pAi, pBi, qAi, qBi, omega_AA, omega_AB, omega_BA, omega_BB, ffi)

        # E1–E2: normalization around free states per site class
        eq1 = ffi + pAi + pBi - 1
        eq2 = ffi * (1 - theta_i) + vAi * qAi + vBi * qBi - (1 - theta_i)

        # E3–E4: implementing cooperativity constraints for adjacent/non-adjacent conditional probabilities
        eq3 = qAi + omega_AA * qAi * pAi / max(ffi, eps) + omega_AB * qAi * pBi / max(ffi, eps) - 1
        eq4 = qBi + omega_BA * qBi * pAi / max(ffi, eps) + omega_BB * qBi * pBi / max(ffi, eps) - 1

        # E5–E6: mass action equations per site class
        eq5 = bound_A_i / res_tot_i - K_A_i * s_A_i * L_A_free * (bound_A_i + bound_B_i) / res_tot_i
        eq6 = bound_B_i / res_tot_i - K_B_i * s_B_i * L_B_free * (bound_A_i + bound_B_i) / res_tot_i

        # E9–E10: linking conditional probabilities to species concentration
        eq9  = pAi * res_free_i / res_tot_i + (omega_AA * qAi * pAi / max(ffi, eps) - 1) * bound_A_i / res_tot_i + (omega_BA * qBi * pAi / max(ffi, eps)) * bound_B_i / res_tot_i
        eq10 = pBi * res_free_i / res_tot_i + (omega_BB * qBi * pBi / max(ffi, eps) - 1) * bound_B_i / res_tot_i + (omega_AB * qAi * pBi / max(ffi, eps)) * bound_A_i / res_tot_i

        res.extend([eq1, eq2, eq3, eq4, eq5, eq6, eq9, eq10])
        total_bound_A += bound_A_i
        total_bound_B += bound_B_i

    # Global ligand conservation 
    res.extend([(L_A_free + total_bound_A - L_A_tot)/res_tot_i, (L_B_free + total_bound_B - L_B_tot)/res_tot_i])
    return res

# ---- Fitting the 2 ligand model across multiple site classes ----
def fit_2lig_model(site_params, L_A_tot_values, L_B_tot, prev=None):
    T, eps = len(site_params), GLOBAL_EPS
    out, prev_sol = [], prev

    # Getting values for n_A, n_B values for warm start
    n_A = site_params[0]['n_A']
    n_B = site_params[0]['n_B']

    # setting up bounds for fitting/modelling
    ### Note - the bounds here can have a significant impact on fitting/modelling success. If the generated data shows irregularities or fails to 
    #     produce sensible outputs, ensure that these bounds are sensible for your situation
    lb = [0]*T + [0]*T + [eps]*T + [eps]*T + [eps]*T + [eps]*T + [eps]*T + [0, 0]
    ub = [1/n_A]*T + [1/n_B]*T + [1]*T + [1]*T + [1]*T + [1]*T + [1]*T + [max(L_A_tot_values), max(L_B_tot, eps)]

    # hot-starting off a previous solution, if available
    for L_A_tot in L_A_tot_values:
        if prev_sol is None:
            x0 = ([1/max(n_A+1, 2)] * T + [1/max(n_B+1, 2)] * T +
                  [0.1] * T + [0.1] * T + [0.9] * T + [0.9] * T + [0.5] * T +
                  [0.5 * L_A_tot, 0.5 * max(L_B_tot, eps)])
        else:
            x0 = prev_sol
        
        x0a = np.asarray(x0, dtype=float)
        lba = np.asarray(lb, dtype=float)
        uba = np.asarray(ub, dtype=float)

        # checking if bounds are sensible. Note - for strict bounds and if parameters are used with large order-of-magnitude differences, the code can get
        #    trapped at edge cases. If you're hitting this error try to normalise data, or reduce order-of-magnitude differences in input if possible
        bad = np.where((x0a < lba) | (x0a > uba) | ~np.isfinite(x0a))[0]
        if bad.size:
            raise ValueError(
                f"x0 infeasible at idx {bad[:10]}: x0={x0a[bad[:10]]}, lb={lba[bad[:10]]}, ub={uba[bad[:10]]}"
            )

        # solving
        sol = least_squares(
            equations_2lig_multisite, x0,
            args=({"site_params": site_params, "L_A_tot": L_A_tot, "L_B_tot": L_B_tot},),
            bounds=(lb, ub), method="trf", xtol=1e-10, ftol=1e-10, diff_step=1e-6, jac="3-point"
        ).x
        prev_sol = sol

        v_A = sol[0*T:1*T]; v_B = sol[1*T:2*T]
        p_A = sol[2*T:3*T]; p_B = sol[3*T:4*T]
        q_A = sol[4*T:5*T]; q_B = sol[5*T:6*T]
        ff  = sol[6*T:7*T]
        L_A_free, L_B_free = sol[7*T:7*T+2]

        # outputting useful values for visualisation and analysis
        for i, spi in enumerate(site_params):
            res_tot_i = spi["res_tot"]
            out.append(dict(
                site=f"Z{i+1}", L_A_tot=L_A_tot, L_B_tot=L_B_tot,
                v_A=v_A[i], v_B=v_B[i], theta_A=n_A*v_A[i], theta_B=n_B*v_B[i],
                p_A=p_A[i], p_B=p_B[i], q_A=q_A[i], q_B=q_B[i], ff=ff[i],
                L_A_free=L_A_free, L_B_free=L_B_free,
                bound_A=v_A[i]*res_tot_i, bound_B=v_B[i]*res_tot_i,
                res_tot=res_tot_i
            ))
    return pd.DataFrame(out)

# %% [code]
### ------- One ligand, multi-site code -------- ###
# This code is for modelling the binding of one ligand to multiple different types of binding sites
# This is based off the two ligand code, hence retention of the *_A notation - faster and more reliable fitting is obtained using the below code

# ---- Equations for 1 ligand across multiple site classes ----
def equations_1lig_multisite(vars_flat, params):
    """
    Residuals for the 1-ligand model.

    Unknowns per site class i (block size 4):
      [v_A_i, p_A_i, q_A_i, ff_i]
    plus global free pool:
      [L_A_free]

    params keys:
      - site_params (list of dicts): n_A, res_tot, K_A, omega_AA
      - L_A_tot
    """
    site_params = params["site_params"]
    T = len(site_params)    
    L_A_tot = params["L_A_tot"]

    v_A = vars_flat[0*T:1*T]
    p_A = vars_flat[1*T:2*T]
    q_A = vars_flat[2*T:3*T]
    ff  = vars_flat[3*T:4*T]
    L_A_free = vars_flat[4*T]

    res, total_bound_A = [], 0.0
    eps = GLOBAL_EPS

    for i, spi in enumerate(site_params):
        # extracting parameters
        res_tot_i = spi["res_tot"]
        n_A = spi["n_A"]
        K_A_i = spi["K_A"]
        omega_AA = spi.get("omega_AA", 1.0)
        vAi, pAi, qAi, ffi = v_A[i], p_A[i], q_A[i], ff[i]
        
        # calculating parameters
        theta_i = n_A * vAi
        bound_A_i = vAi * res_tot_i
        res_free_i = res_tot_i * (1 - theta_i)

        # calculating binding site concentration
        s_A_i = calculate_1_s_x(n_A, vAi, pAi, qAi, omega_AA, ffi)

        # E1–E2: normalization around free states per site class
        eq1 = ffi + pAi - 1
        eq2 = ffi * (1 - theta_i) + vAi * qAi - (1 - theta_i)

        # E3: implementing cooperativity constraints for adjacent/non-adjacent conditional probabilities
        eq3 = qAi + omega_AA * qAi * pAi / max(ffi, eps) - 1

        # E4: mass action equations per site class
        eq4 = 1 - K_A_i * s_A_i * L_A_free

        # E5: linking conditional probabilities to species concentration
        eq5 = pAi * res_free_i / res_tot_i + (omega_AA * qAi * pAi / max(ffi, eps) - 1) * bound_A_i / res_tot_i

        res.extend([eq1, eq2, eq3, eq4, eq5])
        total_bound_A += bound_A_i

    # Global ligand conservation (A only)
    res.append((L_A_free + total_bound_A - L_A_tot) / res_tot_i)
    return res

# ---- Fitting the 1 ligand model across multiple site classes ----
def fit_1lig_model(site_params, L_A_tot_values, L_B_tot, prev=None):
    
    T, eps = len(site_params), GLOBAL_EPS
    out, prev_sol = [], prev

    n_A = site_params[0]['n_A']

    # setting up bounds for fitting/modelling
    ### Note - the bounds here can have a significant impact on fitting/modelling success. If the generated data shows irregularities or fails to 
    #     produce sensible outputs, ensure that these bounds are sensible for your situation    
    lb = [0]*T + [eps]*T + [eps]*T + [eps]*T + [0]
    ub = [1/n_A]*T + [1]*T + [1]*T + [1]*T + [max(L_A_tot_values)]

    # hot-starting off a previous solution, if available
    for L_A_tot in L_A_tot_values:
        if prev_sol is None:
            x0 = ([1/max(n_A+1, 2)] * T + [0.1] * T + [0.9] * T + [0.5] * T + [0.5 * L_A_tot])
        else:
            x0 = prev_sol

        # solving
        sol = least_squares(
            equations_1lig_multisite, x0,
            args=({"site_params": site_params, "L_A_tot": L_A_tot},),
            bounds=(lb, ub), method="trf", xtol=1e-10, ftol=1e-10, diff_step=1e-6, jac="3-point"
        ).x
        prev_sol = sol

        v_A = sol[0*T:1*T]
        p_A = sol[1*T:2*T]
        q_A = sol[2*T:3*T]
        ff  = sol[3*T:4*T]
        L_A_free = sol[4*T]

        # outputting useful values for visualisation and analysis - inputting empty *_B values to harmonise downstream code
        for i, spi in enumerate(site_params):
            res_tot_i = spi["res_tot"]
            n_A = spi['n_A']
            out.append(dict(
                site=f"Z{i+1}", L_A_tot=L_A_tot, L_B_tot=0.0,
                v_A=v_A[i], v_B=0.0, theta_A=n_A*v_A[i], theta_B=0.0,
                p_A=p_A[i], p_B=0.0, q_A=q_A[i], q_B=0.0, ff=ff[i],
                L_A_free=L_A_free, L_B_free=0.0,
                bound_A=v_A[i]*res_tot_i, bound_B=0.0,
                res_tot=res_tot_i
            ))
    return pd.DataFrame(out)

# %% [code]
### ------- One ligand, new-site code -------- ###
# This code is for modelling the binding of one ligand where, upon binding, a new site is created

def _secondary_slots_active_per_bound(S1_total: float, L1_bound: float) -> float:
    """
    Active secondary slots for a site class:
      - one secondary slot per primary-bound ligand on that class
      - cannot exceed the number of primary slots
    Secondary has zero footprint here, so no lattice packing cap is required.
    """
    return min(L1_bound, S1_total)

# ---- Core: equations for 1 ligand across multiple site classes, where ligand binding produces a new site ----
def equations_1lig_multisite_with_secondary(vars_flat, params):
    """
    1-ligand multisite model with induced secondary binding (one secondary slot per primary-bound ligand).

    vars_flat layout:
      v_A[0:T], p_A[0:T], q_A[0:T], ff[0:T], y2[0:T], L_A_free

    params dict keys:
      site_params: list of dicts with res_tot, K_A (or K1), omega_AA, K2
      n_A: global footprint
      L_A_tot: total ligand concentration
    """
    site_params = params["site_params"]
    T = len(site_params)
    L_A_tot = params["L_A_tot"]
    eps = GLOBAL_EPS

    v_A = vars_flat[0*T:1*T]
    p_A = vars_flat[1*T:2*T]
    q_A = vars_flat[2*T:3*T]
    ff  = vars_flat[3*T:4*T]
    y2  = vars_flat[4*T:5*T]
    L_A_free = vars_flat[5*T]

    res, total_bound_1, total_bound_2 = [], 0.0, 0.0

    for i, spi in enumerate(site_params):
        # extracting parameters
        res_tot_i = float(spi["res_tot"])
        n_A = spi["n_A"]
        K1_i = float(spi.get("K1", spi.get("K_A")))
        omega_i = float(spi.get("omega_AA", 1.0))
        K2_i = float(spi.get("K2", 0.0))
        v, p, q, ffi, y = float(v_A[i]), float(p_A[i]), float(q_A[i]), float(ff[i]), float(y2[i])

        # calculating parameters
        theta = n_A * v
        bound1 = v * res_tot_i
        res_free = res_tot_i * (1.0 - theta)

        # calculating binding site concentration
        s_A = calculate_1_s_x(n_A, v, p, q, omega_i, ffi)

        # E1-E2: normalization around free states per site class
        eq1 = ffi + v * q / (1.0 - theta) - 1.0
        eq2 = q + omega_i * q * p / max(ffi, eps) - 1.0

        # Primary mass action equation
        eq3 = 1 - K1_i * s_A * L_A_free

        # implementing cooperativity constraints for adjacent/non-adjacent conditional probabilities
        eq4 = p * (1.0 - theta) + (omega_i * q * p / max(ffi, eps) - 1.0) * v

        res.extend([eq1, eq2, eq3, eq4])
        total_bound_1 += bound1

        # Secondary mass action equation: Langmuir on induced sites
        if K2_i > 0:
            S1_total = res_tot_i / n_A
            S2_active = _secondary_slots_active_per_bound(S1_total, bound1)
            bound2 = y * S2_active
            total_bound_2 += bound2
            eq_sec = y - (K2_i * L_A_free) / (1.0 + K2_i * L_A_free)
            res.append(eq_sec)
        else:
            # keep y2 pinned if K2 is off
            res.append(y)

    # Global mass balance
    res.append(L_A_free + total_bound_1 + total_bound_2 - L_A_tot)
    return res

    # ---- Fitting the 1 ligand-new site model across multiple site classes ----
def fit_1lig_newsite_model(site_params, L_A_tot_values, L_B_tot, prev=None):
    
    T, eps = len(site_params), GLOBAL_EPS
    out, prev_sol = [], prev
    n_A = site_params[0]['n_A']
    
    # setting up bounds for fitting/modelling
    ### Note - the bounds here can have a significant impact on fitting/modelling success. If the generated data shows irregularities or fails to 
    #     produce sensible outputs, ensure that these bounds are sensible for your situation
    lb = [0]*T + [eps]*T + [eps]*T + [eps]*T + [0]*T + [0]
    ub = [1/n_A]*T + [1]*T + [1]*T + [1]*T + [1]*T + [max(L_A_tot_values)]
    
    out, prev_sol = [], prev

    # hot-starting off a previous solution, if available
    for L_A_tot in L_A_tot_values:
        x0 = (prev_sol if prev_sol is not None else
              ([1/max(n_A+1, 2)]*T + [0.1]*T + [0.9]*T + [0.5]*T + [0.05]*T + [0.5*L_A_tot]))

        # solving
        sol = least_squares(
            equations_1lig_multisite_with_secondary, x0,
            args=({"site_params": site_params, "n_A": n_A, "L_A_tot": L_A_tot},),
            bounds=(lb, ub), method="trf", xtol=1e-10, ftol=1e-10, diff_step=1e-6, jac="3-point"
        ).x
        prev_sol = sol

        vA = sol[0*T:1*T]
        pA = sol[1*T:2*T]
        qA = sol[2*T:3*T]
        ffA = sol[3*T:4*T]
        y2 = sol[4*T:5*T]
        L_A_free = sol[5*T]

        for i, sp in enumerate(site_params):
            res_tot_i = sp["res_tot"]
            bound1 = vA[i] * res_tot_i
            theta1 = n_A * vA[i]
            S1_total = res_tot_i / n_A
            S2_active = _secondary_slots_active_per_bound(S1_total, bound1)
            bound2 = y2[i] * S2_active

            v_obs = (bound1 + bound2) / res_tot_i  # for Scatchard plot

            # outputting useful values for visualisation and analysis
            out.append(dict(
                site=f"Z{i+1}", L_A_tot=L_A_tot, L_B_tot=0.0,
                v_A=v_obs, v_A_primary=vA[i], theta_A=theta1,
                p_A=pA[i], q_A=qA[i], ff=ffA[i],
                y2=y2[i], S2_active=S2_active, bound_1=bound1, bound_2=bound2,
                L_A_free=L_A_free, L_B_free=0.0,
                bound_A=bound1 + bound2, bound_B=0.0,
                res_tot=res_tot_i,
                v_B=0.0, theta_B=0.0, p_B=0.0, q_B=0.0
            ))

    return pd.DataFrame(out)

# %% [code]
# ------- Main code --------

def fitter_main(site_params, L_A_tot_values, L_B_tot,
                           prev=None, include_B: bool = True, new_site: bool = False):
    """
    Solve either:
      - 2-ligand model: include_B=True
      - 1-ligand model: include_B=False
      - 1-ligand new-site model: include B=False, new_site=True

    Returns
    -------
    df : pd.DataFrame
        For include_B=False, B-related columns are still present (set to 0)
        so downstream code (totals/plotting) can remain unchanged.
    """


    # ---------------- 2-ligand model ----------------
    if include_B:
        if new_site:
            print('New site model cannot be used with multiple ligands. For this calculation, new_site is set to False')
            new_site = False
        return fit_2lig_model(site_params, L_A_tot_values, L_B_tot, prev=None)
        
    # ---------------- 1-ligand model with new site ----------------
    if new_site:
        return fit_1lig_newsite_model(site_params, L_A_tot_values, L_B_tot, prev=None)

    # ---------------- 1-ligand model ----------------
    return fit_1lig_model(site_params, L_A_tot_values, L_B_tot, prev=None)

# %% [code]
# ------- Plotting code --------
# this code generates binding isotherms, Scatchard plots, and Hill plots as required

def plot_isotherms(df: pd.DataFrame, *, include_B: bool = True, per_site=False, vmin_sc=0.01, eps=1e-12, fontsize=13):
    """
    Parameters
    ----------
    df : DataFrame
        Output of fitting functions to pllot.
    include_B : bool
        If False, treat as a 1-ligand run: plot only A-series panels.
        If True, plot both A and B totals and per-site (current behaviour).
    """

    # helper functions
    def _arr(x): return np.asarray(x, dtype=float)

    def hill_xy(L_free, theta):
        L, th = _arr(L_free), _arr(theta)
        return np.log10(L), np.log10(th / (1.0 - th))

    def sc_xy(v, L_free):
        v, L = _arr(v), _arr(L_free)
        return v, v / L

    tot = totals_from_sites(df)

    # masks (A always - ensure only sensible data is plotted)
    mA_h = (tot["L_A_free"] > 0) & tot["theta_A_tot"].between(eps, 1 - eps)
    mA_s = (tot["L_A_free"] > 0) & (tot["v_A_tot"] >= vmin_sc)

    # only if a second legand is present
    # ligand-specific, site-specific, and total values are plotted
    if include_B:
        # plot masks
        mB_h = (tot["L_B_free"] > 0) & tot["theta_B_tot"].between(eps, 1 - eps)
        mB_s = (tot["L_B_free"] > 0) & (tot["v_B_tot"] >= vmin_sc)

        plt.rcParams.update({"font.size": fontsize})
        fig, axs = plt.subplots(2, 3, figsize=(14, 8))

        # 1) Coverage vs [A] (per site + totals)
        ax = axs[0, 0]
        for site, g in df.groupby("site"):
            ax.plot(_arr(g["L_A_tot"]), _arr(g["theta_A"]), alpha=0.6, label=f"{site} theta_A")
            ax.plot(_arr(g["L_A_tot"]), _arr(g["theta_B"]), alpha=0.6, ls="--", label=f"{site} theta_B")
        ax.plot(_arr(tot["L_A_tot"]), _arr(tot["theta_A_tot"]), color="k", lw=2, label="Total θ_A")
        ax.plot(_arr(tot["L_A_tot"]), _arr(tot["theta_B_tot"]), color="k", lw=2, ls="--", label="Total θ_B")
        ax.set(xscale="log", xlabel="[A] (M)", ylabel="Coverage θ", title="Coverage vs [A]")
        ax.legend(ncol=2, fontsize=10)

        # 2) Hill (totals)
        ax = axs[0, 1]
        ax.plot(*hill_xy(tot.loc[mA_h, "L_A_free"], tot.loc[mA_h, "theta_A_tot"]), label="Total A")
        ax.plot(*hill_xy(tot.loc[mB_h, "L_B_free"], tot.loc[mB_h, "theta_B_tot"]), label="Total B")
        ax.set(xlabel="log(L_free)", ylabel="log(θ / (1-θ))", title="Hill (totals)")
        ax.legend()

        # 3) Scatchard (totals)
        ax = axs[0, 2]
        ax.plot(*sc_xy(tot.loc[mA_s, "v_A_tot"], tot.loc[mA_s, "L_A_free"]), label="Total A")
        ax.plot(*sc_xy(tot.loc[mB_s, "v_B_tot"], tot.loc[mB_s, "L_B_free"]), label="Total B")
        ax.set(xlabel="v", ylabel="v / L_free", title="Scatchard (totals)")
        ax.legend()

        # 4) Per-site Hill (A)
        ax = axs[1, 0]
        for site, g in df.groupby("site"):
            m = (g["L_A_free"] > 0) & g["theta_A"].between(eps, 1 - eps)
            ax.plot(*hill_xy(g.loc[m, "L_A_free"], g.loc[m, "theta_A"]), label=f"{site} A")
        ax.set(xlabel="log(L_A_free)", ylabel="log(θ_A / (1-θ_A))", title="Hill per site (A)")
        ax.legend(fontsize=10)

        # 5) Per-site Scatchard (A)
        ax = axs[1, 1]
        for site, g in df.groupby("site"):
            m = (g["L_A_free"] > 0) & (g["v_A"] >= vmin_sc)
            ax.plot(*sc_xy(g.loc[m, "v_A"], g.loc[m, "L_A_free"]), label=str(site))
        ax.set(xlabel="v_A", ylabel="v_A / L_A_free", title="Scatchard per site (A)")
        ax.legend(fontsize=10)

        # 6) Total coverage (A+B)
        ax = axs[1, 2]
        ax.plot(_arr(tot["L_A_tot"]), _arr(tot["theta_A_tot"] + tot["theta_B_tot"]), color="k", label="θ_total")
        ax.set(xscale="log", xlabel="[A] (M)", ylabel="θ_total", title="Total coverage")
        ax.legend()

        fig.tight_layout()
        return fig, axs

    # if a saturation assay
    # plot per-site and total information
    if not include_B:
        plt.rcParams.update({"font.size": fontsize})
        fig, axs = plt.subplots(2, 3, figsize=(14, 8)) if per_site else plt.subplots(1, 3, figsize=(14, 4))
        axs = np.atleast_2d(axs)

        # --- totals row (same as 1-ligand plots) ---
        ax = axs[0,0]
        for site, g in df.groupby("site"):
            ax.plot(_arr(g["L_A_tot"]), _arr(g["theta_A"]), alpha=0.6, label=f"{site} θ_A")
        ax.plot(_arr(tot["L_A_tot"]), _arr(tot["theta_A_tot"]), color="k", lw=2, label="Total θ_A")
        ax.set(xscale="log", xlabel="[A] (M)", ylabel="Coverage θ_A", title="Coverage vs [A] (A)")
        ax.legend(ncol=2, fontsize=10)

        ax = axs[0,1]
        ax.plot(*hill_xy(tot.loc[mA_h,"L_A_free"], tot.loc[mA_h,"theta_A_tot"]), label="Total A")
        ax.set(xlabel="log(L_A_free)", ylabel="log(θ_A/(1-θ_A))", title="Hill (total)")
        ax.legend()

        ax = axs[0,2]
        ax.plot(*sc_xy(tot.loc[mA_s,"v_A_tot"], tot.loc[mA_s,"L_A_free"]), label="Total A")
        ax.set(xlabel="v_A", ylabel="v_A / L_A_free", title="Scatchard (total)")
        ax.legend()

        if per_site:
            # --- per-site row ---
            ax = axs[1,0]
            for site, g in df.groupby("site"):
                ax.plot(_arr(g["L_A_tot"]), _arr(g["theta_A"]), alpha=0.6, label=str(site))
            ax.set(xscale="log", xlabel="[A] (M)", ylabel="θ_A", title="Coverage per site (A)")
            ax.legend(fontsize=10)

            ax = axs[1,1]
            for site, g in df.groupby("site"):
                m = (g["L_A_free"] > 0) & g["theta_A"].between(eps, 1-eps)
                ax.plot(*hill_xy(g.loc[m,"L_A_free"], g.loc[m,"theta_A"]), label=str(site))
            ax.set(xlabel="log(L_A_free)", ylabel="log(θ_A/(1-θ_A))", title="Hill per site (A)")
            ax.legend(fontsize=10)

            ax = axs[1,2]
            for site, g in df.groupby("site"):
                m = (g["L_A_free"] > 0) & (g["v_A"] >= vmin_sc)
                ax.plot(*sc_xy(g.loc[m,"v_A"], g.loc[m,"L_A_free"]), label=str(site))
            ax.set(xlabel="v_A", ylabel="v_A / L_A_free", title="Scatchard per site (A)")
            ax.legend(fontsize=10)

        fig.tight_layout()
        return fig, axs
    
    # -------- 1-ligand plotting (A only): single series of plots --------
    plt.rcParams.update({"font.size": fontsize})
    fig, axs = plt.subplots(1, 3, figsize=(14, 4))

    # 1) Coverage vs [A] (per site + total) for A only
    ax = axs[0]
    for site, g in df.groupby("site"):
        ax.plot(_arr(g["L_A_tot"]), _arr(g["theta_A"]), alpha=0.6, label=f"{site} θ_A")
    ax.plot(_arr(tot["L_A_tot"]), _arr(tot["theta_A_tot"]), color="k", lw=2, label="Total θ_A")
    ax.set(xscale="log", xlabel="[A] (M)", ylabel="Coverage θ_A", title="Coverage vs [A] (A only)")
    ax.legend(ncol=2, fontsize=10)

    # 2) Hill (totals) for A only
    ax = axs[1]
    ax.plot(*hill_xy(tot.loc[mA_h, "L_A_free"], tot.loc[mA_h, "theta_A_tot"]), label="Total A")
    ax.set(xlabel="log(L_A_free)", ylabel="log(θ_A / (1-θ_A))", title="Hill (A only)")
    ax.legend()

    # 3) Scatchard (totals) for A only
    ax = axs[2]
    ax.plot(*sc_xy(tot.loc[mA_s, "v_A_tot"], tot.loc[mA_s, "L_A_free"]), label="Total A")
    ax.set(xlabel="v_A", ylabel="v_A / L_A_free", title="Scatchard (A only)")
    ax.legend()

    fig.tight_layout()
    return fig, axs

# %% [code]
# --- Example: one-ligand, one-site system ---
site_params = [
    dict(n_A = 1, n_B = 1, res_tot=100, K_A=10, K_B=0, omega_AA=4, omega_AB=1, omega_BA=1, omega_BB=1)
]
L_A_tot_values = np.logspace(-1, 3, 60)  # sweep total [A]

# 1-ligand (A only)
df = fitter_main(site_params, L_A_tot_values, L_B_tot=0.0, include_B=False)
plot_isotherms(df, include_B=False, per_site=False); plt.show()

# %% [code]
# --- Example: one-ligand, two-site system ---
site_params = [
    dict(n_A = 4, n_B = 1, res_tot=100, K_A=1, K_B=0, omega_AA=1, omega_AB=1, omega_BA=1, omega_BB=1),
    dict(n_A = 4, n_B = 1, res_tot=100, K_A=1, K_B=0, omega_AA=1, omega_AB=1, omega_BA=1, omega_BB=1)
]
L_A_tot_values = np.logspace(0, 3, 60)  # sweep total [A]

# 1-ligand
df = fitter_main(site_params, L_A_tot_values, L_B_tot=0.0, include_B=False)
plot_isotherms(df, include_B=False); plt.show()

# %% [code]
# --- Example: one-ligand, new-site system ---
site_params = [
    dict(n_A = 4, n_B = 1, res_tot=100, K_A=10, omega_AA=1, K2=1)
]
L_A_tot_values = np.logspace(-1, 3, 80)  # sweep total [A]

# 1-ligand (A only)
df = fitter_main(site_params, L_A_tot_values, L_B_tot=0.0, include_B=False, new_site=True)
plot_isotherms(df, include_B=False, per_site = True); plt.show()

# %% [code]
# --- Example: two-ligand, one-site system ---
site_params = [
    dict(n_A = 4, n_B = 4, res_tot=100, K_A=1, K_B=10, omega_AA=1, omega_AB=1, omega_BA=1, omega_BB=1)
]
L_B_tot = 1  # fixed total [B] (units consistent with your fitting function)
L_A_tot_values = np.logspace(-3, 4, 80)  # sweep total [A]

# 2-ligand
df = fitter_main(site_params, L_A_tot_values, L_B_tot, include_B=True)
plot_isotherms(df, include_B=True); plt.show()

# %% [code]
# --- Example: two-ligand, two-site system ---
site_params = [
    dict(n_A = 1, n_B = 1, res_tot=100, K_A=10, K_B=1, omega_AA=4, omega_AB=1, omega_BA=1, omega_BB=1),
    dict(n_A = 1, n_B = 1, res_tot=100, K_A=1, K_B=10, omega_AA=1, omega_AB=1, omega_BA=1, omega_BB=1)
]
L_B_tot = 1  # fixed total [B] (units consistent with your fitting function)
L_A_tot_values = np.logspace(-3, 4, 80)  # sweep total [A]

# 2-ligand
df = fitter_main(site_params, L_A_tot_values, L_B_tot, include_B=True)
plot_isotherms(df, include_B=True); plt.show()

# %% [code]
```
