## Supplementary material for "Two Distinct Binding Modes Govern High-Affinity Ligand Interactions with Amyloid Fibrils": readme.txt

Two files containing code are provided that simulate (Model - Main Fitter) or fit (speciation_1Dbinding) 1D lattice binding models. These models implement cooperative multisite binding accounting for ligand size.

Model - Main Fitter.ipynb 	Model definitions used for generating synthetic data and modelling different isotherms
speciation_1Dbinding.py		Example fitting code containing new functions that can be implemented in Musketeer software

To run this code, open Model - Main Fitter.ipynb in Jupyter Notebook or equivalent, or use speciation_1Dbinding.py in place of speciation.py in Musketeer. Alternatively, the three 1D speciation functions can be added into an existing Musketeer speciation.py file.

Note that the developed models assume equilibrium, symmetric binding to a 1D lattice, and reversible binding.

The binding models can produce unreliable results in some scenarios, such as when large order-of-magnitude differences are used for stating parameters, and very small or large ligand concentrations are used. Errors are apparent in discontinuous or noisy output, and further details of at-risk areas are given in comments within the code.

All figures in the manuscript were generated using Model - Main Fitter.ipynb, with the following parameters in site_params:

Figure 3 (include_B=False)
- One standard 1:1 binding site (equivalent to non-cooperative stacked binding): dict(n_A = 1, n_B = 1, res_tot=100, K_A=1, K_B=0, omega_AA=1, omega_AB=1, omega_BA=1, omega_BB=1)
- One cooperative stacked binding site: dict(n_A = 1, n_B = 1, res_tot=100, K_A=1, K_B=0, omega_AA=4, omega_AB=1, omega_BA=1, omega_BB=1)
- One linear binding site: dict(n_A = 4, n_B = 1, res_tot=100, K_A=1, K_B=0, omega_AA=1, omega_AB=1, omega_BA=1, omega_BB=1)
- One linear new-site binding site: dict(n_A = 4, n_B = 1, res_tot=100, K_A=1, omega_AA=1, K2=1) and new_site=True

Figure 4 (include_B=True)
- Two standard 1:1 binding sites (equivalent to non-cooperative stacked binding): dict(n_A = 1, n_B = 1, res_tot=100, K_A=10, K_B=1, omega_AA=1, omega_AB=1, omega_BA=1, omega_BB=1)
- One cooperative stacked and one linear binding site: dict(n_A = 4, n_B = 1, res_tot=100, K_A=1, K_B=1, omega_AA=1, omega_AB=1, omega_BA=1, omega_BB=4)
- One standard and one cooperative stacked binding site: dict(n_A = 1, n_B = 1, res_tot=100, K_A=1, K_B=1, omega_AA=4, omega_AB=1, omega_BA=1, omega_BB=1)
- One standard and one linear binding site: dict(n_A = 4, n_B = 1, res_tot=100, K_A=1, K_B=1, omega_AA=1, omega_AB=1, omega_BA=1, omega_BB=4)
- Two cooperative stacked binding sites: dict(n_A = 1, n_B = 1, res_tot=100, K_A=10, K_B=1, omega_AA=4, omega_AB=1, omega_BA=1, omega_BB=4)
- Two linear binding sites: dict(n_A = 4, n_B = 1, res_tot=100, K_A=10, K_B=1, omega_AA=1, omega_AB=1, omega_BA=1, omega_BB=1)
