## Supplementary material for "Two Distinct Binding Modes Govern High-Affinity Ligand Interactions with Amyloid Fibrils": speciation_1Dbinding.py

```
import math
import tkinter.ttk as ttk
import warnings
from abc import abstractmethod

import numpy as np
from numpy import ma
import scipy
from scipy.optimize import least_squares, minimize

from . import moduleFrame
from .scrolledFrame import ScrolledFrame
from .style import padding
from .table import ButtonFrame, Table, WrappedLabel

LN_10 = np.log(10)


def stoichiometriesToBoundNames(freeNames, stoichiometries):
    trans = str.maketrans("0123456789", "₀₁₂₃₄₅₆₇₈₉")

    boundNames = []
    for row in stoichiometries:
        boundName = ""
        for freeName, stoichiometry in zip(freeNames, row):
            if stoichiometry == 0:
                continue

            if boundName != "":
                boundName += "·"

            if stoichiometry == 1:
                boundName += freeName
            elif stoichiometry == -1:
                boundName += freeName + "ₙ"
            else:
                boundName += freeName + str(stoichiometry).translate(trans)

        allNames = set(freeNames) | set(boundNames)
        if boundName in allNames:
            copyIndex = 2
            while boundName + f" ({copyIndex})" in allNames:
                copyIndex += 1
            boundName += f" ({copyIndex})"

        boundNames.append(boundName)
    return np.array(boundNames)


class ComplexSpeciationMixin:
    @property
    def complexIndices(self):
        return ~np.any(self.stoichiometries < 0, 1)

    @property
    def complexCount(self):
        return np.count_nonzero(self.complexIndices)

    @property
    def complexStoichiometries(self):
        return self.stoichiometries[self.complexIndices]

    @property
    def complexBoundNames(self):
        return stoichiometriesToBoundNames(self.freeNames, self.complexStoichiometries)

    def complexFormsBinaryComplex(self, i, j):
        if i == j:
            desiredRow = [2 if index == i else 0 for index in range(self.freeCount)]
        else:
            desiredRow = [
                1 if index in (i, j) else 0 for index in range(self.freeCount)
            ]
        return np.any(np.all(self.complexStoichiometries == desiredRow, axis=1))

    @property
    def complexMaxValencyPerGuest(self):
        # output[i, j] = the max number of js that can bind to one i
        output = np.zeros([self.freeCount, self.freeCount], dtype=int)
        for row in self.complexStoichiometries:
            if np.count_nonzero(row) == 1:
                host = guest = np.where(row > 0)[0][0]
                valency = 1 if row[host] == 1 else 2
            elif np.count_nonzero(row) == 2 and np.count_nonzero(row == 1) == 1:
                host = np.where(row == 1)[0][0]
                guest = np.where(row > 1)[0][0]
                valency = row[guest]
            elif np.count_nonzero(row) == 2 and np.count_nonzero(row == 1) == 2:
                host, guest = np.where(row == 1)[0]
                valency = 1
                output[host, guest] = max(output[host, guest], valency)
                output[guest, host] = max(output[guest, host], valency)
            else:
                continue
            output[host, guest] = max(output[host, guest], valency)

        return output

    def getComplexNoCooperativityValues(self):
        # Decomposes global Ks into products of microscopic Ks

        # Go through the complexes in order of increasing size, so that we can use
        # statistical factors of sub-complexes.
        sortedRowIndices = np.argsort(self.complexStoichiometries.sum(axis=1))
        M = self.complexStoichiometries[sortedRowIndices]

        statisticalFactors = np.ones(self.complexCount)
        ksMatrix = np.zeros(
            [self.complexCount, self.freeCount, self.freeCount], dtype=int
        )
        microKsNames = [
            [f"microK_{i}·{j}" for j in self.freeNames] for i in self.freeNames
        ]

        for rowIndex, row in enumerate(M):
            if np.count_nonzero(row) >= 1:
                statisticalFactor = 1
                for componentIndex in np.nonzero(row)[0]:
                    componentStatisticalFactor = 1
                    # find largest super-complex wrt this component, if it exists
                    for superRow in M[rowIndex + 1 : :][::-1]:
                        if (
                            np.count_nonzero(superRow) == np.count_nonzero(row)
                            and np.count_nonzero(superRow - row) == 1
                            and np.nonzero(superRow - row)[0].item() == componentIndex
                        ):
                            componentStatisticalFactor = math.comb(
                                superRow[componentIndex], row[componentIndex]
                            )
                            break
                    # find sub-complex lacking this component
                    targetSubRow = row.copy()
                    targetSubRow[componentIndex] = 0
                    for subRowIndex, subRow in enumerate(M[:rowIndex]):
                        if np.array_equal(subRow, targetSubRow):
                            componentStatisticalFactor *= statisticalFactors[
                                subRowIndex
                            ]
                            break

                    statisticalFactor = max(
                        statisticalFactor, componentStatisticalFactor
                    )
                statisticalFactors[rowIndex] = statisticalFactor

            # now calculate the exponents of the microscopic Ks
            for index, i in enumerate(np.nonzero(row)[0]):
                for j in np.nonzero(row)[0][index:]:
                    if not self.complexFormsBinaryComplex(i, j):
                        continue
                    if i == j:
                        # assume dimers form a single bond, while larger complexes are rings
                        if row[i] == 1:
                            continue
                        elif row[i] == 2:
                            ksMatrix[rowIndex, i, j] = 1
                        else:
                            ksMatrix[rowIndex, i, j] = row[i]
                    else:
                        maxIBonds = row[i] * self.complexMaxValencyPerGuest[i, j]
                        maxJBonds = row[j] * self.complexMaxValencyPerGuest[j, i]
                        maxTotalBonds = row[i] * row[j]
                        ksMatrix[rowIndex, i, j] = min(
                            maxIBonds, maxJBonds, maxTotalBonds
                        )

        # flatten i and j into a single index
        ksMatrix = ksMatrix.reshape(self.complexCount, self.freeCount**2)
        microKsNames = np.array(microKsNames).flatten()

        # filter out zero columns
        nonZeroColumns = np.any(ksMatrix != 0, axis=0)
        ksMatrix = ksMatrix[:, nonZeroColumns]
        microKsNames = microKsNames[nonZeroColumns]

        # unsort rows, and transpose Ks matrix to format expected by equilibriumConstants
        unsortedIndices = np.argsort(sortedRowIndices)
        return (
            statisticalFactors[unsortedIndices].copy(),
            ksMatrix[unsortedIndices].T.copy(),
            microKsNames,
        )

    complexVariableNames = complexOutputNames = complexBoundNames

    complexOutputStoichiometries = complexStoichiometries

    def complexFreeToBoundConcs(self, freeConcs, complexKs):
        return complexKs * np.prod(freeConcs**self.complexStoichiometries, axis=1)

    def complexObjective(self, free, complexKs, total, M):
        return complexKs @ np.prod(free**M, 1)

    def complexJacobian(self, free, complexKs, total, M):
        return M.T @ (complexKs * np.prod(free**M, 1))

    def complexHessian(self, free, complexKs, total, M):
        return M.T @ (M * np.outer(complexKs * np.prod(free**M, 1), 1 / free))

    def complexGetUpperBounds(self, complexKs, total, M):
        return total


class PolymerSpeciationMixin:
    @property
    def componentsThatFormPolymers(self):
        return np.any(self.stoichiometries < 0, axis=0)

    @property
    def polymerIndices(self):
        return np.any(self.stoichiometries < 0, axis=1)

    @property
    def polymerCount(self):
        return np.count_nonzero(self.polymerIndices)

    @property
    def polymerStoichiometries(self):
        return self.stoichiometries[self.polymerIndices]

    @property
    def polymerBoundNames(self):
        return stoichiometriesToBoundNames(self.freeNames, self.polymerStoichiometries)

    @property
    def polymerVariableNames(self):
        variableNames = []
        for row, boundName in zip(self.polymerStoichiometries, self.polymerBoundNames):
            if np.count_nonzero(row) == 1:
                variableNames.append(boundName.replace("ₙ", "₂"))
            variableNames.append(boundName)
        return np.array(variableNames)

    @property
    def polymerVariableCount(self):
        return len(self.polymerVariableNames)

    @property
    def polymerOutputNames(self):
        return np.ravel(
            [
                [name + " terminal", name + " internal"]
                for name in self.polymerBoundNames
            ]
        )

    @property
    def polymerOutputCount(self):
        return len(self.polymerOutputNames)

    @property
    def polymerOutputStoichiometries(self):
        # stoichiometries in terminal+internal mode: duplicate each polymer
        outputStoichiometries = np.empty([self.polymerOutputCount, self.freeCount])
        M = self.polymerStoichiometries
        outputStoichiometries[::2] = np.where(M > 0, M / 2, abs(M))  # terminal rows
        outputStoichiometries[1::2] = np.where(M > 0, 0, abs(M))  # internal rows
        return outputStoichiometries

    @property
    def polymerMaxValencyPerGuest(self):
        # output[i, j] = the max number of js that can bind to one i
        output = np.zeros([self.freeCount, self.freeCount], dtype=int)
        for row in self.polymerStoichiometries:
            if np.count_nonzero(row) == 1:
                host = guest = np.where(row < 0)[0][0]
                valency = 2
            else:
                continue
            output[host, guest] = max(output[host, guest], valency)

        return output

    def getPolymerNoCooperativityValues(self):
        statisticalFactors = np.ones(self.polymerVariableCount)
        ksMatrix = np.empty((0, self.polymerCount), dtype=int)
        microKsNames = [f"microK_{name}" for name in self.polymerBoundNames]

        for rowIndex, row in enumerate(self.polymerStoichiometries):
            outputRow = np.zeros(self.polymerCount, dtype=int)
            outputRow[rowIndex] = 1

            if np.count_nonzero(row) == 1:
                # K2 and Kn are identical, i.e. isodesmic polymerisation
                ksMatrix = np.vstack([ksMatrix, outputRow, outputRow])
            else:
                # for end-capped polymers, do not attempt to decompose the Kab
                ksMatrix = np.vstack([ksMatrix, outputRow])

        return statisticalFactors, ksMatrix.T, np.array(microKsNames)

    def splitPolymerKs(self, polymerKs):
        k2s = np.zeros(self.freeCount)
        kns = np.zeros(self.freeCount)
        kabs = np.ones(self.polymerCount)
        ks = list(polymerKs)
        for polymerIndex, row in enumerate(self.polymerStoichiometries):
            freeIndex = np.where(row < 0)[0][0]
            if np.count_nonzero(row) == 1:
                k2s[freeIndex], kns[freeIndex] = ks.pop(0), ks.pop(0)
            else:
                kabs[polymerIndex] = ks.pop(0)
        return k2s, kns, kabs

    def getTerminalInternalConcs(self, freeConcs, k2s, kns, kabs):
        terminal = 2 * freeConcs**2 * k2s / (1 - freeConcs * kns)
        internal = freeConcs**3 * k2s * kns / (1 - freeConcs * kns)
        return terminal, internal

    def polymerFreeToBoundConcs(self, freeConcs, k2s, kns, kabs):
        # Treat the terminal and internal units as additional "free" components, so
        # that end-capped polymers can be treated as if they're regular complexes.
        terminal = 2 * freeConcs**2 * k2s / (1 - freeConcs * kns)
        internal = freeConcs**3 * k2s * kns / (1 - freeConcs * kns) ** 2
        componentConcs = np.concatenate([freeConcs, terminal, internal])

        freeCount = self.freeCount
        polymerCount = self.polymerCount

        # split end group (pos) and polymer (neg) stoichiometries
        pos = np.where(self.polymerStoichiometries < 0, 0, self.polymerStoichiometries)
        neg = np.where(
            self.polymerStoichiometries < 0, np.abs(self.polymerStoichiometries), 0
        )

        # each polymer gives two outputs: terminal and internal
        fullStoichiometries = np.zeros([polymerCount * 2, freeCount * 3])

        # terminal and internal states both require end cap concentrations
        fullStoichiometries[::2, :freeCount] = pos
        fullStoichiometries[1::2, :freeCount] = pos
        fullStoichiometries[::2, freeCount : freeCount * 2] = neg
        fullStoichiometries[1::2, freeCount * 2 :] = neg
        return np.repeat(kabs, 2) * np.prod(componentConcs**fullStoichiometries, axis=1)

    def polymerFreeExactSolutionSingle(self, k2, kn, totalSingle):
        roots = np.roots(
            [
                k2 * kn - kn**2,
                totalSingle * kn**2 + 2 * kn - 2 * k2,
                -1 - 2 * totalSingle * kn,
                totalSingle,
            ]
        )
        real_roots = np.real(roots[np.isreal(roots)])
        return np.min(real_roots[real_roots > 0])

    def polymerFreeExactSolution(self, k2s, kns, total):
        return np.array(
            [
                self.polymerFreeExactSolutionSingle(k2, kn, totalSingle)
                for k2, kn, totalSingle in zip(k2s, kns, total)
            ]
        )

    def polymerObjective(self, free, k2s, kns, kabs, total, M):
        if self.polymerCount == 0:
            return 0.0
        pos = np.where(M < 0, 0, M)
        neg = np.where(M < 0, np.abs(M), 0)
        polymerWithoutFactorOfN = free**2 * k2s / (1 - free * kns)

        polymerIntegral = kabs @ np.prod(
            free**pos * polymerWithoutFactorOfN**neg, axis=1
        )
        return np.sum(polymerIntegral)

    def polymerJacobian(self, free, k2s, kns, kabs, total, M):
        if self.polymerCount == 0:
            return np.full(len(free), 0.0)

        pos = np.where(M < 0, 0, M)
        neg = np.where(M < 0, np.abs(M), 0)
        polymerWithFactorOfN = free**2 * k2s * (2 - free * kns) / (1 - free * kns) ** 2
        polymerWithoutFactorOfN = free**2 * k2s / (1 - free * kns)
        polymerConcentration = neg.T @ (
            kabs * np.prod(free**pos * polymerWithFactorOfN**neg, axis=1)
        )
        endCapConcentration = pos.T @ (
            kabs * np.prod(free**pos * polymerWithoutFactorOfN**neg, axis=1)
        )
        return polymerConcentration + endCapConcentration

    def polymerGetUpperBounds(self, k2s, kns, kabs, total, M):
        if not np.any(M < 0):
            return np.full(len(total), np.inf)
        componentsThatFormPolymers = np.any(M < 0, axis=0)
        return np.where(
            componentsThatFormPolymers,
            self.polymerFreeExactSolution(k2s, kns, total),
            np.inf,
        )


class Speciation(ComplexSpeciationMixin, PolymerSpeciationMixin, moduleFrame.Strategy):
    requiredAttributes = ("stoichiometries",)

    @property
    def stoichiometries(self):
        return self._stoichiometries

    @stoichiometries.setter
    def stoichiometries(self, value):
        self._stoichiometries = value
        self._noCooperativityValues = self.getNoCooperativityValues()

    @abstractmethod
    def run(self, variables, totalConcs):
        pass

    # Other modules need to access freeNames and freeCount, but totalConcentrations
    # may not yet be loaded.
    @property
    def freeNames(self):
        try:
            return self.titration.totalConcentrations.freeNames
        except AttributeError:
            return np.array(["Host", "Guest"])

    @property
    def freeCount(self):
        return len(self.freeNames)

    @property
    def boundNames(self):
        return np.append(self.complexBoundNames, self.polymerBoundNames)

    @property
    def boundCount(self):
        return len(self.boundNames)

    @property
    def variableNames(self):
        return np.concatenate([self.complexVariableNames, self.polymerVariableNames])

    @property
    def outputNames(self):
        return np.concatenate(
            [self.freeNames, self.complexOutputNames, self.polymerOutputNames]
        )

    @property
    def outputStoichiometries(self):
        return np.vstack(
            [
                np.eye(self.freeCount),
                self.complexOutputStoichiometries,
                self.polymerOutputStoichiometries,
            ]
        )

    @property
    def maximumValencyPerGuest(self):
        return np.maximum(
            self.complexMaxValencyPerGuest, self.polymerMaxValencyPerGuest
        )

    @property
    def noCooperativityValues(self):
        try:
            return self._noCooperativityValues
        except AttributeError:
            self._noCooperativityValues = self.getNoCooperativityValues()
            return self._noCooperativityValues

    def getNoCooperativityValues(self):
        statisticalFactorsC, ksMatrixC, microKsNamesC = (
            self.getComplexNoCooperativityValues()
        )
        statisticalFactorsP, ksMatrixP, microKsNamesP = (
            self.getPolymerNoCooperativityValues()
        )

        statisticalFactors = np.concatenate([statisticalFactorsC, statisticalFactorsP])
        ksMatrix = scipy.linalg.block_diag(ksMatrixC, ksMatrixP)
        microKsNames = np.concatenate([microKsNamesC, microKsNamesP])

        return statisticalFactors, ksMatrix, microKsNames

    def variablesToKs(self, variables):
        complexKs = variables[: self.complexCount]
        polymerKs = variables[self.complexCount :]
        k2s, kns, kabs = self.splitPolymerKs(polymerKs)
        return complexKs, k2s, kns, kabs

    # TODO: rewrite all non-mixin functions to be agnostic to the components of
    # polymerKs, by just working with complexKs and polymerKs, or possibly *polymerKs
    def freeToBoundConcs(self, freeConcs, complexKs, k2s, kns, kabs):
        return np.hstack(
            [
                self.complexFreeToBoundConcs(freeConcs, complexKs),
                self.polymerFreeToBoundConcs(freeConcs, k2s, kns, kabs),
            ]
        )


class SpeciationTable(Table):
    def __init__(self, master, titration):
        sortedTitleLengths = sorted(
            [len(name) for name in titration.totalConcentrations.freeNames]
        )
        self.width = max(6, sortedTitleLengths[-1] + 2)
        self.rowTitleWidth = min(20, sum(sortedTitleLengths[-2:]) + 3 + 2)

        super().__init__(
            master,
            0,
            0,
            titration.totalConcentrations.freeNames,
            rowOptions=("readonlyTitles", "new", "delete"),
            columnOptions=("readonlyTitles",),
            boldTitles=True,
            callback=self.updateTitles,
        )

        if (
            titration.speciation.stoichiometries.shape[1]
            == titration.totalConcentrations.freeCount
        ):
            for boundName, stoichiometry in zip(
                titration.speciation.boundNames, titration.speciation.stoichiometries
            ):
                stoichiometry = stoichiometry.astype(str)
                stoichiometry[stoichiometry == "-1"] = "n"
                self.addRow(boundName, stoichiometry)
        else:
            if titration.totalConcentrations.freeCount == 1:
                self.addRow("", np.array([1]))
            else:
                self.addRow(
                    "",
                    np.concatenate(
                        [
                            [1, 1],
                            [0] * (titration.totalConcentrations.freeCount - 2),
                        ]
                    ).astype(int),
                )
        self.updateTitles()

    def updateTitles(self, *args, **kwargs):
        for cell, title in zip(
            self.cells[2 : self.headerCells :, 1], self.columnTitles
        ):
            cell.set(title)

        self.rowTitles = stoichiometriesToBoundNames(self.columnTitles, self.data)

    def addFreeRow(self, index=-1):
        row = self.cells.shape[0]
        freeRow = np.full(self.cells.shape[1], None)

        freeRow[0] = self.deleteRowButton(row, 0, "Delete Row", state="disabled")
        freeRow[1] = self.readonlyEntry(
            row, 1, self.columnTitles[index], font=self.titleFont
        )

        freeRow[2:] = [
            self.readonlyEntry(row, 2 + column, "0", align="right")
            for column in range(self.cells.shape[1] - 2)
        ]
        freeRow[-1].set("1")

        self.cells = np.insert(self.cells, self.headerCells, freeRow, axis=0)
        self.headerCells += 1

        self.redraw()

    def addColumn(self, firstEntry="", data=None):
        super().addColumn(firstEntry, data)

        for row in range(2, self.headerCells):
            self.cells[row, -1] = self.readonlyEntry(
                row, self.cells.shape[-1], "0", align="right"
            )

        self.addFreeRow()

    def deleteColumn(self, column):
        super().deleteColumn(column)
        self.headerCells -= 1
        self.deleteRow(column)
        self.updateTitles()

    def convertData(self, number):
        if number == "n":
            return -1
        elif number == "":
            return 0
        else:
            return int(number)


class SpeciationPopup(moduleFrame.Popup):
    def __init__(self, titration, *args, **kwargs):
        super().__init__(*args, **kwargs)
        self.titration = titration
        self.title("Enter speciation table")

        height = int(self.master.winfo_height() * 0.4)
        self.frame = ttk.Frame(self, height=height)
        self.frame.pack(expand=True, fill="both")

        label = WrappedLabel(
            self.frame,
            padding=padding * 2,
            text=(
                "Each column corresponds to a component, and each row to a species.\n"
                "Define each complex by adding a row with the stoichiometry of each"
                " component in the complex. For polymers, use 'n'. Leaving a cell blank"
                " is identical to entering '0'."
            ),
        )
        label.pack(expand=False, fill="both")

        scrolledFrame = ScrolledFrame(
            self.frame, max_width=self.winfo_toplevel().master.winfo_width() - 200
        )
        scrolledFrame.pack(expand=True, fill="both")
        self.innerFrame = scrolledFrame.display_widget(ttk.Frame, stretch=True)

        self.speciationTable = SpeciationTable(self.innerFrame, titration)
        self.speciationTable.pack(expand=True, fill="both")

        buttonFrame = ButtonFrame(self.frame, self.reset, self.saveData, self.destroy)
        buttonFrame.pack(expand=False, fill="x", side="bottom")

    def reset(self):
        self.speciationTable.destroy()
        self.speciationTable = SpeciationTable(self.innerFrame, self.titration)
        self.speciationTable.pack(expand=True, fill="both")

    def saveData(self):
        self.stoichiometries = self.speciationTable.data

        self.saved = True
        self.destroy()


class SpeciationDimerisation(Speciation):
    @property
    def stoichiometries(self):
        M = np.array([[2]])
        M.resize([1, self.freeCount])
        return M

    noCooperativityValues = (np.array([1]), np.array([[1]]), np.array(["microK_Host₂"]))

    def run(self, variables, totalConcs):
        K = variables[0]
        Htot = totalConcs.T[0]
        H = (-1 + np.sqrt(1 + 8 * Htot * K)) / (4 * K)
        H2 = (1 + 4 * Htot * K - np.sqrt(1 + 8 * Htot * K)) / (8 * K)
        return np.array([H, H2]).T


class SpeciationHG(Speciation):
    @property
    def stoichiometries(self):
        M = np.array([[1, 1]])
        M.resize([1, self.freeCount])
        return M

    noCooperativityValues = (
        np.array([1]),
        np.array([[1]]),
        np.array(["microK_Host·Guest"]),
    )

    def run(self, variables, totalConcs):
        K = variables[0]
        Htot, Gtot = totalConcs.T
        H = (
            np.sqrt(
                Gtot**2 * K**2 - 2 * Gtot * K * (Htot * K - 1) + (Htot * K + 1) ** 2
            )
            - Gtot * K
            + Htot * K
            - 1
        ) / (2 * K)
        G = (
            np.sqrt(
                Htot**2 * K**2 - 2 * Htot * K * (Gtot * K - 1) + (Gtot * K + 1) ** 2
            )
            - Htot * K
            + Gtot * K
            - 1
        ) / (2 * K)
        HG = H * G * K

        return np.array([H, G, HG]).T


class SpeciationHG2(Speciation):
    @property
    def stoichiometries(self):
        if self.freeCount < 2:
            return np.array([[1] * self.freeCount])
        else:
            return np.pad(
                array=[[1, 1], [1, 2]],
                pad_width=[[0, 0], [0, self.freeCount - 2]],
                mode="constant",
                constant_values=0,
            )

    noCooperativityValues = (
        np.array([2, 1]),
        np.array([[1, 2]]),
        np.array(["microK_Host·Guest"]),
    )

    def run(self, variables, totalConcs):
        K1, K2 = variables
        output = np.empty([totalConcs.shape[0], 4])

        for i, (Htot, Gtot) in enumerate(totalConcs):
            if Htot == 0 or Gtot == 0:
                output[i] = [Htot, Gtot, 0, 0]
                continue

            # When K2 is very small, solving a cubic in [G] can be numerically unstable,
            # finding an inaccurate root, or not finding any real positive roots at
            # all. After some testing, it seems that solving a cubic in [G]/Gtot is more
            # stable, but I have not fully investigated the exact conditions under which
            # the eigenvalues algorithm used by LAPACK (used by np.roots) becomes
            # unstable, so adding error handling just in case.

            # Solve for a([G]/Gtot)^3 + b([G]/Gtot)^2 + c([G]/Gtot) + d == 0
            a = K2 * Gtot**3
            b = (K2 * (2 * Htot - Gtot) + K1) * Gtot**2
            c = (K1 * (Htot - Gtot) + 1) * Gtot
            d = -Gtot

            polynomial = np.array([a, b, c, d])

            roots = np.roots(polynomial)

            # Find smallest positive real root:
            select = np.all([np.imag(roots) == 0, np.real(roots) >= 0], axis=0)
            if np.count_nonzero(select) == 0:
                raise RuntimeError(
                    "No positive real roots found for cubic in [G]/Gtot when solving "
                    "speciation.\n\nThe most common cause is when some Ks and/or total "
                    "concentrations become very small or very large, leading to "
                    "precision errors. Please check that the initial guesses for all "
                    "variables are of a realistic order of magnitude, and that the "
                    "model isn't overdetermined. If the problem persists, try "
                    "selecting the 'Custom' binding isotherm option, which uses a "
                    f"slower but more robust algorithm.\n\nDetails: {K1=}, {K2=}, "
                    f"{Htot=}, {Gtot=}"
                )
            G = float(np.real(roots[select].min())) * Gtot

            H = Htot / (1 + K1 * G + K2 * (G**2))
            HG = K1 * H * G
            HG2 = K2 * H * G**2

            output[i] = [H, G, HG, HG2]

        return output


class SpeciationPolymerisation(Speciation):
    @property
    def stoichiometries(self):
        M = np.array([[-1]])
        M.resize([1, self.freeCount])
        return M

    noCooperativityValues = (
        np.array([1]),
        np.array([[1, 1]]),
        np.array(["microK_Hostₙ"]),
    )

    def run(self, variables, totalConcs):
        k2, kn = variables
        return np.array(
            [
                [
                    freeConc := self.polymerFreeExactSolutionSingle(k2, kn, totalConc),
                    *self.getTerminalInternalConcs(freeConc, k2, kn, None),
                ]
                for totalConc in totalConcs[:, 0]
            ]
        )


# Old, slower speciation algorithm, currently unused. Left in in case results need to
# be compared against it.
class SpeciationCOGS(Speciation):
    def COGS(self, M, y, ks, polymerKs):
        free = y.copy()
        bound = np.empty(len(ks))
        polymers = np.any(M < 0, 1)
        # the index of the species making up each of the polymers
        polymerParents = np.nonzero(M[polymers])[1]
        complexes = ~polymers

        P = np.empty(len(ks))
        P[complexes] = np.sum(M[complexes, :], 1)
        P[polymers] = 2 * ks[polymers] * y[polymerParents]
        P = 1 / max(P)
        tol = 1e-7
        while True:
            bound[complexes] = ks[complexes] * np.prod(free ** M[complexes, :], 1)
            # cap the maximum guess to avoid divergence
            bound[polymers] = np.where(
                free[polymerParents] * polymerKs[polymers] >= 1,
                free[polymerParents] ** 2 * polymerKs[polymers] / P,
                (2 - polymerKs[polymers] * free[polymerParents])
                * (ks[polymers] * free[polymerParents] ** 2)
                / ((1 - polymerKs[polymers] * free[polymerParents]) ** 2),
            )
            total = free + abs(M.T) @ bound  # total concentrations of species
            if all((total - y) <= tol * y):
                break
            # to handle 0 total concentration
            invTotal = np.where(total == 0, 1, 1 / total)
            free *= (y * invTotal) ** P

        # For polymers, return separate entries for terminal and internal groups
        bound = []
        paddedPolymerParents = ma.array(np.empty(len(ks)), mask=True)
        paddedPolymerParents[polymers] = polymerParents
        for k, polymerK, stoichiometries, isPolymer, parent in zip(
            ks, polymerKs, M, self.polymerIndices, paddedPolymerParents
        ):
            if not isPolymer:
                bound.append(k * np.prod(free**stoichiometries))
            else:
                # terminal
                bound.append(2 * free[parent] ** 2 * k / (1 - free[parent] * polymerK))
                # internal
                bound.append(
                    (free[parent] ** 3 * k * polymerK)
                    / (1 - free[parent] * polymerK) ** 2
                )

        return free, np.array(bound)

    def run(self, variables, totalConcs):
        ks, polymerKs = self.variablesToKs(variables)
        numPoints = totalConcs.shape[0]
        free = np.zeros((numPoints, self.titration.totalConcentrations.freeCount))
        bound = np.zeros((numPoints, self.complexCount + 2 * self.polymerCount))
        for i in range(numPoints):
            free[i], bound[i] = self.COGS(
                self.stoichiometries, totalConcs[i], ks, polymerKs
            )
        return free, bound


class SpeciationSolver(Speciation):
    # Optimise f(total * log10(free))
    # This makes the derivative equal to ln(10) * (free + bound - total) / total
    #
    # So gtol in the minimisation algorithm can be set to the desired precision in
    # the total concentrations.
    def objective(
        self, logFreeTimesTotal, complexKs, k2s, kns, kabs, total, complexM, polymerM
    ):
        free = 10 ** (logFreeTimesTotal / total)
        return (
            np.sum(free)
            - np.sum(logFreeTimesTotal) * LN_10
            + self.complexObjective(free, complexKs, total, complexM)
            + self.polymerObjective(free, k2s, kns, kabs, total, polymerM)
        )

    def jacobian(
        self, logFreeTimesTotal, complexKs, k2s, kns, kabs, total, complexM, polymerM
    ):
        free = 10 ** (logFreeTimesTotal / total)
        return (
            (
                free
                + self.complexJacobian(free, complexKs, total, complexM)
                + self.polymerJacobian(free, k2s, kns, kabs, total, polymerM)
                - total
            )
            * LN_10
            / total
        )

    def objectiveScaled(
        self, logFreeTimesTotal, complexKs, k2s, kns, kabs, total, complexM, polymerM
    ):
        free = 10 ** (logFreeTimesTotal / self.scaling_factor / total)
        return (
            np.sum(free)
            - np.sum(logFreeTimesTotal / self.scaling_factor) * LN_10
            + self.complexObjective(free, complexKs, total, complexM)
            + self.polymerObjective(free, k2s, kns, kabs, total, polymerM)
        ) * self.scaling_factor

    def jacobianScaled(
        self, logFreeTimesTotal, complexKs, k2s, kns, kabs, total, complexM, polymerM
    ):
        free = 10 ** (logFreeTimesTotal / self.scaling_factor / total)
        return (
            (
                free
                + self.complexJacobian(free, complexKs, total, complexM)
                + self.polymerJacobian(free, k2s, kns, kabs, total, polymerM)
                - total
            )
            * LN_10
            / total
        )

    def smoothObjective(self, logFreeTimesTotal, *args, **kwargs):
        upperBounds = self.getDomainUpperBounds(*args, **kwargs)
        if np.all(logFreeTimesTotal <= upperBounds):
            return self.objective(logFreeTimesTotal, *args, **kwargs)
        print("used truncation in objective")

        truncatedX = np.clip(logFreeTimesTotal, None, upperBounds)
        return self.objective(truncatedX, *args, **kwargs) + np.sum(
            (logFreeTimesTotal - truncatedX)
            * self.jacobian(truncatedX, *args, **kwargs)
        )

    def smoothJacobian(self, logFreeTimesTotal, *args, **kwargs):
        upperBounds = self.getDomainUpperBounds(*args, **kwargs)
        if np.all(logFreeTimesTotal <= upperBounds):
            return self.jacobian(logFreeTimesTotal, *args, **kwargs)
        print("used truncation in jacobian")

        truncatedX = np.clip(logFreeTimesTotal, None, upperBounds)
        return self.jacobian(truncatedX, *args, **kwargs)

    def getDomainUpperBounds(
        self, complexKs, k2s, kns, kabs, total, complexM, polymerM
    ):
        return total * np.log10(
            self.polymerGetUpperBounds(k2s, kns, kabs, total, polymerM),
        )

    # Could be refined iteratively, by computing the LB using this method, then taking
    # UB = self.freeToBoundConcs(free=LB), calculating a new LB using that UB, etc.
    def getUpperBounds(self, complexKs, k2s, kns, kabs, total, complexM, polymerM):
        return total * np.log10(
            np.minimum(
                self.complexGetUpperBounds(complexKs, total, complexM),
                self.polymerGetUpperBounds(k2s, kns, kabs, total, polymerM),
            )
        )

    def getLowerBounds(self, complexKs, k2s, kns, kabs, total, complexM, polymerM):
        maxFree = np.minimum(
            self.complexGetUpperBounds(complexKs, total, complexM),
            self.polymerGetUpperBounds(k2s, kns, kabs, total, polymerM),
        )
        return total * np.log10(
            (total * maxFree)
            / (
                maxFree
                + self.complexJacobian(maxFree, complexKs, total, complexM)
                + self.polymerJacobian(maxFree, k2s, kns, kabs, total, polymerM)
            )
        )

    def run(self, variables, totalConcs):
        complexKs, k2s, kns, kabs = self.variablesToKs(variables)
        numPoints = totalConcs.shape[0]

        free = np.empty((numPoints, self.freeCount))
        bound = np.empty((numPoints, self.outputCount - self.freeCount))

        for i in range(numPoints):
            additionTotalConcs = totalConcs[i]

            # Filter the to exclude species and complexes that will have a concentration
            # of 0
            zeroFree = additionTotalConcs == 0
            zeroBound = np.any(self.stoichiometries[:, zeroFree], axis=1)
            if all(zeroBound):
                free[i] = additionTotalConcs
                bound[i] = 0
                continue
            zeroComplexes = zeroBound[~self.polymerIndices]
            zeroPolymers = zeroBound[self.polymerIndices]

            filteredKs = complexKs[~zeroComplexes]
            filteredK2s = k2s[~zeroFree]
            filteredKns = kns[~zeroFree]
            filteredKabs = kabs[~zeroPolymers]

            filteredTotal = additionTotalConcs[~zeroFree]
            filteredComplexM = self.complexStoichiometries[~zeroComplexes, :][
                :, ~zeroFree
            ]
            filteredPolymerM = self.polymerStoichiometries[~zeroPolymers, :][
                :, ~zeroFree
            ]

            args = (
                filteredKs,
                filteredK2s,
                filteredKns,
                filteredKabs,
                filteredTotal,
                filteredComplexM,
                filteredPolymerM,
            )

            lb = self.getLowerBounds(*args)
            ub = self.getUpperBounds(*args)
            if any(lb > ub):
                # Correct for rounding errors. Unsure if this is necessary, as the only
                # obvious case when it should happen is if no complexes are formed,
                # which should be caught above.
                mask = np.logical_and(lb > ub, np.isclose(lb, ub))
                lb[mask], ub[mask] = ub[mask], lb[mask]

            # TODO: deal with cases where lb and ub are very close together!
            # self.scaling_factor = 1000 / min(ub - lb)
            self.scaling_factor = 1000 / np.min(
                np.abs(filteredTotal * np.log10(filteredTotal))
            )
            self.scaling_factor = 1

            if i == 0:
                # Initial guess: all species 100% free, only polymers are formed
                x0 = ub
            else:
                # If the total concentration increased, the initial guess is that all
                # the added molecules are free.
                # If the total concentration decreased, the initial guess is that the
                # free concentration decreases by the same fraction.
                initialGuess = free[i - 1].copy()
                difference = additionTotalConcs - totalConcs[i - 1]
                concsIncreased = difference >= 0

                initialGuess[concsIncreased] += difference[concsIncreased]
                initialGuess[~concsIncreased] *= (
                    totalConcs[i][~concsIncreased] / totalConcs[i - 1][~concsIncreased]
                )
                x0 = filteredTotal * np.log10(initialGuess[~zeroFree])
                x0 = np.clip(x0, lb, ub)

            result = minimize(
                self.objectiveScaled,
                jac=self.jacobianScaled,
                args=args,
                x0=x0 * self.scaling_factor,
                bounds=np.vstack([lb, ub]).T * self.scaling_factor,
                method="L-BFGS-B",
                options={
                    "ftol": 0.0,
                    "gtol": 1e-6 * LN_10,
                },
            )
            if result.success and "jac" not in result.keys():
                # Happens if all lower bounds are equal to upper bounds, and possibly
                # also in other cases.
                result.jac = self.jacobianScaled(result.x, *args)
            if max(abs(result.jac)) > 1e-6 * LN_10:
                self.scaling_factor *= 10_000
                improvedResult = minimize(
                    self.objectiveScaled,
                    jac=self.jacobianScaled,
                    args=args,
                    x0=result.x,
                    bounds=np.vstack([lb, ub]).T * self.scaling_factor,
                    method="L-BFGS-B",
                    options={
                        "ftol": 0.0,
                        "gtol": 1e-6 * LN_10,
                    },
                )
                if improvedResult.success and "jac" not in improvedResult.keys():
                    improvedResult.jac = self.jacobianScaled(improvedResult.x, *args)

                if max(abs(improvedResult.jac)) < max(abs(result.jac)):
                    result = improvedResult
                else:
                    warnings.warn(
                        "Desired accuracy not achieved in speciation",
                        RuntimeWarning,
                    )

            logFree = result.x / self.scaling_factor / filteredTotal

            free[i, ~zeroFree] = 10**logFree
            free[i, zeroFree] = 0
            # get the concentrations of the bound species from those of the free
            bound[i] = self.freeToBoundConcs(free[i], complexKs, k2s, kns, kabs)

        return np.hstack([free, bound])


class SpeciationCustom(SpeciationSolver):
    Popup = SpeciationPopup
    popupAttributes = ("stoichiometries",)

class SpeciationPolymer1D_1s(Speciation):
    """
    Custom speciation model implementing equations for binding to a 1D linear polymer.
    Code is incorporated for modelling ligand binding to a single type of site.
    """

    # Stoichiometry table
    @property
    def stoichiometries(self):
        # free species + 1 bound species
        st = np.zeros((1, self.freeCount), dtype=int)

        # mark that this output involves ligand A (assume A is the 2nd component for now)
        if self.freeCount >= 2:
            st[0, 1] = 1
        return st

    # Model variables
    variableNames = np.array(["K_A","omega_AA","n_A"])

    @property
    def outputNames(self):
        # First columns are free species, then Bound Guest and v_A
        return np.concatenate([self.freeNames, ["Bound Guest"]])

    def calculate_1_s_x(self, n_A, v_A, p_A, q_A, omega_AA, ff):
        """
        Average number of binding sites per lattice unit for a single-ligand model.   

        Returns
        -------
        s_A : float
            Av. binding site density  used in the mass-action equations for A.
        """
        eps = 1e-14
        denom_cov = max(n_A * v_A, eps)
        ff_ratio = ff / max(1 - ff, eps)

        sum_left_A  = q_A * (omega_AA + ff_ratio)
        sum_right_A = p_A * (omega_AA + ff_ratio)

        s_A = sum_left_A * ff ** (n_A - 1) * sum_right_A
        return s_A

    def equations_1lig_multisite(self, vars_flat, params):
        """
        Residuals for the 1-ligand, multi-site-class equilibrium model.

        Unknowns per site class i (block size 4):
        [v_A_i, p_A_i, q_A_i, ff_i]
        plus global free pool:
        [L_A_free]

        params keys:
        - site_params (list of dicts): res_tot, K_A, omega_AA
        - n_A
        - L_A_tot
        """

        site_params = params["site_params"]
        T = len(site_params) # number of site classes
        n_A = params["n_A"]     
        L_A_tot = params["L_A_tot"]

        v_A = vars_flat[0*T:1*T]
        p_A = vars_flat[1*T:2*T]
        q_A = vars_flat[2*T:3*T]
        ff  = vars_flat[3*T:4*T]
        L_A_free = vars_flat[4*T]

        res, total_bound_A = [], 0.0
        eps = 1e-14

        for i, spi in enumerate(site_params):
            res_tot_i = spi["res_tot"]
            K_A_i = spi["K_A"]
            ω_AA = spi.get("omega_AA", 1.0)

            vAi, pAi, qAi, ffi = v_A[i], p_A[i], q_A[i], ff[i]
            theta_i = n_A * vAi
            bound_A_i = vAi * res_tot_i
            res_free_i = res_tot_i * (1 - theta_i)

            s_A_i = self.calculate_1_s_x(n_A, vAi, pAi, qAi, ω_AA, ffi)

            # E1–E2: normalization 
            eq1 = ffi + pAi - 1
            eq2 = ffi * (1 - theta_i) + vAi * qAi - (1 - theta_i)

            # E3: cooperativity influence on conditional probabilities 
            eq3 = qAi + ω_AA * qAi * pAi / max(ffi, eps) - 1

            # E4: mass action 
            eq4 = 1 - K_A_i * s_A_i * L_A_free

            # E5: linking conditional probabilities and concentrations
            eq5 = pAi * res_free_i / res_tot_i + (ω_AA * qAi * pAi / max(ffi, eps) - 1) * bound_A_i / res_tot_i

            res.extend([eq1, eq2, eq3, eq4, eq5])
            total_bound_A += bound_A_i

        # Global ligand conservation 
        res.append((L_A_free + total_bound_A - L_A_tot) / res_tot_i)

        return res # returns residuals

    def run(self, variables, totalConcs):

        additionCount = totalConcs.shape[0]
        n_concs = totalConcs.shape[1]
        output = np.zeros([additionCount, self.outputCount], dtype=float)

        K_A, omega_AA, n_A = variables

        # negative cooperativity is unlikely, and omega higher than 10 is unlikely
        omega_AA = float(np.clip(omega_AA, 0.9, 10.0)) 

        # also constraining possible n_A values between 0.5 and 15, rounded to nearest 0.1
        n_A = float(np.clip(n_A, 0.5, 15.0))

        res_tot_values = totalConcs[:, 0] # first column is total host concentration
        L_A_tot_values = totalConcs[:,1]  # second column is total guest concentration

        # site_params for one-site model still includes redundant B variables for compatibility
        # This is an inelegant fix as I redefine site_params in the main code
        site_params = [
            dict(res_tot=0, K_A=K_A, K_B=0, omega_AA=omega_AA, omega_AB=1, omega_BA=1, omega_BB=1)
        ]

        T = len(site_params)
        eps = 1e-14

        # setting up bounds for unknowns
        lb = [0]*T + [eps]*T + [eps]*T + [eps]*T + [0]
        ub_v = min(1.0, 1.0 / n_A)
        ub = [ub_v]*T + [1]*T + [1]*T + [1]*T + [max(L_A_tot_values)]
        
        prev_sol = None

        free_lattice = []
        free_guest = []
        bound_guest = []
        v_A_list = []

        for res_tot, L_A_tot in zip(res_tot_values, L_A_tot_values):

            # --- Early exit for zero (or effectively zero) ligand ---
            tiny = 1e-12
            if L_A_tot <= tiny:
                # Physical limit
                free_lattice.append(res_tot)   # all lattice free
                free_guest.append(0.0)
                bound_guest.append(0.0)
                v_A_list.append(0.0)

                prev_sol = None  # reset initial guess safely
                continue         # IMPORTANT: continue loop, do NOT return
            
            site_params = [
                dict(res_tot=res_tot, K_A=K_A, K_B=0, omega_AA=omega_AA, omega_AB=1, omega_BA=1, omega_BB=1)
            ]

            # x0 is the initial guess for unknowns in least_squares, format v_A, p_A, q_A, ff, L_A_free
            if prev_sol is None:
                x0 = ([1/max(n_A+1, 2)] * T + [0.1] * T + [0.9] * T + [0.5] * T + [0.5 * L_A_tot])
            else:
                x0 = prev_sol

            res = least_squares(
                self.equations_1lig_multisite, x0,
                args=({"site_params": site_params, "n_A": n_A, "L_A_tot": L_A_tot},),
                bounds=(lb, ub), method="trf",
                xtol=1e-10, ftol=1e-10, diff_step=1e-6, jac="3-point"
            )

            sol = res.x

            prev_sol = sol

            v_A = sol[0*T:1*T]
            p_A = sol[1*T:2*T]
            q_A = sol[2*T:3*T]
            ff  = sol[3*T:4*T]
            L_A_free = sol[4*T]

            theta = float(n_A * v_A[0])
            theta = min(max(theta, 0.0), 1.0) # making sure theta is within [0,1]
            free_lattice.append(res_tot * (1 - theta))
            free_guest.append(L_A_free)
            bound_guest.append(L_A_tot - L_A_free)
            v_A_list.append(n_A * v_A[0])  # only one site class for now
        
        #["Total Guest", "Total Lattice", "Bound Guest", "v_A"]])
        # First column: free host
        output[:,0] = free_lattice
        # Second column: free guest
        output[:,1] = free_guest
        # Third column: bound guest
        output[:,2] = bound_guest
        return output

class SpeciationPolymer1D_2s(Speciation):
    """
    Custom speciation model implementing equations for binding to a 1D linear polymer.
    Code is incorporated for modelling ligand binding to two types of sites.
    """

    # Stoichiometry table
    @property
    def stoichiometries(self):
        # free species + 2 bound species
        st = np.zeros((2, self.freeCount), dtype=int)

        # stoichiometry format: Host 1, Host 2, Guest
        if self.freeCount >= 2:
            # Site 1 + guest
            st[0, 0] = 1
            st[0, 2] = 1
            # Site 2 + guest
            st[1, 1] = 1
            st[1, 2] = 1
        return st

    # Model variables
    variableNames = np.array(["K_A1","omega_AA1","n_A1","K_A2","omega_AA2","n_A2"])

    @property
    def outputNames(self):
        # First columns are free species, then Bound Guest for each site
        return np.concatenate([self.freeNames, ["Bound Guest s1","Bound Guest s2"]])

    def calculate_1_s_x(self, n_A, v_A, p_A, q_A, omega_AA, ff):
        """
        Average number of binding sites per lattice unit for a single-ligand model.   

        Returns
        -------
        s_A : float
            Av. binding site density  used in the mass-action equations for A.
        """
        eps = 1e-14
        denom_cov = max(n_A * v_A, eps)
        ff_ratio = ff / max(1 - ff, eps)

        sum_left_A  = q_A * (omega_AA + ff_ratio)
        sum_right_A = p_A * (omega_AA + ff_ratio)

        s_A = sum_left_A * ff ** (n_A - 1) * sum_right_A
        return s_A

    def equations_1lig_multisite(self, vars_flat, params):
        """
        Residuals for the 1-ligand (A only), multi-site-class equilibrium model.

        Unknowns per site class i (block size 4):
        [v_A_i, p_A_i, q_A_i, ff_i]
        plus global free pool:
        [L_A_free]

        params keys:
        - site_params (list of dicts): res_tot, K_A, omega_AA, n_A
        - L_A_tot
        """

        site_params = params["site_params"]
        T = len(site_params) # number of site classes
        L_A_tot = params["L_A_tot"]

        v_A = vars_flat[0*T:1*T]
        p_A = vars_flat[1*T:2*T]
        q_A = vars_flat[2*T:3*T]
        ff  = vars_flat[3*T:4*T]
        L_A_free = vars_flat[4*T]

        res, total_bound_A = [], 0.0
        eps = 1e-14

        for i, spi in enumerate(site_params):
            res_tot_i = spi["res_tot"]
            K_A_i = spi["K_A"]
            n_A_i = spi["n_A"]
            ω_AA = spi.get("omega_AA", 1.0)

            vAi, pAi, qAi, ffi = v_A[i], p_A[i], q_A[i], ff[i]
            theta_i = n_A_i * vAi
            bound_A_i = vAi * res_tot_i
            res_free_i = res_tot_i * (1 - theta_i)

            s_A_i = self.calculate_1_s_x(n_A_i, vAi, pAi, qAi, ω_AA, ffi)
            # E1–E2: normalization 
            eq1 = ffi + pAi - 1
            eq2 = ffi * (1 - theta_i) + vAi * qAi - (1 - theta_i)

            # E3: cooperativity influence on conditional probabilities 
            eq3 = qAi + ω_AA * qAi * pAi / max(ffi, eps) - 1

            # E4: mass action 
            eq4 = 1 - K_A_i * s_A_i * L_A_free

            # E5: linking conditional probabilities and concentrations
            eq5 = pAi * res_free_i / res_tot_i + (ω_AA * qAi * pAi / max(ffi, eps) - 1) * bound_A_i / res_tot_i

            res.extend([eq1, eq2, eq3, eq4, eq5])
            total_bound_A += bound_A_i

        # Global ligand conservation (A only)
        scale = max(L_A_tot, 1e-14)
        res.append((L_A_free + total_bound_A - L_A_tot) / scale)

        return res # returns residuals

    def run(self, variables, totalConcs):

        additionCount = totalConcs.shape[0]
        n_concs = totalConcs.shape[1]
        output = np.zeros([additionCount, self.outputCount], dtype=float)

        K_A1, omega_AA1, n_A1, K_A2, omega_AA2, n_A2 = variables

        # negative cooperativity is unlikely, and omega higher than 10 is unlikely
        omega_AA1 = float(np.clip(omega_AA1, 0.9, 10.0)) 
        omega_AA2 = float(np.clip(omega_AA2, 0.9, 10.0)) 

        # also constraining possible n_A values between 1 and 10, rounded to nearest 0.1
        n_A1 = float(np.clip(n_A1, 1, 10.0))
        n_A2 = float(np.clip(n_A2, 1, 10.0))
 
        res1_tot_values = totalConcs[:, 0] # first column is total site 1 concentration
        res2_tot_values = totalConcs[:, 1] # second column is total site 2 concentration
        L_A_tot_values = totalConcs[:,2]  # third column is total guest concentration

        # site_params for two-site model still includes redundant B variables for compatibility
        # This is an inelegant fix as I redefine site_params in the main code
        site_params = [
            dict(res_tot=0, K_A=K_A1, K_B=0, omega_AA=omega_AA1, omega_AB=1, omega_BA=1, omega_BB=1, n_A=n_A1),
            dict(res_tot=0, K_A=K_A2, K_B=0, omega_AA=omega_AA2, omega_AB=1, omega_BA=1, omega_BB=1, n_A=n_A2)
        ]

        T = len(site_params)
        eps = 1e-14

        # setting up bounds for unknowns
        lb = [0]*T + [eps]*T + [eps]*T + [eps]*T + [0]
        ub_v = [min(1.0, 1.0/n_A1), min(1.0, 1.0/n_A2)]
        ub = ub_v + [1]*T + [1]*T + [1]*T + [max(L_A_tot_values)]
        
        prev_sol = None

        free_lattice_1 = []
        free_lattice_2 = []
        free_guest = []
        bound_guest_1 = []
        bound_guest_2 = []
        v_A_list_1 = []
        v_A_list_2 = []

        for res_tot_1, res_tot_2, L_A_tot in zip(res1_tot_values, res2_tot_values, L_A_tot_values):

            # --- Early exit for zero (or effectively zero) ligand ---
            tiny = 1e-12
            if L_A_tot <= tiny:
                # Physical limit
                free_lattice_1.append(res_tot_1)   # all lattice free
                free_lattice_2.append(res_tot_2)   # all lattice free
                free_guest.append(0.0)
                bound_guest_1.append(0.0)
                bound_guest_2.append(0.0)
                v_A_list_1.append(0.0)
                v_A_list_2.append(0.0)

                prev_sol = None  # reset initial guess safely
                continue         # IMPORTANT: continue loop, do NOT return
            
            site_params = [
                dict(res_tot=res_tot_1, K_A=K_A1, K_B=0, omega_AA=omega_AA1, omega_AB=1, omega_BA=1, omega_BB=1, n_A=n_A1),
                dict(res_tot=res_tot_2, K_A=K_A2, K_B=0, omega_AA=omega_AA2, omega_AB=1, omega_BA=1, omega_BB=1, n_A=n_A2)
            ]

            n_As = []
            for site in site_params:
                n_As.append(site["n_A"])
            n_A1, n_A2 = n_As

            # x0 is the initial guess for unknowns in least_squares, format v_A, p_A, q_A, ff, L_A_free
            if prev_sol is None:
                x0 = ([1/(n+1) for n in n_As] + [0.1]*T + [0.9]*T + [0.5]*T + [0.5*L_A_tot])
            else:
                x0 = prev_sol

            # Note - ocassionally issue where n escapes bounds before least_squares from prev_sol
            x0 = np.asarray(x0, dtype=float)
            lb_arr = np.asarray(lb, dtype=float)
            ub_arr = np.asarray(ub, dtype=float)

            # Project into bounds (with a tiny margin to avoid edge singularities)
            x0 = np.maximum(x0, lb_arr + 1e-12)
            x0 = np.minimum(x0, ub_arr - 1e-12)

            r0 = np.array(self.equations_1lig_multisite(
                x0, {"site_params": site_params, "L_A_tot": L_A_tot}
            ), dtype=float)

            if not np.all(np.isfinite(r0)):
                bad = np.where(~np.isfinite(r0))[0]
                raise RuntimeError(f"Non-finite residuals at x0: idx={bad}, vals={r0[bad]}")

            res = least_squares(
                self.equations_1lig_multisite, x0,
                args=({"site_params": site_params, "L_A_tot": L_A_tot},),
                bounds=(lb, ub), method="trf",
                xtol=1e-10, ftol=1e-10, diff_step=1e-6, jac="3-point"
            )

            sol = res.x

            prev_sol = sol

            v_A = sol[0*T:1*T]
            p_A = sol[1*T:2*T]
            q_A = sol[2*T:3*T]
            ff  = sol[3*T:4*T]
            L_A_free = sol[4*T]

            theta1 = float(n_A1 * v_A[0])
            theta2 = float(n_A2 * v_A[1])
            theta1 = min(max(theta1, 0.0), 1.0) # making sure theta is within [0,1]
            theta2 = min(max(theta2, 0.0), 1.0) # making sure theta is within [0,1]     
            free_lattice_1.append(res_tot_1 * (1 - theta1))
            free_lattice_2.append(res_tot_2 * (1 - theta2))
            free_guest.append(L_A_free)

            bound_guest_1.append(v_A[0] * res_tot_1)
            bound_guest_2.append(v_A[1] * res_tot_2)

            v_A_list_1.append(n_A1 * v_A[0]) 
            v_A_list_2.append(n_A2 * v_A[1])

        #["Total Guest", "Total Lattice", "Bound Guest", "v_A"]])
        # First column: free lattice 1
        output[:,0] = free_lattice_1
        # First column: free lattice 2
        output[:,1] = free_lattice_2
        # Third column: free guest
        output[:,2] = free_guest
        # Fourth column: bound guest-lattice 1
        output[:,3] = bound_guest_1
        # Fifth column: bound guest-lattice 2
        output[:,4] = bound_guest_2

        return output

class SpeciationPolymer1D_1s_adjacency(Speciation):
    """
    Custom speciation model implementing equations for binding to a 1D linear polymer.
    Code is incorporated for modelling ligand binding to a single type of site, 
    with separate outputs for adjacent vs non-adjacent binding to enable excimer modelling.
    """

    # Stoichiometry table
    @property
    def stoichiometries(self):
        # free species + 2 bound species
        st = np.zeros((2, self.freeCount), dtype=int)

        # mark that this output involves ligand A (assume A is the 2nd component for now)
        if self.freeCount >= 2:
            # non-adjacent bound state
            st[0, 0] = 1
            st[0, 1] = 1
            # adjacent bound state
            st[1, 0] = 1
            st[1, 1] = 1
        return st

    # Model variables
    variableNames = np.array(["K_A","omega_AA","n_A"])

    @property
    def outputNames(self):
        # First columns are free species, then Bound Guest and v_A
        return np.concatenate([self.freeNames, ["Bound Guest Non-Adjacent", "Bound Guest Adjacent"]])

    def calculate_1_s_x(self, n_A, v_A, p_A, q_A, omega_AA, ff):
        """
        Average number of binding sites per lattice unit for a single-ligand model.   

        Returns
        -------
        s_A : float
            Av. binding site density  used in the mass-action equations for A.
        """
        eps = 1e-14
        ff_ratio = ff / max(1 - ff, eps)

        sum_left_A  = q_A * (omega_AA + ff_ratio)
        sum_right_A = p_A * (omega_AA + ff_ratio)

        s_A = sum_left_A * ff ** (n_A - 1) * sum_right_A
        return s_A

    def equations_1lig_multisite(self, vars_flat, params):
        """
        Residuals for the 1-ligand (A only), multi-site-class equilibrium model.

        Unknowns per site class i (block size 4):
        [v_A_i, p_A_i, q_A_i, ff_i]
        plus global free pool:
        [L_A_free]

        params keys:
        - site_params (list of dicts): res_tot, K_A, omega_AA
        - n_A
        - L_A_tot
        """

        site_params = params["site_params"]
        T = len(site_params) # number of site classes
        n_A = params["n_A"]     
        L_A_tot = params["L_A_tot"]

        v_A = vars_flat[0]
        p_A = vars_flat[1]
        q_A = vars_flat[2]
        ff  = vars_flat[3]
        L_A_free = vars_flat[4]

        res, total_bound_A = [], 0.0
        eps = 1e-14

        for i, spi in enumerate(site_params):
            res_tot_i = spi["res_tot"]
            K_A_i = spi["K_A"]
            ω_AA = spi.get("omega_AA", 1.0)

            vAi, pAi, qAi, ffi = v_A, p_A, q_A, ff
            theta_i = n_A * vAi
            bound_A_i = vAi * res_tot_i
            res_free_i = res_tot_i * (1 - theta_i)

            s_A_i = self.calculate_1_s_x(n_A, vAi, pAi, qAi, ω_AA, ffi)

            # E1–E2: normalization 
            eq1 = ffi + pAi - 1
            eq2 = ffi * (1 - theta_i) + vAi * qAi - (1 - theta_i)

            # E3: cooperativity influence on conditional probabilities 
            eq3 = qAi + ω_AA * qAi * pAi / max(ffi, eps) - 1

            # E4: mass action 
            eq4 = 1 - K_A_i * s_A_i * L_A_free

            # E5: linking conditional probabilities and concentrations
            eq5 = pAi * res_free_i / res_tot_i + (ω_AA * qAi * pAi / max(ffi, eps) - 1) * bound_A_i / res_tot_i

            res.extend([eq1, eq2, eq3, eq4, eq5])
            total_bound_A += bound_A_i

        # Global ligand conservation
        res.append((L_A_free + total_bound_A - L_A_tot) / res_tot_i)

        return res # returns residuals

    def run(self, variables, totalConcs):

        additionCount = totalConcs.shape[0]
        output = np.zeros([additionCount, self.outputCount], dtype=float)

        K_A, omega_AA, n_A = variables

        # negative cooperativity is unlikely, and omega higher than 10 is unlikely
        omega_AA = float(np.clip(omega_AA, 0.9, 10.0)) 

        # also constraining possible n_A values between 0.5 and 10, rounded to nearest 0.1
        n_A = float(np.clip(n_A, 0.5, 10.0))

        res_tot_values = totalConcs[:, 0] # first column is total host concentration
        L_A_tot_values = totalConcs[:,1]  # second column is total guest concentration

        # site_params for one-site model still includes redundant B variables for compatibility
        # This is an inelegant fix as I redefine site_params in the main code
        site_params = [
            dict(res_tot=0, K_A=K_A, K_B=0, omega_AA=omega_AA, omega_AB=1, omega_BA=1, omega_BB=1)
        ]

        T = len(site_params)
        eps = 1e-14

        # setting up bounds for unknowns
        lb = [0]*T + [eps]*T + [eps]*T + [eps]*T + [0]
        ub_v = min(1.0, 1.0 / n_A)
        ub = [ub_v]*T + [1]*T + [1]*T + [1]*T + [max(L_A_tot_values)]
        
        prev_sol = None

        free_lattice = []
        free_guest = []
        bound_guest_nonAdjacent = []
        bound_guest_adjacent = []
        v_A_list = []

        for res_tot, L_A_tot in zip(res_tot_values, L_A_tot_values):

            # --- Early exit for zero (or effectively zero) ligand ---
            tiny = 1e-12
            if L_A_tot <= tiny:
                # Physical limit
                free_lattice.append(res_tot)   # all lattice free
                free_guest.append(0.0)
                bound_guest_nonAdjacent.append(0.0)
                bound_guest_adjacent.append(0.0)
                v_A_list.append(0.0)

                prev_sol = None  # reset initial guess safely
                continue         # IMPORTANT: continue loop, do NOT return
            
            site_params = [
                dict(res_tot=res_tot, K_A=K_A, K_B=0, omega_AA=omega_AA, omega_AB=1, omega_BA=1, omega_BB=1)
            ]

            # x0 is the initial guess for unknowns in least_squares, format v_A, p_A, q_A, ff, L_A_free
            if prev_sol is None:
                x0 = ([1/max(n_A+1, 2)] * T + [0.1] * T + [0.9] * T + [0.5] * T + [0.5 * L_A_tot])
            else:
                x0 = prev_sol

            res = least_squares(
                self.equations_1lig_multisite, x0,
                args=({"site_params": site_params, "n_A": n_A, "L_A_tot": L_A_tot},),
                bounds=(lb, ub), method="trf",
                xtol=1e-10, ftol=1e-10, diff_step=1e-6, jac="3-point"
            )

            sol = res.x

            prev_sol = sol

            v_A = sol[0]
            p_A = sol[1]
            q_A = sol[2]
            ff  = sol[3]
            L_A_free = sol[4]

            bb = 1 - q_A

            theta = float(n_A * v_A)
            theta = min(max(theta, 0.0), 1.0) # making sure theta is within [0,1]
            free_lattice.append(res_tot * (1 - theta))
            free_guest.append(L_A_free)
            bound_guest = L_A_tot - L_A_free

            bound_guest_a = bound_guest * bb
            bound_guest_nonA = bound_guest * (1 - bb)

            bound_guest_adjacent.append(bound_guest_a)
            bound_guest_nonAdjacent.append(bound_guest_nonA)
        
        #["Total Guest", "Total Lattice", "Bound Guest Non-Adjacent", "Bound Guest Adjacent"]])
        # First column: free host
        output[:,0] = free_lattice
        # Second column: free guest
        output[:,1] = free_guest
        # Third column: bound guest non-adjacent
        output[:,2] =   bound_guest_nonAdjacent
        # Fourth column: bound guest adjacent
        output[:,3] = bound_guest_adjacent

        return output


class ModuleFrame(moduleFrame.ModuleFrame):
    group = "Equilibria"
    dropdownLabelText = "Select a binding isotherm:"
    dropdownOptions = {
        "1:1 binding": SpeciationHG,
        "1:2 binding": SpeciationHG2,
        "Dimerisation": SpeciationDimerisation,
        "Polymerisation": SpeciationPolymerisation,
        "1D Polymer Binding, 1-site": SpeciationPolymer1D_1s,
        "1D Polymer Binding, 2-site": SpeciationPolymer1D_2s,
        "1D Polymer Binding, 1-site (Adjacency)": SpeciationPolymer1D_1s_adjacency,
        "Custom": SpeciationCustom,
        # "Custom Grad": SpeciationCustomGrad,
    }
    attributeName = "speciation"
```
