## Supplementary Information for "Two Distinct Binding Modes Govern High-Affinity Ligand Interactions with Amyloid Fibrils"

Timothy S. Chisholm [a,*]

[a] Yusuf Hamied Department of Chemistry, University of Cambridge, Lensfield Road, Cambridge CB2 1EW, UK..

### Supporting Information

### Contents

Materials and Instrumentation.......... S3
Small Molecule Synthesis .......... S4
UV-Visible Characterisation.......... S15
Fluorescence Characterisation.......... S16
ThT Hill Plots .......... S18
Preparation of αSyn Fibrils.......... S18
Binding Model Derivation .......... S19
Cooperativity parameter .......... S19
Gap probability .......... S19
Computing the number of free binding sites .......... S19
Final system of equations .......... S21
Synthetic Binding Data Generation .......... S23
Additional Synthetic Binding Data .......... S24
Radioligand Self-Displacement Assay Data .......... S25
Code for Isotherm Generation and Fitting. .......... S26
Comparing Fits .......... S26
Symbols Used in Equations .......... S27

#### Materials and Instrumentation

All solvents and chemicals were obtained from commercial sources and used without further purification unless otherwise stated. Protein LoBind (Eppendorf) microtubes were used for preparing and storing all solutions containing protein. All buffers were prepared with Milli-Q water and filtered through 0.22 μm filters.

Reactions were monitored by TLC or LCMS. TLC analyses were performed on Merck TLC Silica gel 60 $F_{254}$ glass plates (0.2 mm). LCMS analyses of samples were performed using a Waters Acquity H-class UPLC coupled with a single quadrupole Waters SQD2. An Acquity UPLC CSH C18 Column, 130Å, 1.7 μm, 2.1 mm x 50 mm was used as the UPLC column.

Purification of compounds by silica column chromatography were performed using an automated system (Combiflash® Rf+) with prepackaged silica cartridges (25 μm or 50 μm PuriFlash® columns). $^{1}H$ and $^{13}C$ NMR spectra were recorded using a Bruker 600 MHz Avance 600 BBI spectrometer, a 500 MHZ Acance III Smart Probe spectrometer, or a 400 MHz Avance III HD Smart Probe spectrometer at 298.0 ± 0.1 K. Residual solvent peaks were used as an internal standard for calibration. All chemical shifts are quoted in ppm on the δ scale and the coupling constants are expressed in Hz. Signal splitting patterns are described as a singlet (s), broad singlet (br s), doublet (d), triplet (t), quartet (q), or multiplet (m).

UV-vis spectra were collected on an Agilent Cary 60 UV-vis spectrophotometer controlled by Cary WinUV software. Fluorescence spectroscopic data were recorded using an Agilent Cary Eclipse Fluorescence Spectrophotometer controlled by Cary WinUV software, and equipped with a Cary Eclipse Automated Polarizer for anisotropy measurements.

#### Small Molecule Synthesis

**2-(4-(dimethylamino)phenyl)benzo[*d*]thiazol-6-ol** (**S1**)

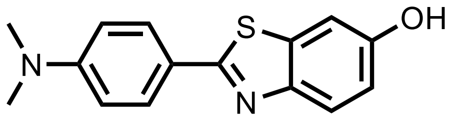


**S1** was prepared as previously reported.[1]

**6,6'-((1,4-phenylenebis(methylene))bis(oxy))bis(2-(4-(dimethylamino)phenyl)-3-methylbenzo[*d*]thiazol-3-ium) diiodide** (**1**)

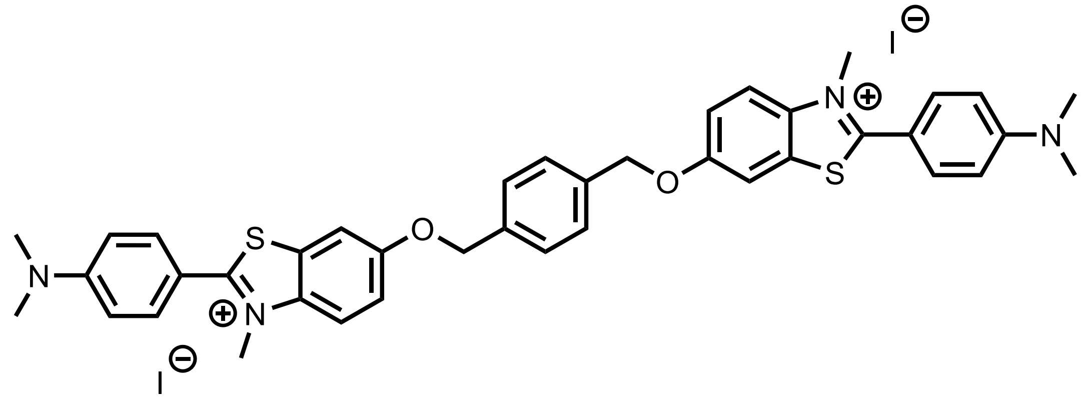

To **S1** (216 mg, 0.80 mmol, 2.0 equiv.) in DMF (5 mL) was added sodium hydride (60% in mineral oil, 29 mg, 1.20 mmol, 3.0 equiv.) in one portion at room temperature. The resultant yellow suspension was stirred for 1 h, producing a green solution. 1,4-*bis*(bromomethyl)benzene (102 mg, 0.39 mmol, 0.98 eq.) was then added, and the mixture heated to 90 °C for 19 h. After cooling to room temperature, ice water (15 mL) was added with vigorous stirring. The resultant precipitate was collected by filtration and washed with ice water (10 mL) then ice-cold ethanol (10 mL) to afford a poorly soluble intermediate (77.9 mg).

A mixture of the intermediate (19.3 mg, 30 μmol) and iodomethane (0.5 mL) in nitrobenzene (2.0 mL) was heated to 110 °C for 4 h under microwave irradiation. The reaction was cooled to room temperature and triturated using diethyl ether on ice. The precipitate was collected by filtration and washed with ice-cold diethyl ether (50 mL) to afford **1** as a yellow solid (28.0 mg, 30 μmol, 100%).

**Yield:** 28.0 mg (30 μmol, 100%)

**Aspect:** yellow solid

**$^1$H NMR (400 MHz, DMSO-*d*$_6$) δ (ppm):** 8.17 (d, *J* = 9.2 Hz, 2H), 8.07 (s, 2H), 7.79 (d, *J* = 8.7 Hz, 4H), 7.60 – 7.48 (m, 6H), 6.97 (d, *J* = 8.7 Hz, 4H), 5.29 (s, 4H), 4.21 (s, 6H), 3.12 (s, 12H).

**$^{13}$C NMR (176 MHz, DMSO-*d*$_6$) δ (ppm):** 171.5, 157.7, 153.5, 137.1, 136.1, 132.1, 129.7, 128.2, 118.6, 117.9, 112.0, 111.0, 107.8, 69.9, 39.7, 38.2.

**HRMS (ESI+):** 336.5048 m/z: Calculated for $C_{40}H_{40}N_4O_2S_2^{2+}$ = 336.5308 $[M]^{2+}$.

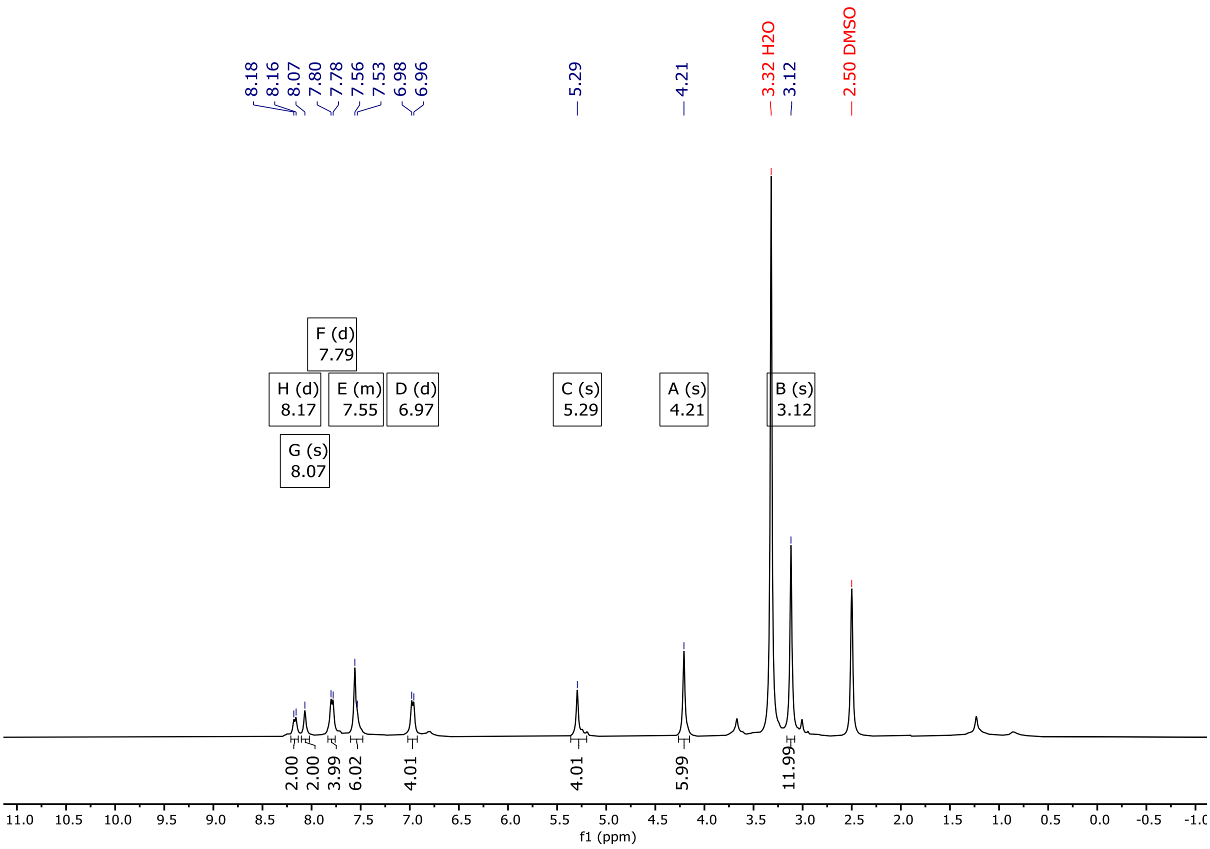


***Figure S1.*** $^{1}$H NMR (400 MHz, DMSO-$d_6$) spectra of **1**.

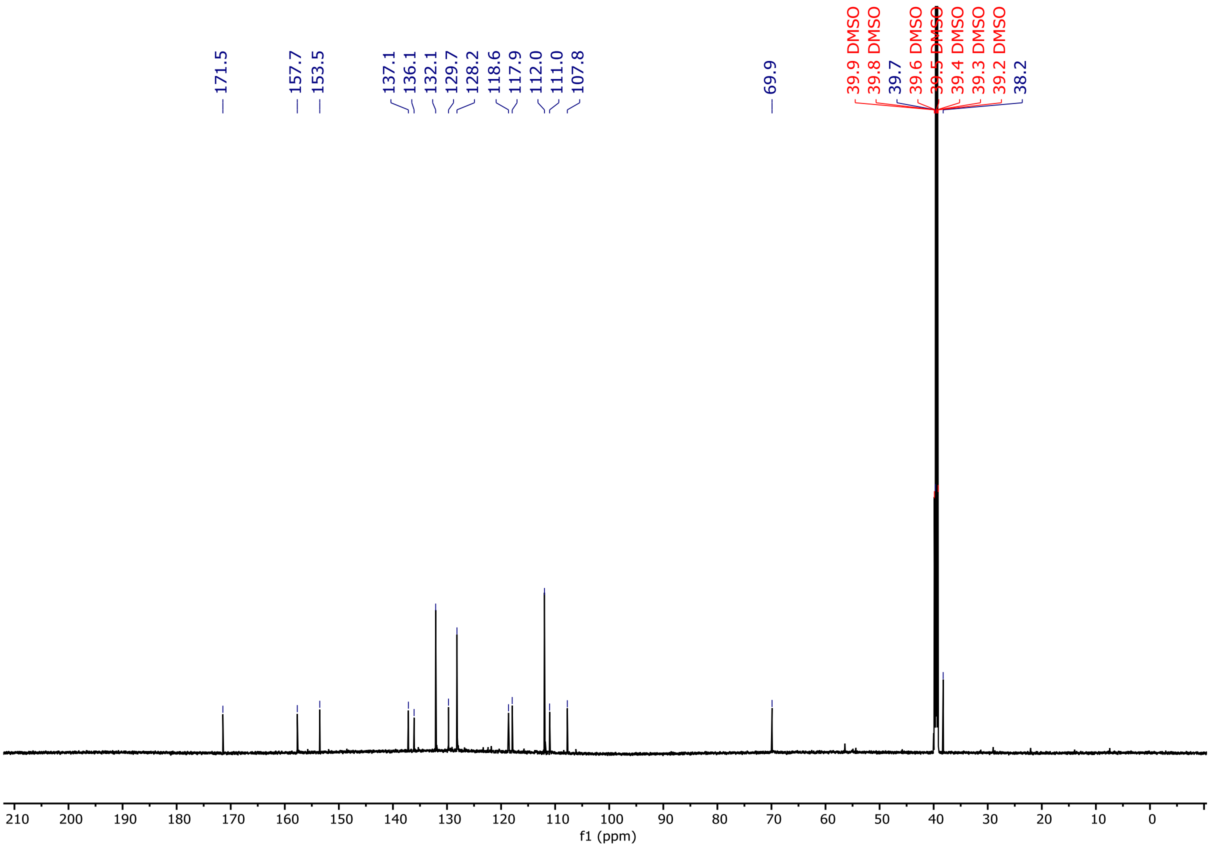


***Figure S2.*** $^{13}$C NMR (176 MHz, DMSO-$d_6$) spectra of **1**.

**2,7-bis(2-(2-(2-hydroxyethoxy)ethoxy)ethyl)benzo[lmn][3,8]phenanthroline-1,3,6,8(2*H*,7*H*)-tetraone** (**S2**)

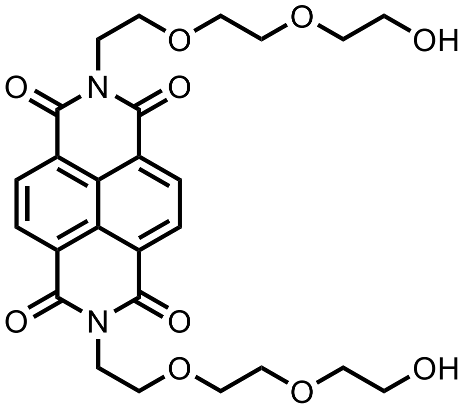


1,4,5,8-Naphthalenetetracarboxylic dianhydride (805 mg, 3.0 mmol, 1.0 equiv.) and 2-(2-aminoethoxy)ethanol (1.66 mL, 12.0 mmol, 4.0 equiv.) in NMP (9 mL) were heated to 110 °C for 2 h. The reaction mixture was cooled, poured into water, and **S2** was collected by filtration as a brown solid.

**Yield:** 1.451 g (2.73 mmol, 91%)

**Aspect:** brown solid

**$^1$H NMR (400 MHz, $CDCl_3$) δ (ppm):** 8.75 (s, 4H), 4.47 (t, *J* = 5.8 Hz, 4H), 3.87 (t, *J* = 5.7 Hz, 4H), 3.72 – 3.69 (m, 4H), 3.64 – 3.60 (m, 8H), 3.54 – 3.50 (m, 4H).

**$^{13}$C NMR (101 MHz, $CDCl_3$) δ (ppm):** 163.1, 131.2, 126.9, 126.7, 72.5, 70.6, 70.3, 68.0, 61.8, 39.8.

**HRMS (ESI+):** 531.2298 m/z: Calculated for $C_{26}H_{31}N_2O_{10}^+$ = 531.2174 [M+H]$^+$.

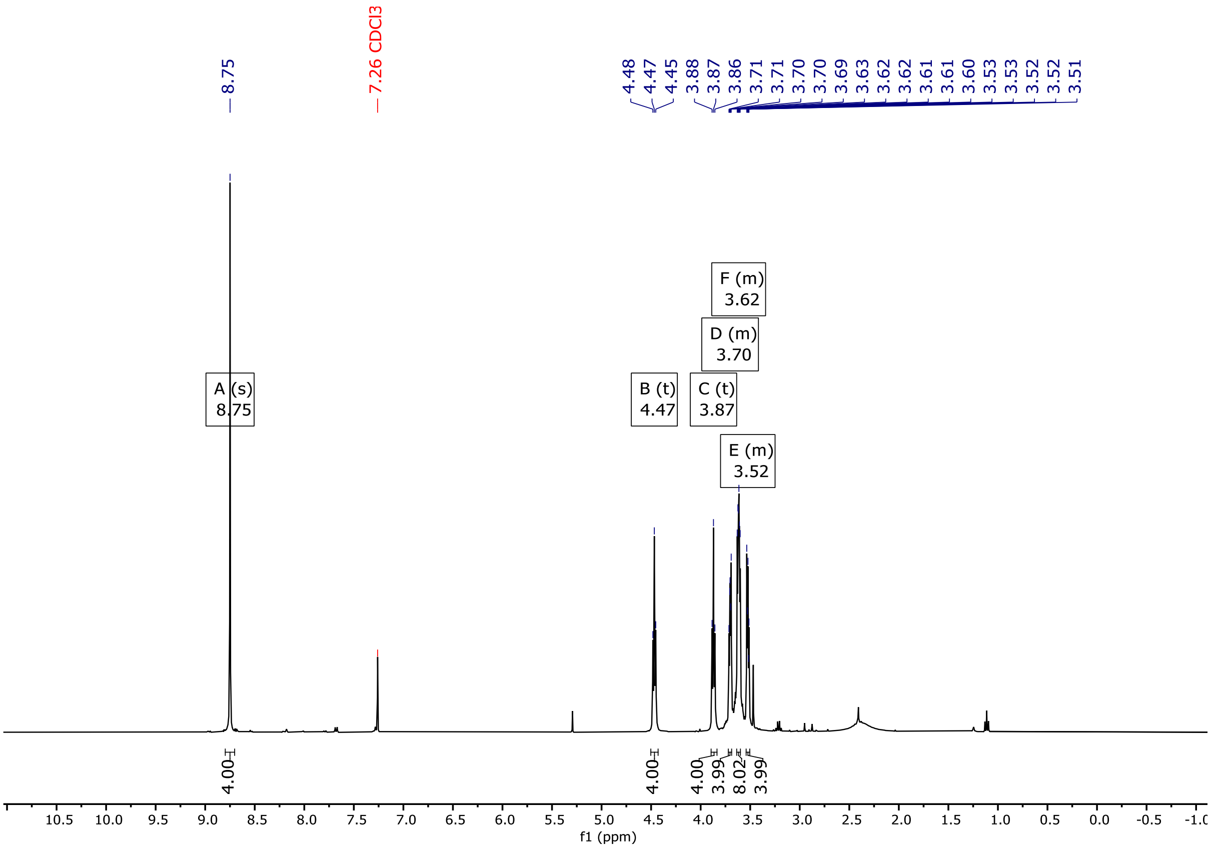


***Figure S3.*** $^1$H NMR (400 MHz, $CDCl_3$) spectra of **S2**.

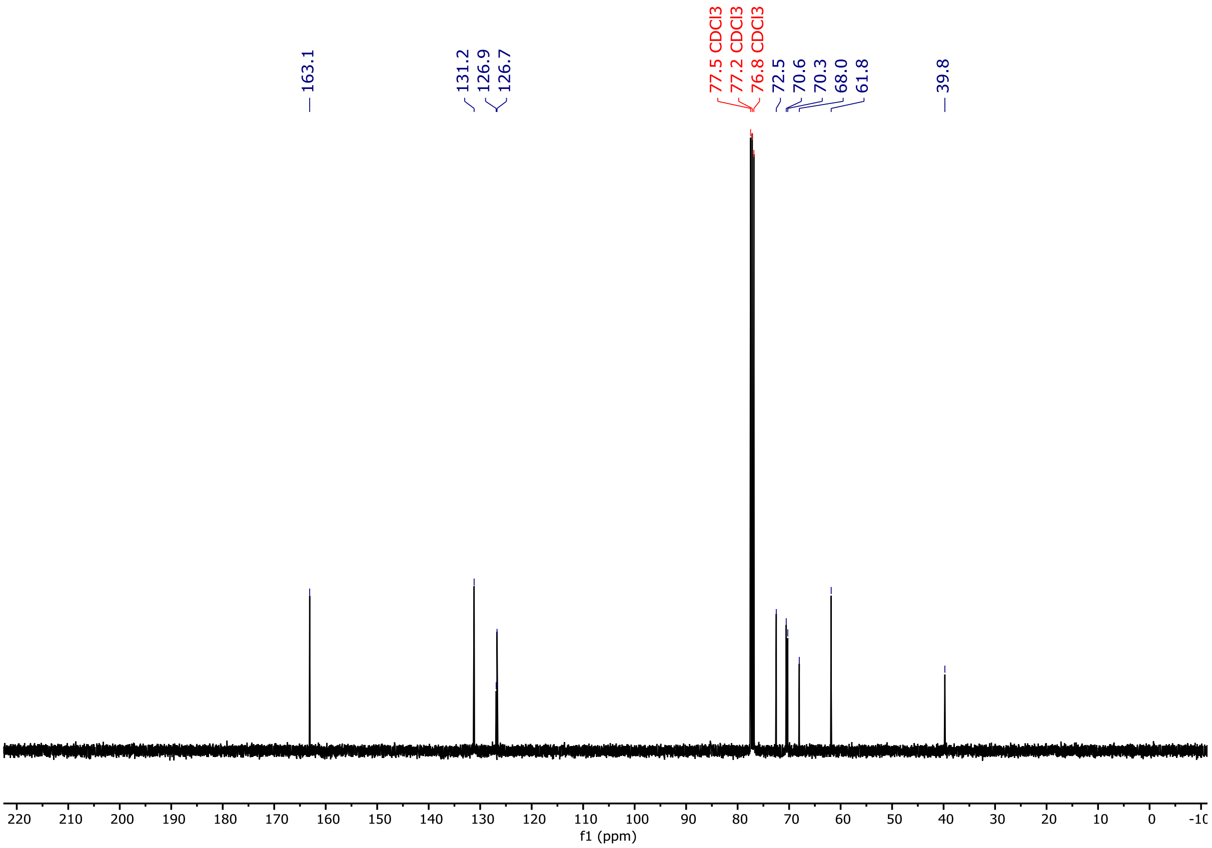


***Figure S4.*** $^{13}$C NMR (101 MHz, $CDCl_3$) spectra of **S2**.

**2-(2-(2-(7-(2-(2-(2-hydroxyethoxy)ethoxy)ethyl)-1,3,6,8-tetraoxo-3,6,7,8-tetrahydrobenzo[lmn][3,8]phenanthrolin-2(1*H*)-yl)ethoxy)ethoxy)ethyl 4-methylbenzenesulfonate** (**S3**)

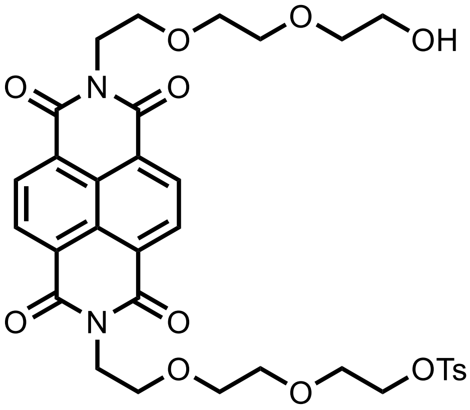


To a mixture of DMAP (2 mg, catalytic) and **S2** (212.2 mg, 0.40 mmol, 1.0 equiv.) in $CH_2Cl_2$ (2 mL) was added a solution of tosyl chloride (91.5 mg, 0.48 mmol, 1.2 equiv.) and triethylamine (223 µL, 1.6 mmol, 4.0 equiv.) in $CH_2Cl_2$ (2 mL) dropwise on ice. The reaction mixture was stirred for 1 h, and upon completion by LCMS, 10w/v% $NH_4Cl$ (10 mL) was added and the mixture was warmed to room temperature. The aqueous layer was washed with $CH_2Cl_2$ (3 x 20 mL) and the combined organic extracts were washed with brine (20 mL), dried over $Na_2SO_4$, and the solvent removed *in vacuo*. The crude product was then purified by flash silica column chromatography (PE/EtOAc) to afford the monotosylated **S4** (54.9 mg, 80 µmol, 20%) as a yellow-orange solid, and the ditosylated product (177.3 mg, 211 µL, 53%).

**Yield:** 54.9 mg (80 µmol, 20%)

**Aspect:** yellow-orange solid

**$^1$H NMR (400 MHz, $CDCl_3$) δ (ppm):** 8.71 (d, *J* = 1.3 Hz, 4H), 7.72 (dt, *J* = 8.4, 1.8 Hz, 2H), 7.32 – 7.29 (m, 2H), 4.43 (dt, *J* = 13.3, 5.8 Hz, 4H), 4.05 – 4.01 (m, 2H), 3.86 (t, *J* = 5.8 Hz, 2H), 3.80 (t, *J* = 5.8 Hz, 2H), 3.71 – 3.67 (m, 2H), 3.63 – 3.58 (m, 8H), 3.52 (dt, *J* = 5.9, 3.2 Hz, 4H), 2.42 (s, 3H).

**$^{13}$C NMR (101 MHz, $CDCl_3$) δ (ppm):** 163.0, 162.9, 144.9, 133.0, 131.1, 131.1, 129.9, 128.0, 126.8, 126.8, 126.7, 126.6, 72.5, 70.8, 70.5, 70.2, 70.2, 69.2, 68.7, 68.0, 67.9, 61.8, 39.7, 39.6, 21.7.

**HRMS (ESI+):** 685.2063 m/z: Calculated for $C_{33}H_{37}N_2O_{12}S^+$ = 685.2062 $[M+H]^+$. 707.1892 m/z: Calculated for $C_{33}H_{36}N_2O_{12}SNa^+$ = 707.1882.

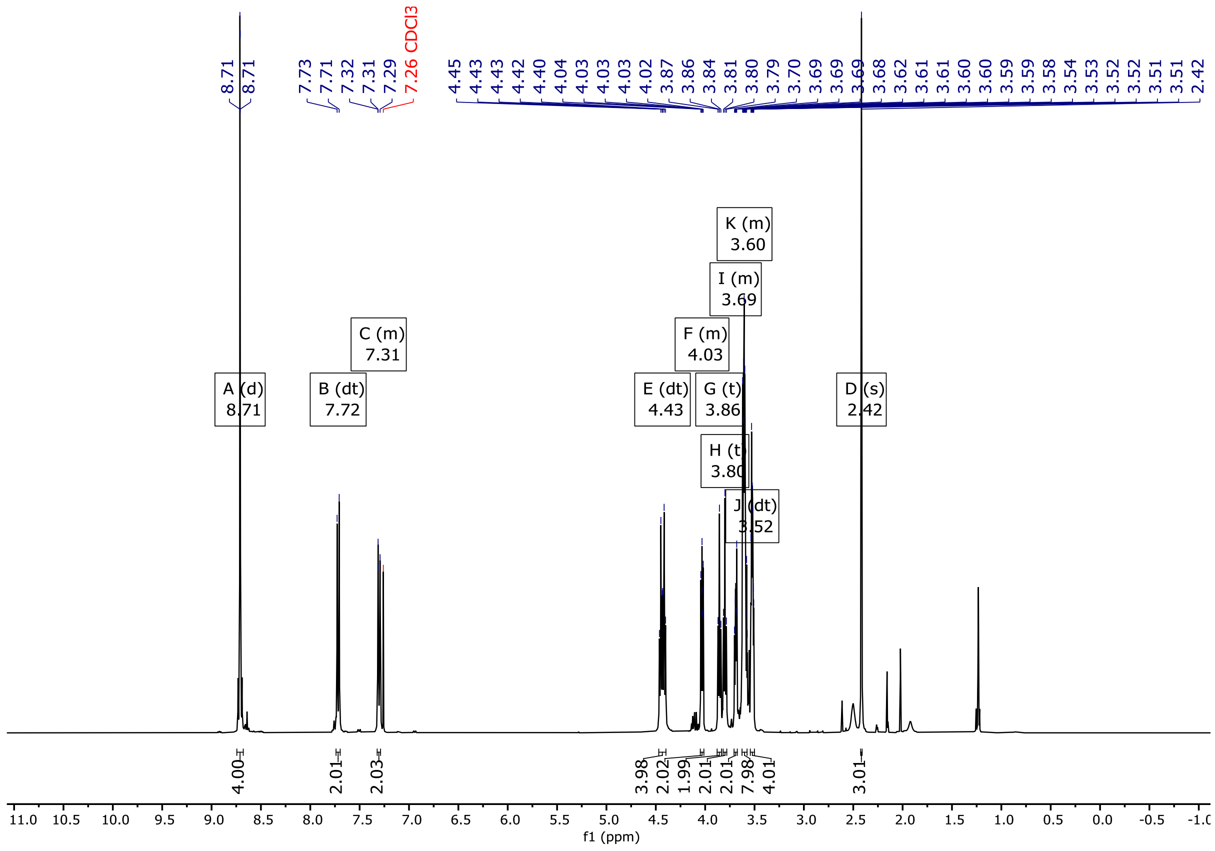

***Figure S5.*** $^{1}H$ NMR (400 MHz, $CDCl_3$) spectra of **S3**.

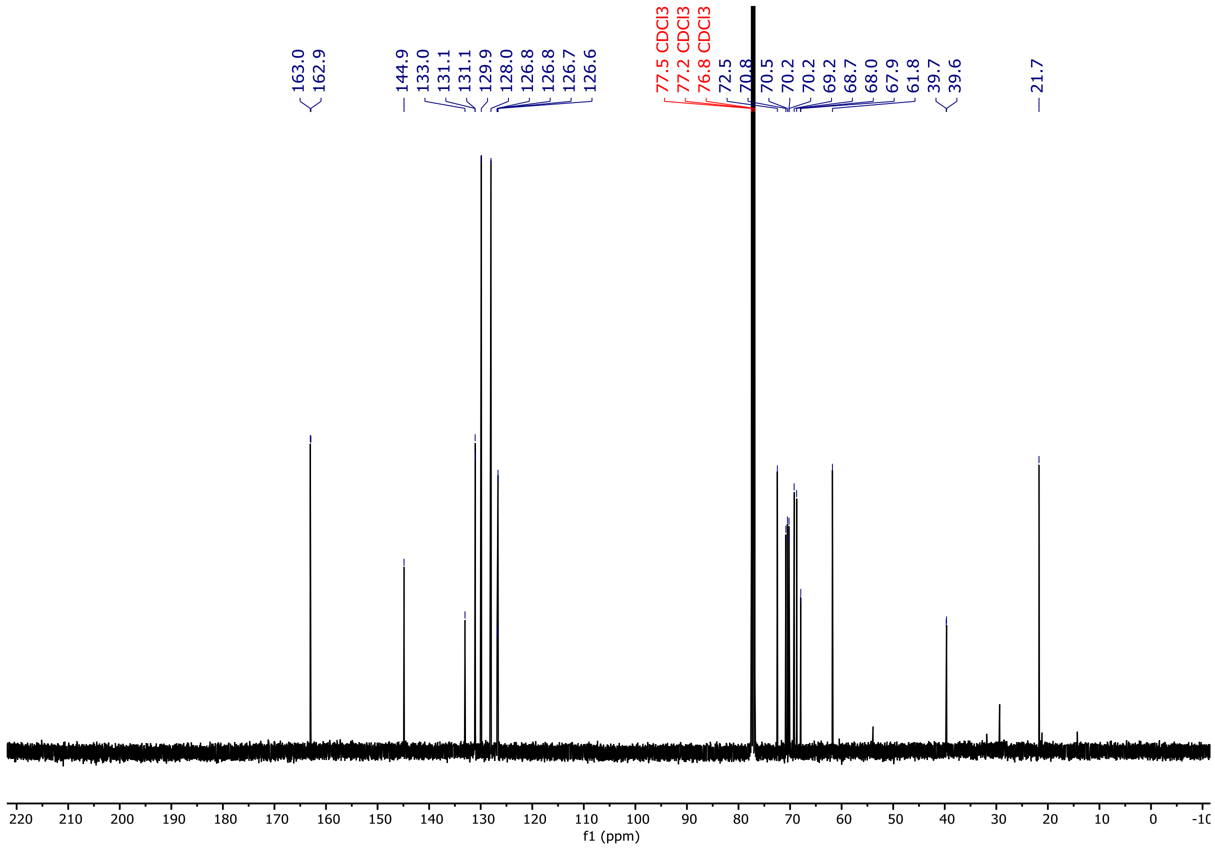

***Figure S6.*** $^{13}C$ NMR (101 MHz, $CDCl_3$) spectra of **S3**.

**2-(2-(2-(2-((2-(4-(dimethylamino)phenyl)benzo[*d*]thiazol-6-yl)oxy)ethoxy)ethoxy)ethyl)-7-(2-(2-(2-hydroxyethoxy)ethoxy)ethyl)benzo[lmn][3,8]phenanthroline-1,3,6,8(2*H*,7*H*)-tetraone** (**S4**)

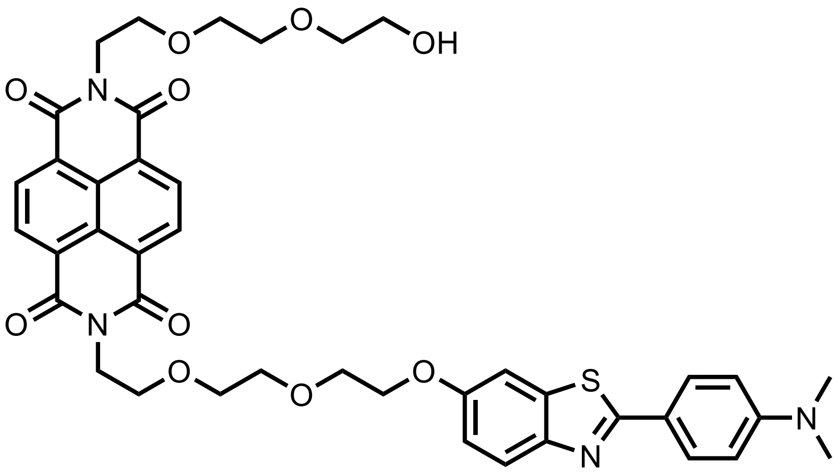


To **S1** (23.0  mg, 85 μmol, 1.1 equiv.) and sodium hydride (60% in mineral oil, 8.2 mg, 4.4 equiv.) in DMF (0.8 mL) was added **S3** (52.9 mg, 77 μmol, 1.0 equiv.) in DMF (0.7 mL) on ice. The reaction was warmed to room temperature and stirred for 30 min, producing a green solution that was then heated to 80 °C for 17 h. The residue was diluted in $CH_2Cl_2$ (50 mL) and the organic layer was washed with 1 M KOH (3 x 20 mL), brine (20 mL), and dried over $Na_2SO_4$. The crude residue was then purified by flash silica column chromatography ($CH_2Cl_2$/MeOH) to afford **S5** as a yellow solid (25.5 mg, 33 μmol, 43%).

**Yield:** 25.5 mg (33 μmol, 43%)

**Aspect:** yellow solid

**$^1$H NMR (400 MHz, $CDCl_3$) δ (ppm):** 8.64 (s, 4H), 7.78 (d, *J* = 8.9 Hz, 2H), 7.67 (d, *J* = 8.9 Hz, 1H), 7.13 (d, *J* = 2.5 Hz, 1H), 6.86 (dd, *J* = 8.9, 2.6 Hz, 1H), 6.69 (dt, *J* = 8.9, 2.2 Hz, 2H), 4.40 (t, *J* = 6.0 Hz, 2H), 4.34 (t, *J* = 5.8 Hz, 2H), 4.03 (t, *J* = 4.8 Hz, 2H), 3.88 – 3.78 (m, 6H), 3.74 – 3.70 (m, 4H), 3.67 – 3.65 (m, 2H), 3.63 – 3.58 (m, 4H), 3.57 – 3.48 (m, 2H), 3.04 (s, 6H).

**$^{13}$C NMR (101 MHz, $CDCl_3$) δ (ppm):** 162.9, 156.2, 152.0, 149.0, 135.6, 130.9, 128.5, 122.6, 121.5, 115.4, 111.8, 105.5, 77.4, 72.5, 70.9, 70.6, 70.5, 70.2, 69.9, 68.2, 61.8, 40.3, 29.8.

**HRMS (ESI+):** 783.2698 m/z: Calculated for $C_{41}H_{43}N_4O_{10}S^+$ = 783.2695 [M+H]$^+$. 805.2520 m/z: Calculated for $C_{41}H_{42}N_4O_{10}SNa^+$ = 805.2519.

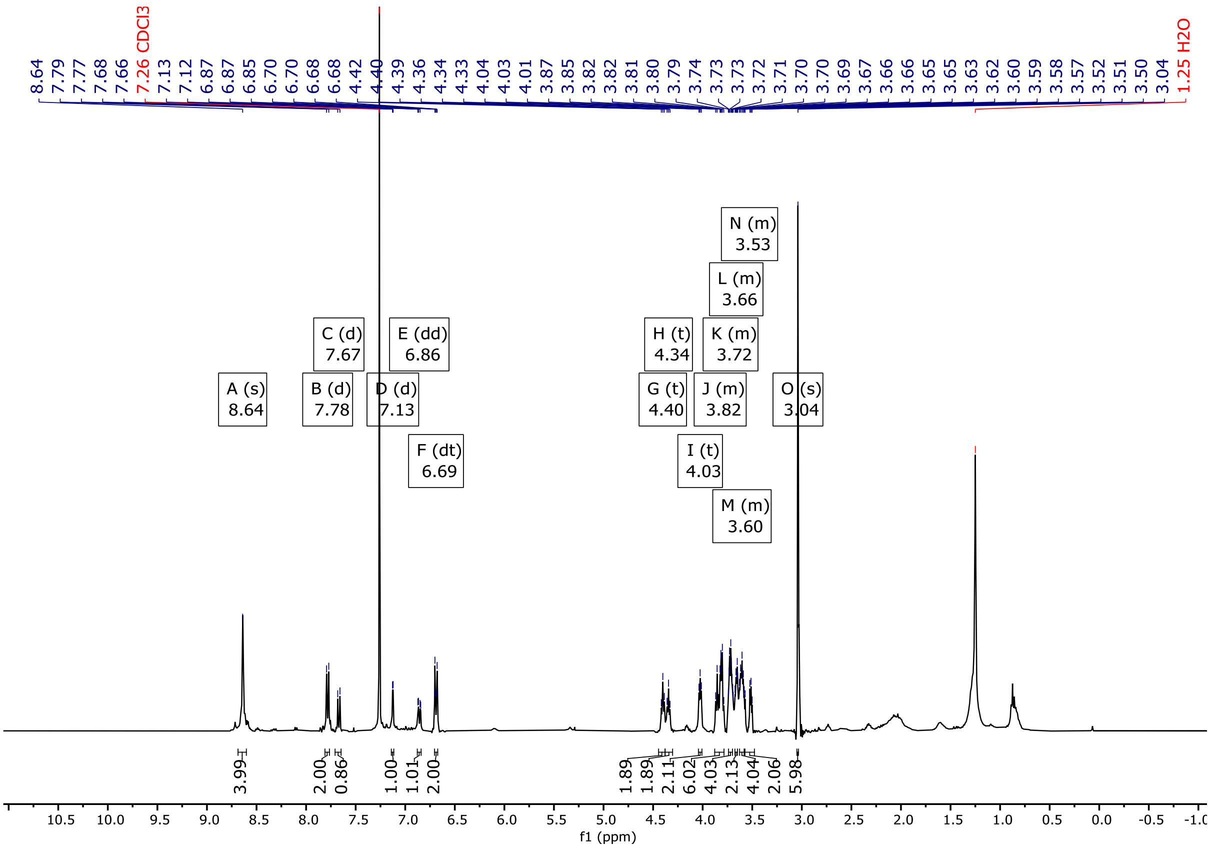


***Figure S7.*** $^{1}$H NMR (400 MHz, $CDCl_3$) spectra of **S4**.

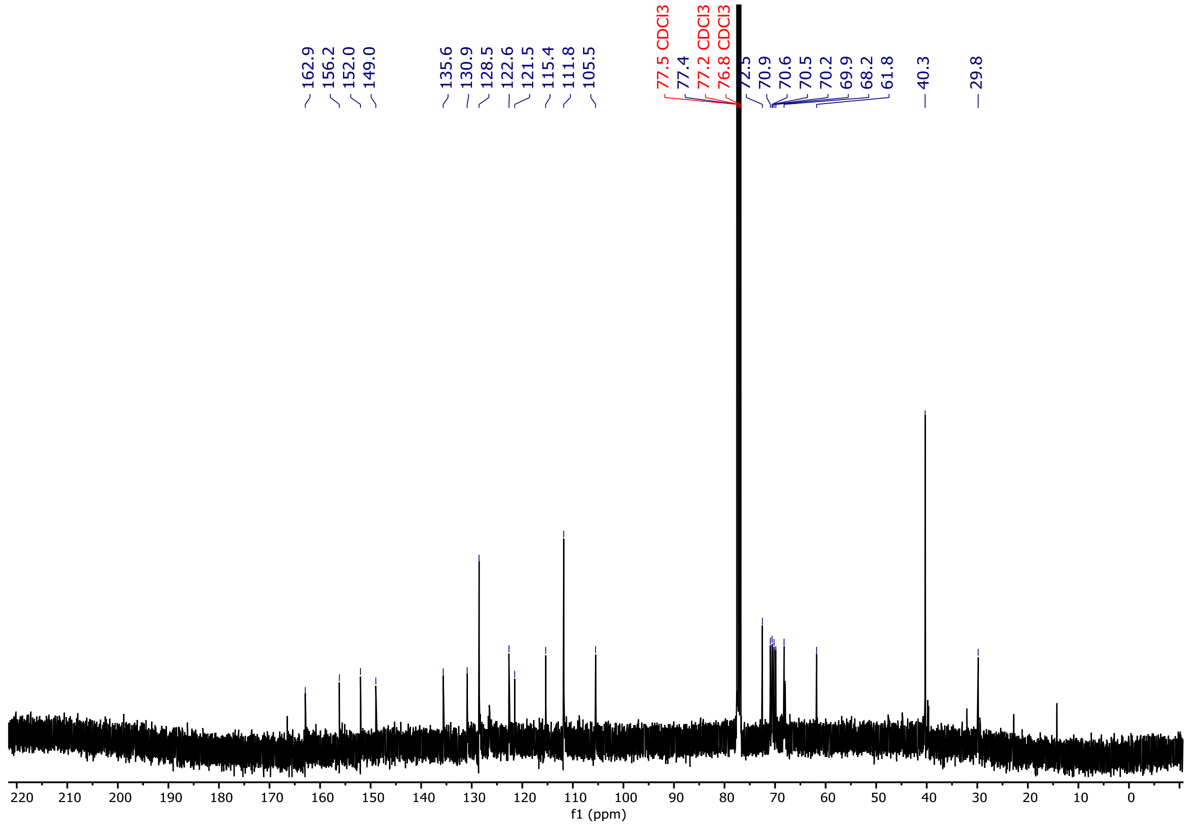


***Figure S8.*** $^{13}$C NMR (101 MHz, $CDCl_3$) spectra of **S4**.

**2-(4-(dimethylamino)phenyl)-6-(2-(2-(2-(7-(2-(2-(2-hydroxyethoxy)ethoxy)ethyl)-1,3,6,8-tetraoxo-3,6,7,8-tetrahydrobenzo[lmn][3,8]phenanthrolin-2(1*H*)-yl)ethoxy)ethoxy)ethoxy)-3-methylbenzo[*d*]thiazol-3-ium iodide** (**2**)

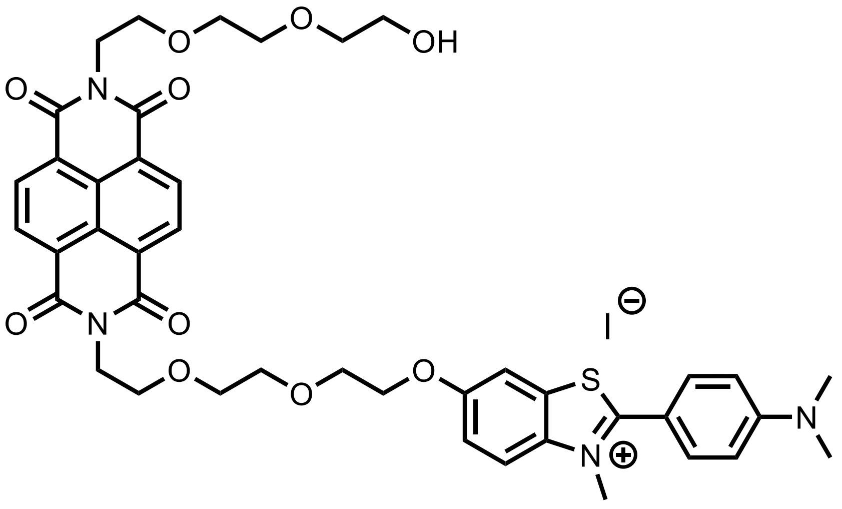


A mixture of **S4** (23.5 mg, 30 μmol) and iodomethane (0.5 mL) in nitrobenzene (2.0 mL) was heated to 110 °C for 5 h under microwave irradiation. The reaction was cooled to room temperature and triturated using diethyl ether on ice. The precipitate was collected by filtration and washed with ice-cold diethyl ether (50 mL) to afford **2** as a brown wax (26.6 mg, 28 μmol, 93%).

**Yield:** 26.6 mg (28 μmol, 93%)

**Aspect:** brown wax

**$^1$H NMR (400 MHz, DMSO) δ (ppm):** 8.62 – 8.57 (m, 4H), 8.05 (d, *J* = 9.4 Hz, 1H), 7.80 – 7.76 (m, 3H), 7.33 (dd, *J* = 9.3, 2.2 Hz, 1H), 6.96 (d, *J* = 8.9 Hz, 2H), 4.51 (t, *J* = 5.3 Hz, 2H), 4.25 (t, *J* = 6.2 Hz, 2H), 4.20 – 4.18 (m, 5H), 4.10 – 4.04 (m, 2H), 3.76 – 3.72 (m, 4H), 3.65 (t, *J* = 6.7 Hz, 2H), 3.62 – 3.60 (m, 3H), 3.57 – 3.50 (m, 2H), 3.49 – 3.45 (m, 2H), 3.37 (t, *J* = 5.0 Hz, 2H), 3.11 (s, 6H).

**$^{13}$C NMR (176 MHz, DMSO) δ (ppm):** 171.2, 162.5, 162.5, 157.8, 153.5, 136.8, 132.0, 130.5, 130.4, 129.6, 126.1, 126.0, 126.0, 118.2, 117.7, 112.0, 112.0, 110.9, 107.1, 72.3, 69.8, 69.6, 69.6, 68.6, 68.1, 66.7, 60.1.

**HRMS (ESI+):**

797.2993 m/z: Calculated for $C_{42}H_{45}N_4O_{10}S^+$ = 797.2851 $[M+H]^+$.

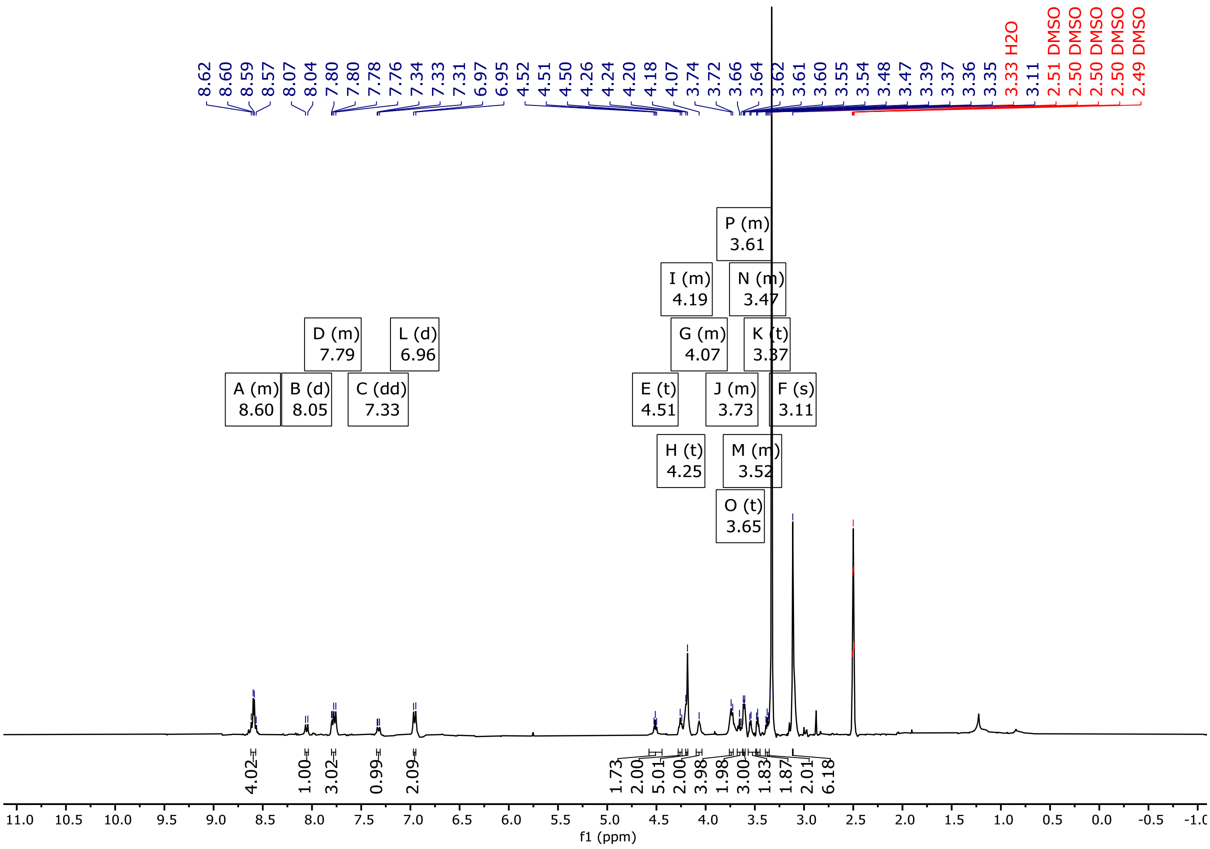

***Figure S9.*** $^{1}H$ NMR (400 MHz, DMSO-$d_6$) spectra of **S4**.

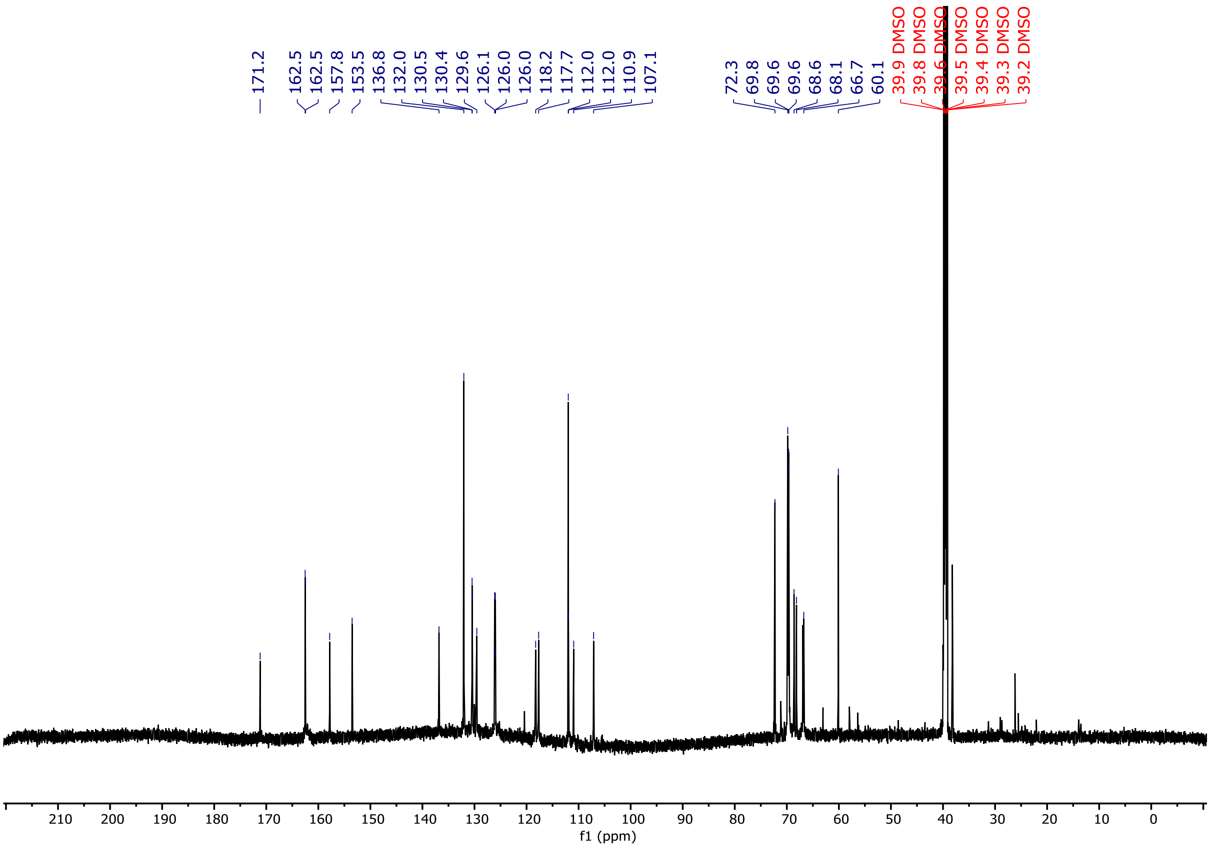

***Figure S10***. $^{13}C$ NMR (176 MHz, DMSO-$d_6$) spectra of **S4**.

#### UV-Visible Characterisation

Ligand stock solutions in DMSO (10 mM) were diluted in ethanol to obtain a 50 µM solution in a quartz fluorescence cuvette (Hellma Analytics, 1 cm pathlength). UV-visible spectra were obtained using an Agilent Cary 60 UV-vis spectrophotometer with Cary WinUV software using a scan rate of 600 nm/min, a data interval of 1.0 nm and an averaging time of 0.10 at 25°C.

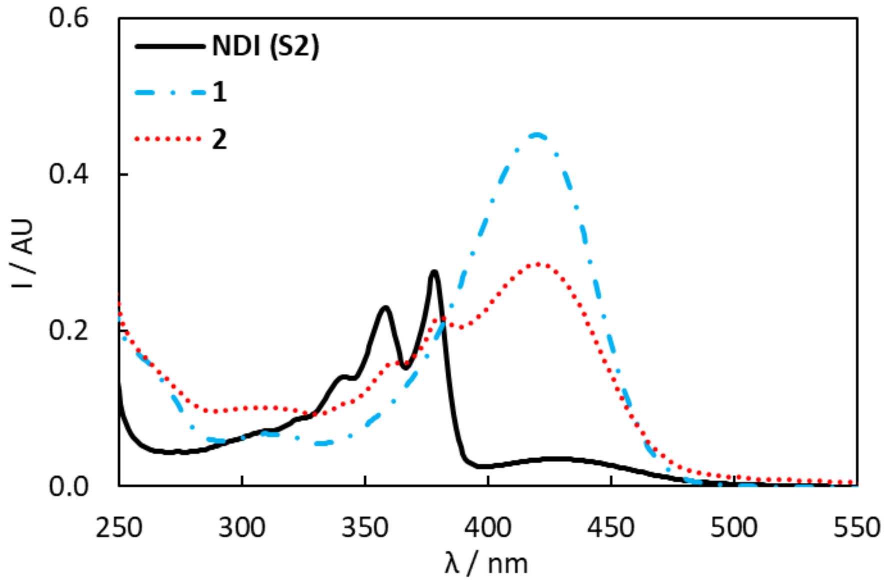


***Figure S11**. UV-vis spectra of **S2** (black line), **1** (blue dash-dotted line), and **2** (red dotted line) (50 µM in EtOH).*

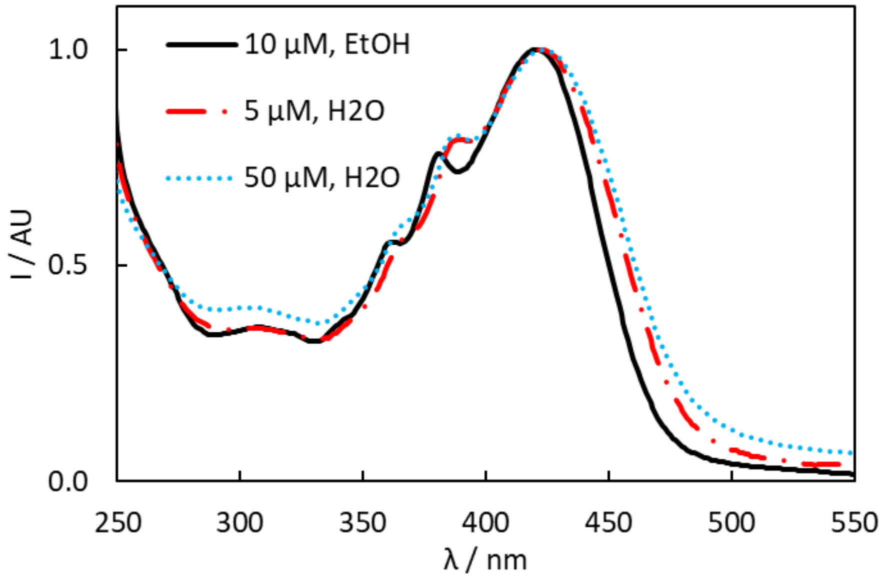


***Figure S12**. UV-vis spectra of **2** in EtOH (black line, 50 µM in EtOH), and $H_2O$ (red dash-dotted line: 5 µM; blue dotted line: 50 µM).*

#### Fluorescence Characterisation

Fluorescence excitation and emission spectra were obtained using an Agilent Cary Eclipse Fluorescence Spectrophotometer with a scan rate of 600 nm/min, a data interval of 1.0 nm, an averaging time of 0.10. 20 nm excitation and emission slits, and medium PMT voltage at 25°C.

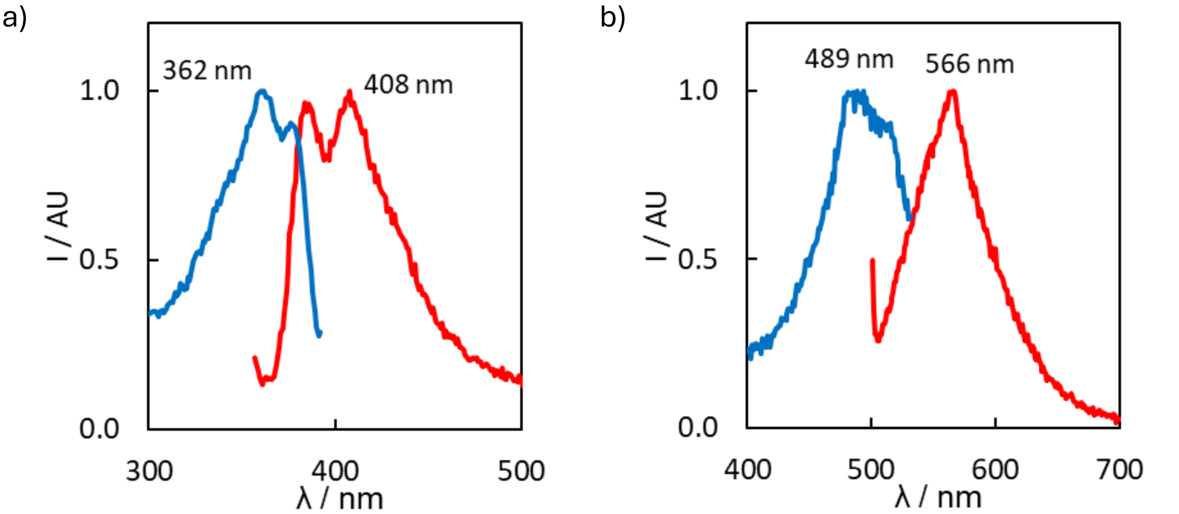


***Figure S13***. Fluorescence spectra of **S2** (10 μM) in EtOH (a: $\lambda_{ex}$ = 342, $\lambda_{em}$ = 407; b: $\lambda_{ex}$ = 486, $\lambda_{em}$ = 557).

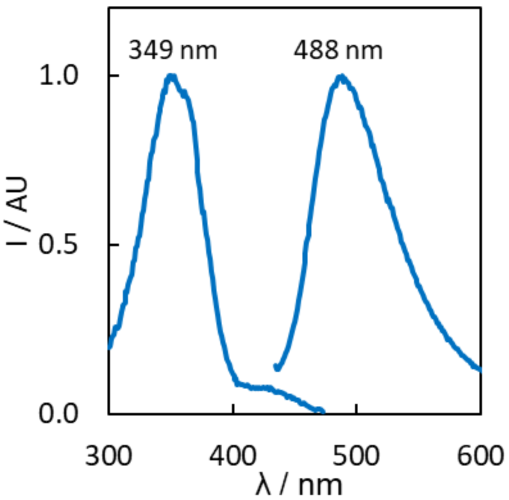


***Figure S14***. Fluorescence spectra of **1** (10 μM) in EtOH ($\lambda_{ex}$ = 440, $\lambda_{em}$ = 488).

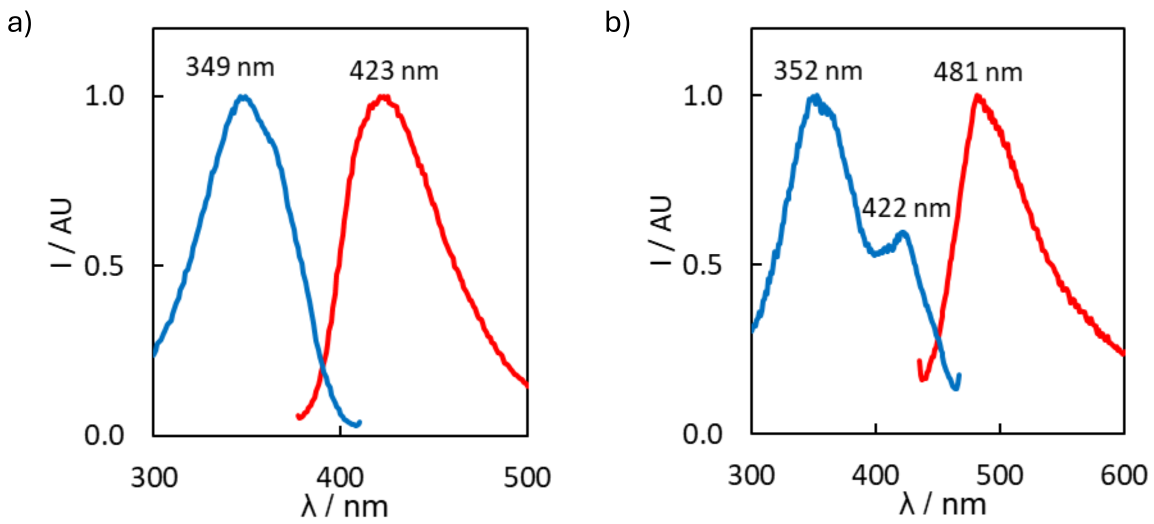


***Figure S15***. Fluorescence spectra of **2** (10 μM) in EtOH (a: $\lambda_{ex}$ = 362, $\lambda_{em}$ = 425; b: $\lambda_{ex}$ = 420, $\lambda_{em}$ = 482).

#### ThT Hill Plots

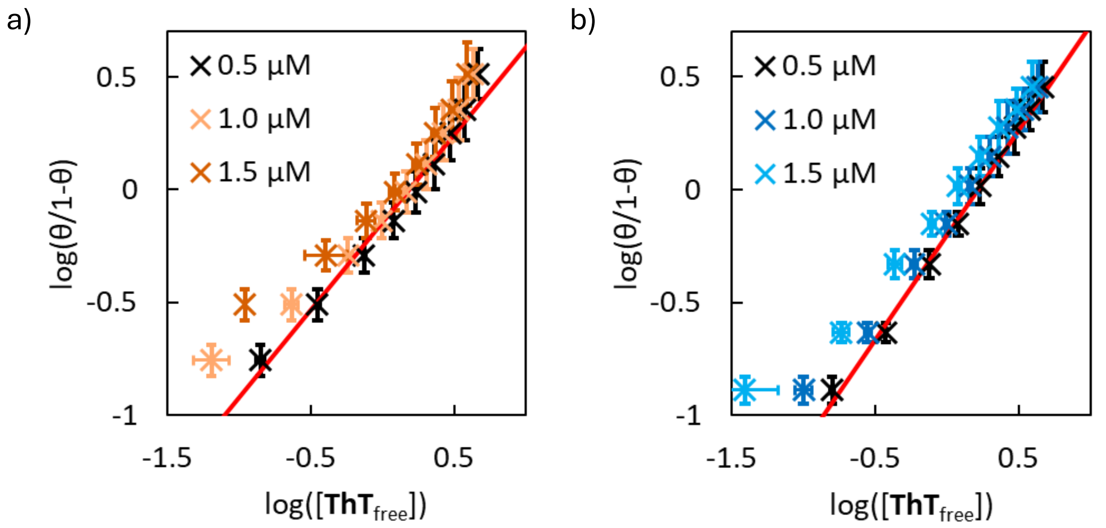


***Figure S16***. Hill plots for binding assays of **ThT** ($\lambda_{ex}$ = 440 nm) with (a) αSyn-Tris and αSyn-PBS fibrils (500 nM) in 1xPBS (pH 7.4). Red line shows weighted linear regression to 0.5 μM data (a: y = (0.8 ± 0.1)x - (0.14 ± 0.06); b: y = (0.9 ± 0.1)x - (0.20 ± 0.05), with errors corresponding to 95% confidence intervals of the fit). Data points are the average of at least three experimental measurements with 95% confidence intervals shown.

#### Preparation of αSyn Fibrils

Wild-type human monomeric αSyn in 1xPBS (180 μM) expressed in E. Coli and purified as previously reported.2 A solution of monomeric αSyn was added to an Amicon Ultra-15 Centrifugal filter (15 kDa MWCO) and centrifuged (15 min, 4000 x g). The retained monomeric αSyn was washed with 5 mL of the aggregation buffer; either 1xPBS (pH 7.4) or Tris·HCl (50 mM, pH 7.5) and NaCl (100 mM), and centrifuged (15 min, 4000 x g). This wash step was repeated four times in total. The retained filtrate was diluted to 1 mL using the aggregation buffer and incubated at 37 °C for 72 h with gentle agitation by a magnetic stir bar in an Eppendorf LoBind microcentrifuge tube (2.0 mL). The resultant fibrils were then pelletised in a centrifuge (15 min, 4000 x g), the supernatant removed, and the fibrils gently resuspended in the desired buffer. The fibril preparation was then stored at -80 °C for 2 years. Characterisation data is in agreement with that previously reported.[3]

#### Binding Model Derivation

Throughout the derivation, $\alpha$ denotes a specific ligand type under consideration, whereas $i$ and $j$ index general ligand types appearing in summations and neighbouring configurations. This derivation is largely based on the procedure outlined by Villaluenga *et al.*,[4] and by McGhee and von Hippel.[5]

##### Cooperativity parameter

The cooperativity parameter is defined as the ratio of the probability of ligands existing in an adjacent configuration (B), to the probability of ligands existing in a non-adjacent configuration (A) (Figure S17),

$$\begin{aligned}\omega_{ij} &= \frac{\text{probability of configuration II}}{\text{probability of configuration I}}\\ &= \frac{(\text{prob. config. A})\left(b^i_{m_i} b^j_1\right)\left(b^j_1 b^j_2\right)\dots\left(b^j_{m_j} f\right)(ff)^{x+y-1}(\text{prob. config. B})}{(\text{prob. config. A})\left(b^i_{m_i} f\right)(ff)^{x-1}\left(f b^j_1\right)\left(b^j_1 b^j_2\right)\dots\left(b^j_{m_j} f\right)(ff)^{y-1}(\text{prob. config. B})}\\ &= \frac{\left(b^i_{m_i} b^j_1\right)(ff)}{\left(b^i_{m_i} f\right)\left(f b^j_1\right)}.\end{aligned} \qquad \text{Eq. S1}$$

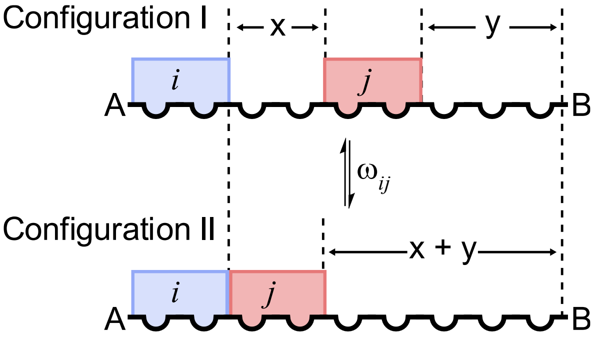


**Figure S17**. Definition of the cooperativity parameter $\omega$ used in this model.

##### Gap probability

A gap is defined as a sequence of $g$ consecutive free lattice residues. A ligand $i$ cannot bind to gaps smaller than their size ($g < m_i$), and $g$ = 0 indicates that two ligands are adjacent. The probability that a given gap between a bound ligand of type $i$ and a bound ligand of type $j$ is $g$ lattice residues long is given by

$$\left(P_g\right)_{ij} = \left(b^i_{m_i} f\right)(ff)^{g-1}\left(f b^j_1\right).$$

##### Computing the number of free binding sites

Derivation of the average number of free binding sites per lattice residue follows the procedure presented by Villaluenga *et al.*[4] The goal is to compute the average number of free ligand binding sites per lattice residue, $\bar{s}_\alpha$. There are three types of free binding sites:

an isolated site with no adjacent ligands ($\bar{s}_{iso}$), a singly contiguous site with one adjacent ligand ($\bar{s}_{sc}$), and a doubly contiguous site with two adjacent ligands ($\bar{s}_{dc}$). Following the notation used in Figure S17, if a ligand binds to an isolated site with affinity $K_i$, the affinity for a single contiguous site is $K_i\omega_{ij}$, and the affinity for a doubly contiguous site is $K_i\omega_{ij}^2$.

The number of binding sites in a gap of $g$ lattice residues between two bound ligands is given by $g - m_\alpha + 1$ if $g \geq m_\alpha$, but zero if the gap is smaller than $\alpha$. The probability of a gap being $g$ free lattice residues long is denoted by $(P_g)$, and is related to $\left(P_g\right)_{ij}$ by the sum

$$(P_g) = \sum_{i=1}^{T}\sum_{j=1}^{T}\left(P_g\right)_{ij}. \quad \text{Eq. S2}$$

Consider a specific ligand $\alpha$. For doubly contiguous sites, the gap size $g$ must be exactly equal to the length of the ligand, $m_\alpha$. The average number of doubly contiguous binding sites for ligand $\alpha$ in a gap flanked by ligand $i$ on the left and ligand $j$ on the right, $(\bar{s}_{dc})_{ij}^{\alpha}$, is given by

$$(\bar{s}_{dc})_{ij}^{\alpha} = \left(P_{m_i}\right)_{ij} = \left(b_{m_i}^{i} f\right)(ff)^{m_\alpha - 1}\left(f b_1^{j}\right). \quad \text{Eq. S3}$$

By summing over different combinations of ligands $i$ and $j$, weighted by the respective cooperativities, the average number of doubly contiguous binding sites for ligand $\alpha$ $(\bar{s}_{dc})^{\alpha}$ is obtained as

$$\begin{aligned}(\bar{s}_{dc})^{\alpha} &= \sum_{i=1}^{T}\sum_{j=1}^{T}\left(P_{m_\alpha}\right)_{ij}\omega_{\alpha i}\omega_{j\alpha}, \\ &= \sum_{i=1}^{T}\sum_{j=1}^{T}\left(b_{m_i}^{i} f\right)(ff)^{m_\alpha - 1}\left(f b_1^{j}\right)\omega_{\alpha i}\omega_{j\alpha}, \\ &= \left[\sum_{i=1}^{T}\frac{\theta_i}{\theta}\left(b_{m_i}^{i} f\right)\omega_{\alpha i}\right](ff)^{m_\alpha - 1}\left[\sum_{j=1}^{T}\left(f b_1^{j}\right)\omega_{j\alpha}\right]. \quad \text{Eq. S4}\end{aligned}$$

For $g \geq m_\alpha + 1$, there are two singly contiguous binding sites. The average number of free singly contiguous binding sites for a specific ligand $\alpha$ flanked by ligand $i$ on the left and ligand $j$ on the right, $(\bar{s}_{sc})_{ij}^{\alpha}$, is given by the sum

$$(\bar{s}_{sc})_{ij}^{\alpha} = 2\sum_{g=m_\alpha+1}^{T}\left(P_g\right)_{ij} = 2\frac{\theta_i}{\theta}\left(b_{m_i}^{i} f\right)\frac{(ff)^{m_\alpha}}{1-(ff)}\left(f b_1^{j}\right). \quad \text{Eq. S5}$$

By summing over different combinations of ligands $i$ and $j$, the average number of singly contiguous binding sites $(\bar{s}_{sc})^{\alpha}$ is obtained as

$$(\bar{s}_{sc})^{\alpha} = \sum_{i=1}^{T}\sum_{j=1}^{T}(\bar{s}_{sc})_{ij}^{\alpha}\frac{\omega_{i\alpha}+\omega_{\alpha j}}{2}$$
$$= \sum_{i=1}^{T}\sum_{j=1}^{T}\frac{\theta_i}{\theta}\left(b_{m_i}^{i}f\right)\frac{(ff)^{m_\alpha}}{1-(ff)}\left(fb_1^{j}\right)(\omega_{i\alpha}+\omega_{\alpha j}). \quad \text{Eq. S6}$$

Finally, for $g \geq m_\alpha + 2$, there are $(g - m_\alpha - 1)$ isolated binding sites. The sum then becomes

$$(\bar{s}_{iso})^{\alpha} = \sum_{i=1}^{T}\sum_{j=1}^{T}\sum_{g=m_\alpha+2}^{\infty}(g-m_\alpha-1)\left(P_g\right)_{ij}$$
$$= \left[\sum_{i=1}^{T}\frac{\theta_i}{\theta}\left(b_{m_i}^{i}f\right)\right]\frac{(ff)^{m_\alpha+1}}{\left(1-(ff)\right)^2}\left[\sum_{j=1}^{T}\left(fb_1^{j}\right)\right]. \quad \text{Eq. S7}$$

Finally, the total average number of free binding sites is expressed as

$$(\bar{s})_{\alpha} = (\bar{s}_{dc})^{\alpha} + (\bar{s}_{sc})^{\alpha} + (\bar{s}_{iso})^{\alpha}$$
$$= \left[\sum_{i=1}^{T}\frac{\theta_i}{\theta}\left(b_{m_i}^{i}f\right)\left[\omega_{i\alpha}+\frac{(ff)}{1-(ff)}\right]\right](ff)^{m_\alpha+1}\left[\sum_{j=1}^{T}\left(fb_1^{j}\right)\left[\omega_{\alpha j}+\frac{(ff)}{1-(ff)}\right]\right]. \quad \text{Eq. S8}$$

#### Final system of equations

The following final system of equations were used when generating synthetic data, or fitting experimental data, using the implemented mathematical model.

The first two equations used describe normalisation of the probability that only two types of sites can lie to the right of a free residue,

$$(ff) + \sum_{i=1}^{T}\left(fb_1^{i}\right) = 1, \quad \text{Eq. S9}$$

$$(1-\theta)(ff) + \sum_{i=1}^{T}\frac{\theta_i}{m_i}\left(b_{m_i}^{i}f\right) = 1-\theta. \quad \text{Eq. S10}$$

Next, the probability of each type of lattice residue lying to the right of a ligand must sum to unity,

$$\left(b_{m_i}^{i}f\right) + \sum_{j=1}^{T}\left(b_{m_i}^{i}b_1^{j}\right) = 1. \quad \text{Eq. S11}$$

Given the cooperativity parameter defines the probability of having two ligands adjacent,

$$\omega_{ij} = \frac{\left(b_{m_i}^{i} b_1^{j}\right)(ff)}{\left(b_{m_i}^{i} f\right)\left(f b_1^{j}\right)}, \qquad \text{Eq. S12}$$

then, for a specific ligand $\alpha$

$$\left(b_{m_\alpha}^{\alpha} f\right) + \sum_{j=1}^{T} \omega_{\alpha j} \frac{\left(b_{m_\alpha}^{\alpha} f\right)\left(f b_1^{j}\right)}{\left(b_{m_\alpha}^{\alpha} b_1^{j}\right)} = 1. \qquad \text{Eq. S13}$$

The mass action equation can then be implemented

$$\frac{[L_\alpha \cdot s]}{[L_\alpha]} = K_\alpha \bar{s}_\alpha \sum_{i=1}^{T} [L_i \cdot s]. \qquad \text{Eq. S14}$$

Note that for a system with only a single ligand, this reduces to

$$1 = K_\alpha \bar{s}_\alpha [L_\alpha] \qquad \text{Eq. S15}$$

Finally, the conditional probabilities and the concentration of each species can be linked. A bound ligand $\alpha$ either follows a free residue, with conditional probability $(f b_1^{\alpha})$, or the final residue of another ligand, with conditional probability $\left(b_{m_i}^{i} b_1^{\alpha}\right)$. Weighting these probabilities with the concentration of free lattice residue and the summed concentrations of bound ligands of type $i$ must yield the total concentration of a bound ligand of type $\alpha$, expressed as

$$\begin{aligned}[L_\alpha \cdot s] &= \left[r_{free}\right](f b_1^{\alpha}) + \sum_{i=1}^{T} [L_i \cdot s]\left(b_{m_i}^{i} b_1^{\alpha}\right) \\ &= \left[r_{free}\right](f b_1^{\alpha}) + \sum_{i=1}^{T} [L_i \cdot s] \omega_{i\alpha} \frac{\left(b_{m_i}^{i} f\right)(f b_1^{\alpha})}{(ff)},\end{aligned} \qquad \text{Eq. S16}$$

where $\left[r_{free}\right]$ is the concentration of free lattice residues.

Finally, conservation is enforced for the total concentration of each ligand by:

$$[L_\alpha] + [L_\alpha \cdot s] = [L_\alpha]_{tot} \qquad \text{Eq. S17}$$

Where $[L_\alpha]_{tot}$ is the total concentration of ligand $\alpha$ present.

Equations S8, S9, S10, S11, S13, S14, and S15 are implemented in the provided code. Note that this approach can be extended to multiple types of binding sites by introducing sums over different site types.

#### Synthetic Binding Data Generation

Unless otherwise specified, all synthetic binding data presented in the manuscript were generated using "*Model - Main Fitter.ipynb*", with a ligand size ($m$) of 1, a cooperativity parameter ($\omega$) of 1, an association constant ($K_a$) of 1, and a receptor concentration of 100.

Explicitly, the following parameters in site_params:

**Figure 3 (include_B=False)**

- One standard 1:1 binding site (equivalent to non-cooperative stacked binding): dict(n_A = 1, n_B = 1, res_tot=100, K_A=1, K_B=0, omega_AA=1, omega_AB=1, omega_BA=1, omega_BB=1)
- One cooperative stacked binding site: dict(n_A = 1, n_B = 1, res_tot=100, K_A=1, K_B=0, omega_AA=4, omega_AB=1, omega_BA=1, omega_BB=1)
- One linear binding site: dict(n_A = 4, n_B = 1, res_tot=100, K_A=1, K_B=0, omega_AA=1, omega_AB=1, omega_BA=1, omega_BB=1)
- One linear new-site binding site: dict(n_A = 4, n_B = 1, res_tot=100, K_A=1, omega_AA=1, K2=1) and new_site=True

**Figure 4 (include_B=True)**

- Two standard 1:1 binding sites (equivalent to non-cooperative stacked binding): dict(n_A = 1, n_B = 1, res_tot=100, K_A=10, K_B=1, omega_AA=1, omega_AB=1, omega_BA=1, omega_BB=1)
- One cooperative stacked and one linear binding site: dict(n_A = 4, n_B = 1, res_tot=100, K_A=1, K_B=1, omega_AA=1, omega_AB=1, omega_BA=1, omega_BB=4)
- One standard and one cooperative stacked binding site: dict(n_A = 1, n_B = 1, res_tot=100, K_A=1, K_B=1, omega_AA=4, omega_AB=1, omega_BA=1, omega_BB=1)
- One standard and one linear binding site: dict(n_A = 4, n_B = 1, res_tot=100, K_A=1, K_B=1, omega_AA=1, omega_AB=1, omega_BA=1, omega_BB=4)
- Two cooperative stacked binding sites: dict(n_A = 1, n_B = 1, res_tot=100, K_A=10, K_B=1, omega_AA=4, omega_AB=1, omega_BA=1, omega_BB=4)
- Two linear binding sites: dict(n_A = 4, n_B = 1, res_tot=100, K_A=10, K_B=1, omega_AA=1, omega_AB=1, omega_BA=1, omega_BB=1)

#### Additional Synthetic Binding Data

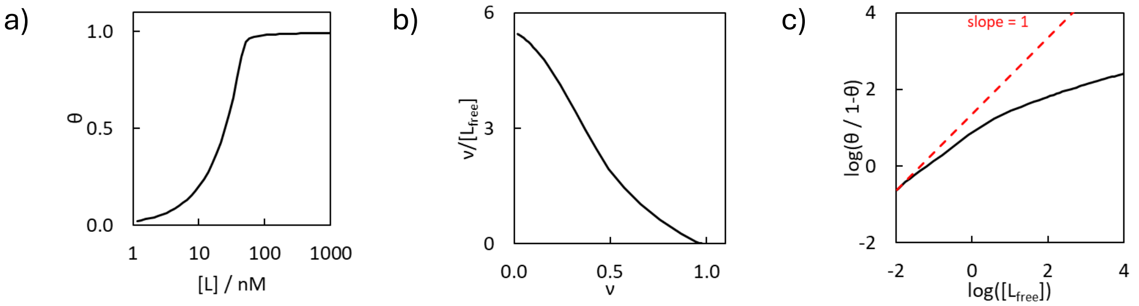


*Figure 18*. Representative (a) binding isotherm, (b) Scatchard plot, and (c) Hill plot for one ligand binding to two linear-cooperative sites ($K_1 = 1$, $K_2 = 10$, $m_1 = m_2 = 4$, $\omega_1 = \omega_2 = 4$).

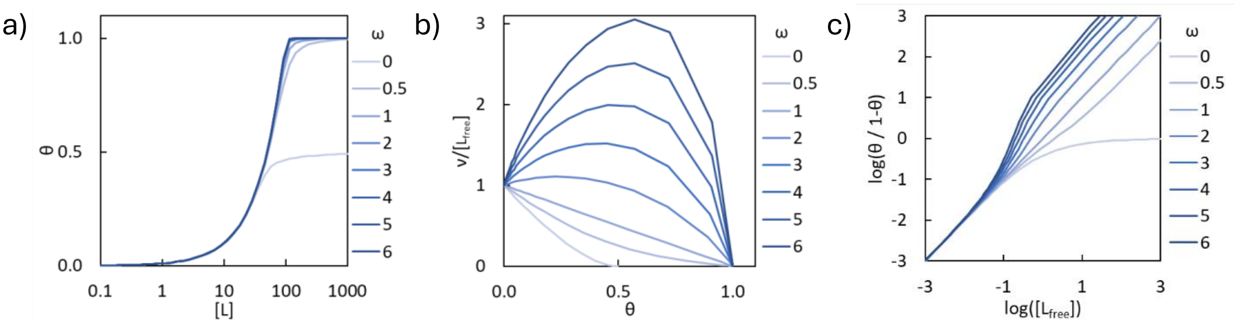


*Figure 19*. Representative (a) binding isotherm, (b) Scatchard plot, and (c) Hill plot for a ligand (m = 1) binding to a single site with different cooperativity parameter values.

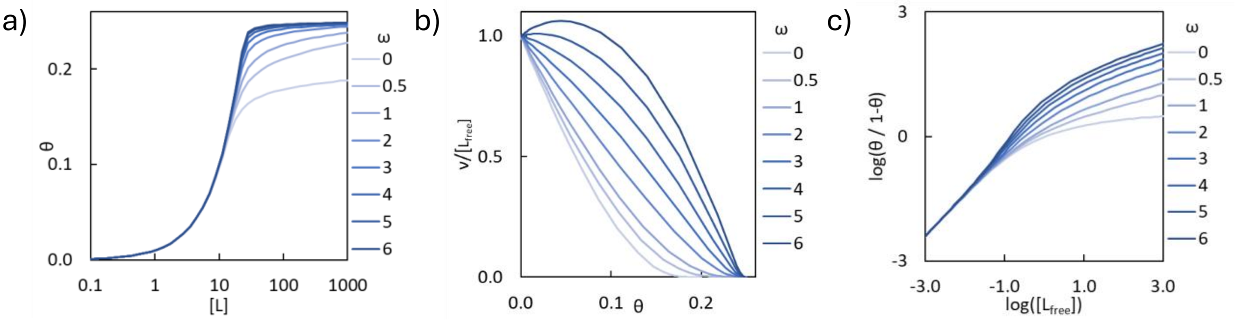


***Figure S20***. Representative (a) binding isotherm, (b) Scatchard plot, and (c) Hill plot for a ligand (m = 4) binding to a single site with different cooperativity parameter values.

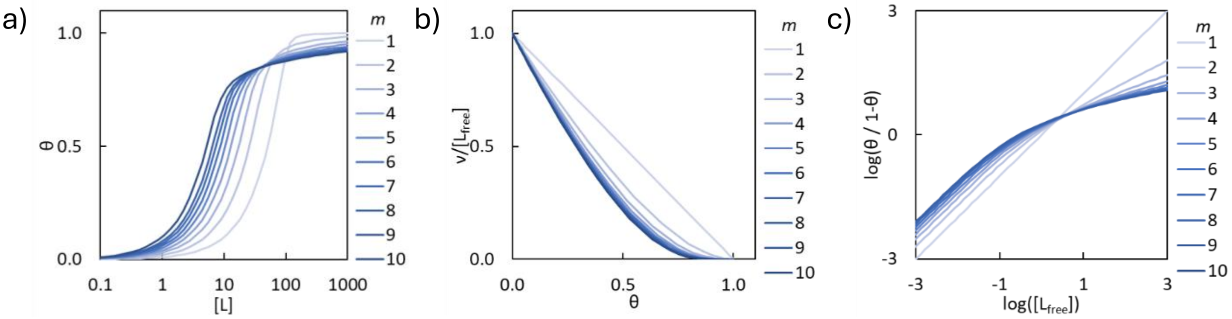


***Figure S21***. Representative (a) binding isotherm, (b) Scatchard plot, and (c) Hill plot for a non-cooperative ligand (ω = 1) binding to a single site with different ligand size values.

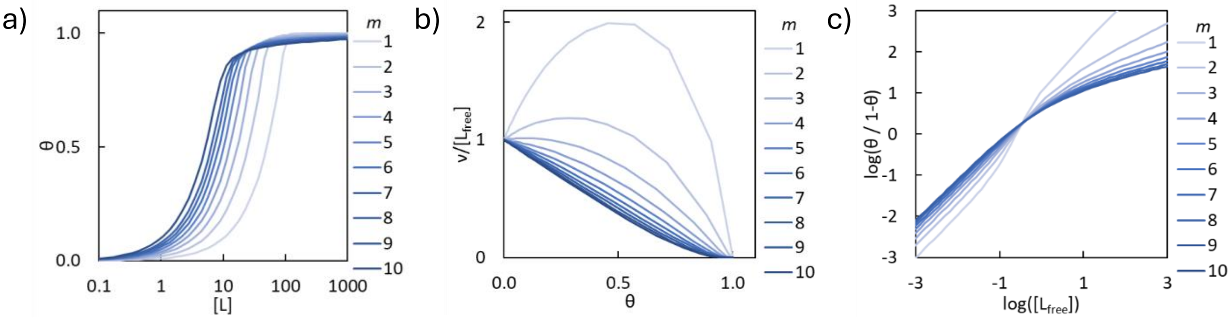


***Figure S22***. Representative (a) binding isotherm, (b) Scatchard plot, and (c) Hill plot for a cooperatively-binding ligand (ω = 4) binding to a single site with different ligand size values.

#### Radioligand Self-Displacement Assay Data

Data were extracted from a previously reported database of amyloid binding ligands.[6] Inclusion criteria were that a radioligand had to have a dissociation constant reported by both a saturation assay and a self-displacement assay, where identical structures were tested with the exception of radioisotopes, and that these data must be reported in a single publication. Assays must be against the same fibril target using a single set of buffer conditions at the same temperature. The data for the ligands used are in a supplementary "*Radioligand data.csv*" file provided.

#### Code for Isotherm Generation and Fitting.

An attached iPython notebook, "*Model - Main Fitter.ipynb*" is provided with code required to generate synthetic binding isotherms for the following models:

- One ligand binding to one or more types of sites
- Two ligands binding to one or more types of sites
- One ligand binding to one or more types of sites. Upon binding to a site, a secondary site is generated

An attached python file, "*speciation_1Dbinding.py*", is provided as a substitution for the "*speciation.py*" file incorporated in *Musketeer*. This code contains three new speciation classes to fit:

- Ligands binding to one type of site (SpeciationPolymer1D_1s)
- Ligands binding to two types of sites (SpeciationPolymer1D_2s)
- Ligands binding to one type of site, allowing for adjacent and non-adjacent ligands to have different fluorescence spectra (SpeciationPolymer1D_1s_adjacency)

A specific input format is required when entering concentrations in the Musketeer speciation table: enter columns of host concentrations first, followed by columns of guest concentrations.

Code was written with assistance from ChatGPT 5.2.

#### Comparing Fits

When fitting raw data, models were compared using either RMSE values generated by *Musketeer*,[7] or the Akaike Information Criterion corrected for small sample sizes (AICc). When using AICc the effective sample size was taken as the number of additions in the binding assay, each addition corresponding to a fluorescence emission spectrum obtained at a specific ligand concentration. The data fitting procedure generates a fitted optical brightness for the bound species at each wavelength; comparable fits were obtained when fitting only a single wavelength, and so each brightness spectrum was treated as a single effective parameter.

#### Symbols Used in Equations

| | |
|---|---|
| $[r_{free}]$ | The concentration of free lattice residues |
| $[s]_t$ | Total concentration of binding sites |
| $[s]$ | Concentration of free binding sites |
| $[L_\alpha]$ | Free concentration of a ligand of type $\alpha$ |
| $[L_\alpha]_{tot}$ | Total concentration of a ligand of type $\alpha$ |
| $[L_\alpha \cdot s]$ | Concentration of a ligand of type $\alpha$ bound to site $s$ |
| $f$ | A free lattice residue |
| $b^i_{m_i}$ | A lattice residue bound to the $m_i$th segment of a ligand of type $i$ |
| $K_\alpha$ | The equilibrium binding constant for ligand $\alpha$ |
| $\omega_{ij}$ | The cooperativity parameter between a ligand of type $i$ and a ligand of type $j$ |
| $n_H$ | The Hill coefficient |
| $m_i$ | Size of ligand of type $i$ in lattice units |
| $\nu_i$ | The average number of ligands of type $i$ bound per unit lattice |
| $\theta_i$ | Coverage for ligand of type $i$ |
| $\theta$ | Total coverage of the lattice |
| $n_i$ | The number of ligands of type $i$ bound |
| $N$ | Total number of lattice units |
| $T$ | Total number of types of ligands |
| $g$ | The size of a gap in lattice units |
| $(P_g)_{ij}$ | The probability that a gap between a ligand of type $i$ and a ligand of type $j$ is $g$ lattice residues long |
| $(P_g)$ | The probability that a gap being $g$ lattice residues long |
| $\bar{s}_\alpha$ | The average number of free binding sites per gap for ligand $\alpha$ |
| $(\bar{s}_{iso})^\alpha_{ij}$ | The average number of isolated binding sites for ligand $\alpha$ in a gap flanked by a ligand of type $i$ on the left and a ligand of type $j$ on the right |
| $(\bar{s}_{sc})^\alpha_{ij}$ | The average number of singly contiguous sites for ligand $\alpha$ in a gap flanked by a ligand of type $i$ on the left and a ligand of type $j$ on the right |
| $(\bar{s}_{dc})^\alpha_{ij}$ | The average number of doubly contiguous binding sites for ligand $\alpha$ in a gap flanked by a ligand of type $i$ on the left and a ligand of type $j$ on the right |
| $(\bar{s}_{iso})^\alpha$ | The average number of isolated binding sites for ligand $\alpha$ in a gap |
| $(\bar{s}_{sc})^\alpha$ | The average number of singly contiguous binding sites for ligand $\alpha$ in a gap |
| $(\bar{s}_{dc})^\alpha$ | The average number of doubly contiguous binding sites for ligand $\alpha$ in a gap |
